# Anoctamin 10/TMEM16K mediates convergent extension and tubulogenesis during notochord formation in the early chordate *Ciona intestinalis*

**DOI:** 10.1101/2023.01.20.524945

**Authors:** Zonglai Liang, Daniel Christiaan Dondorp, Marios Chatzigeorgiou

## Abstract

During embryonic development, cells are organized into complex tissues and organs. A highly conserved organ shape across metazoans is the epithelial tube. Tube morphogenesis is a complex multistep process where the molecular mechanisms underlying the diversity of cell behaviors such as convergent extension, cell elongation, and lumen formation are still intensely studied. Here, using genome editing and quantitative imaging in the early chordate *Ciona intestinalis* we show that Ano10/Tmem16k, a member of the evolutionarily ancient family of transmembrane proteins called Anoctamin/TMEM16 is required for convergent extension, lumen expansion and connection during notochord morphogenesis. In addition, we find that loss of Cii.Ano10/Tmem16k hampers cell behavior and cytoskeletal organization during tubulogenesis. In vivo Ca^2+^ imaging revealed that genetic ablation of Cii.Ano10/Tmem16k hinders the ability of notochord cells to regulate bioelectrical signaling. Finally, we use electrophysiological recordings and a scramblase assay in tissue culture to demonstrate that Cii.Ano10/Tmem16k likely acts as an ion channel but not as a phospholipid scramblase. More generally, this work provides insights into the pre-vertebrate functions of Anoctamins and raises the possibility that Anoctamin/Tmem16 family members have an evolutionarily conserved role in tube morphogenesis.

## Introduction

Biological tubes play an essential role in embryogenesis, organogenesis, and post-embryonic physiology^1,2^. Indeed, tubular structures are widespread across vertebrates and invertebrates, including salivary glands, renal tubules, vasculature, intestinal tract, and bronchial tubules. Reliable formation of these diverse tubular organs depends on the highly coordinated interplay between collective cell behaviors and complex signaling processes ^1,3-5^. Errors in tube formation during development or malfunctions in adult tubular structures can result in severe pathologies^2,6^.

The notochord is an essential tubular organ present in the embryonic midline region of all members of the chordate phylum^7^. Formation of this flexible rod, tapered at both ends ^8^, is fundamental to providing structural support to the developing chordate embryo. In the case of vertebrates, the notochord further serves as a signaling center that secretes factors to pattern surrounding tissues ^9^. Ascidians, which belong to the sister group to vertebrates the tunicates, have a notochord composed of only 40 cells^10-12^. The small cell number makes it an ideally tractable model for dissecting the rudimentary cellular and molecular mechanisms underlying the multistep process of notochord formation. For example, in *Ciona intestinalis*, notochord development begins with cell intercalation-driven convergent extension^13-15^. Next, the notochord elongates to form a cylindrical rod via actomyosin network dependent-cell shape changes^16,17^. Fundamental to tube formation is the establishment of lumens. Here, the notochord cells of *Ciona* undergo a mesenchymal-to-epithelial transition (MET), that leads to the emergence of apical domains at the opposite ends of each cell^10,18^. Extracellular lumens thereby appear and then expand between neighboring cells of the developing tube. Finally, the notochord cells crawl bi-directionally, exhibiting an endothelial-like cell morphology, which leads to merging of the lumens^10,18^.

Tubulogenesis has been intensively studied across a diversity of models, such as the salivary gland of *Drosophila*, the excretory cells of *Caenorhabditis elegans*, the lungs, mammary glands and neural tube of mice and the notochord of zebrafish and *Ciona intestinalis* ^3,19-21^. Transmembrane proteins have since emerged as an important regulator of signal transduction during the process ^19,22-26^. More generally transmembrane proteins such as ion channels and transporters are important in signaling during development ^26-34^.

To further focus on how bioelectrical signaling shapes the developing tubular organ, we turned to transmembrane pumps, transporters and ion channels. Of these, the role of the widely conserved Anoctamin (Ano/Tmem16) family proteins ^35,36^ is essentially unexplored in developmental tube morphogenesis even though several of the Anoctamins have previously been shown to be expressed in mammalian tubular structures including pancreatic acinar cells, proximal renal tubules and several glands (e.g., submandibular glands, Leydig cells) ^37-43^.

Anoctamins generally function as Ca^2+^-activated Cl^-^ channels (CaCCs) and phospholipid scramblases (PS); the latter capable of translocating phospholipids between the two monolayers of a membrane ^37,38,44^. They have been reported to take part in diverse cellular functions, including for example signal transduction, cell migration, Cl^-^ secretion, and volume regulation ^38,40,45,46^. While some Anoctamins, such as ANO1/TMEM16A, act as a CaCC^37,47,48^, and ANO6/TMEM16F, as a PS^44^; for other members of the family, including ANO10/TMEM16K, their functions remain under fruitful debate^38,49,50^.

Here we leverage the simplicity and genetic accessibility of the *C. intestinalis* notochord to study the role of *C*.*intestinalis* Ano10/Tmem16k (gene model: KH.C3.109/KY21.Chr3.1036 from here on referred to as Ano10) in tube morphogenesis. We employ tissue-specific CRISPR/Cas9 knockout and rescue experiments to reveal that Ano10 is required for convergent extension, lumen expansion and lumen connection. At the level of cell behavior, Ano10 modulates actin and microtubule organization affecting cell motility and shape. At the level of signaling, Ano1*0* regulates Ca^2+^ activity during convergent extension and tubulogenesis. We further demonstrate that Ano10 acts not as a phospholipid scramblase but instead as an ion channel, where its role in establishing and/or maintaining an appropriate electrochemical balance across cell membranes is likely essential for notochord morphogenesis.

## Results

### Ano10 is expressed in the notochord during embryonic development and localizes primarily to the ER

In addition to the previously identified three Anoctamins in *C. intestinalis* and *C. savignyi* ^*36*^, this work has yielded an additional Anoctamin for *Ciona intestinalis*. To explore the homology of *C. intestinalis* Anoctamins, we constructed a phylogenetic tree with 89 known Anoctamins from various organisms. Of these, we show that Ciinte.CG.KH.C3.109/ KY21.Chr3.1036 clusters together with vertebrate ANO10/TMEM16K proteins (Figure S1A). Protein alignment analysis additionally revealed that the ten transmembrane domains and the putative calcium-binding sites of Ano10 share strong conservation with vertebrate ANO10/TMEM16K (Figure S1B).

To determine the expression pattern of *Ano10*, we employed in situ hybridization (ISH) and a transcriptional reporter. Using the first approach, we found that in neurula and initial tailbud stages, *Ano10* mRNA expression was strongest in the mesenchyme and the notochord (Figure S2A-D). At later tailbud stages, the expression became localized to the notochord (Figure S2E-K). Second, we generated a transcriptional fusion containing a 2kb region upstream of the *Ano10* start site that drove expression of GFP in the notochord during mid and late tailbud stages (Figure S2L,M). Thus, both approaches suggest that *Ano10* is expressed in the developing notochord.

Next, to determine the subcellular localization of Ano10, we co-expressed Ano10-GFP and KDEL::BFP in a notochord-specific manner under the Brachyury promoter, which revealed that at the end of cell elongation (Stage IV according to Dong et al.^18^) Ano10 is primarily localized to the Endoplasmic Reticulum (ER) (Figure S2 N-P) However, in embryos imaged during lumen connection (Stage VII according to Dong et al. ^18^), Ano10 localized to the plasma membrane in addition to the ER (Figure S2Q-R). Furthermore, we found that in human cells, Ano10::GFP colocalizes with mCherry-Sec61β, an established ER marker ^51^ (Figure S3A-C). A small amount of Ano10 was found additionally on the plasma membrane (Figure S3D-G). Combined, our whole-animal and tissue culture data indicate that Ano10 is primarily localized to the ER, with only a small proportion of the protein reaching cell surface.

### *Ano10* is required for notochord convergent extension

Having established that *Ano10* is expressed in the notochord during embryogenesis, we hypothesized that it may contribute to notochord development. To test this, we conducted tissue-specific loss-of-function analysis using the CRISPR/Cas9 system to target *Ano10* exclusively in the notochord^52^. Confocal microscopy imaging using transgenes labelling the plasma membrane and nuclei of notochord cells showed that in negative control embryos notochord development appeared normal both during the process of convergent extension (Figure 1A) as well as during the elongation period resulting in a single column of 40 stacked cells and full tail extension (Figure 1D). In contrast, *Ano10*^CRISPR^ embryos were characterized by irregular convergent extension at initial tailbud stages (Figure 1B), while at late tailbud stages the embryos exhibited defective Anterior-Posterior (A/P) elongation and abnormal notochord morphology with local thickenings; or multiple columns containing groups of notochord cells that failed to properly intercalate (Figure 1E). We were able to restore notochord morphology by tissue-specific rescue, where we co-electroporated together with the *Ano10* targeting gRNA and the Cas9 mix, a notochord-specific rescue plasmid encoding *Ano10* cDNA with 5 nucleotides replaced (without changing the amino acids encoded) to avoid being targeted by the gRNA (Figures 1C,F).

**Figure 1.**
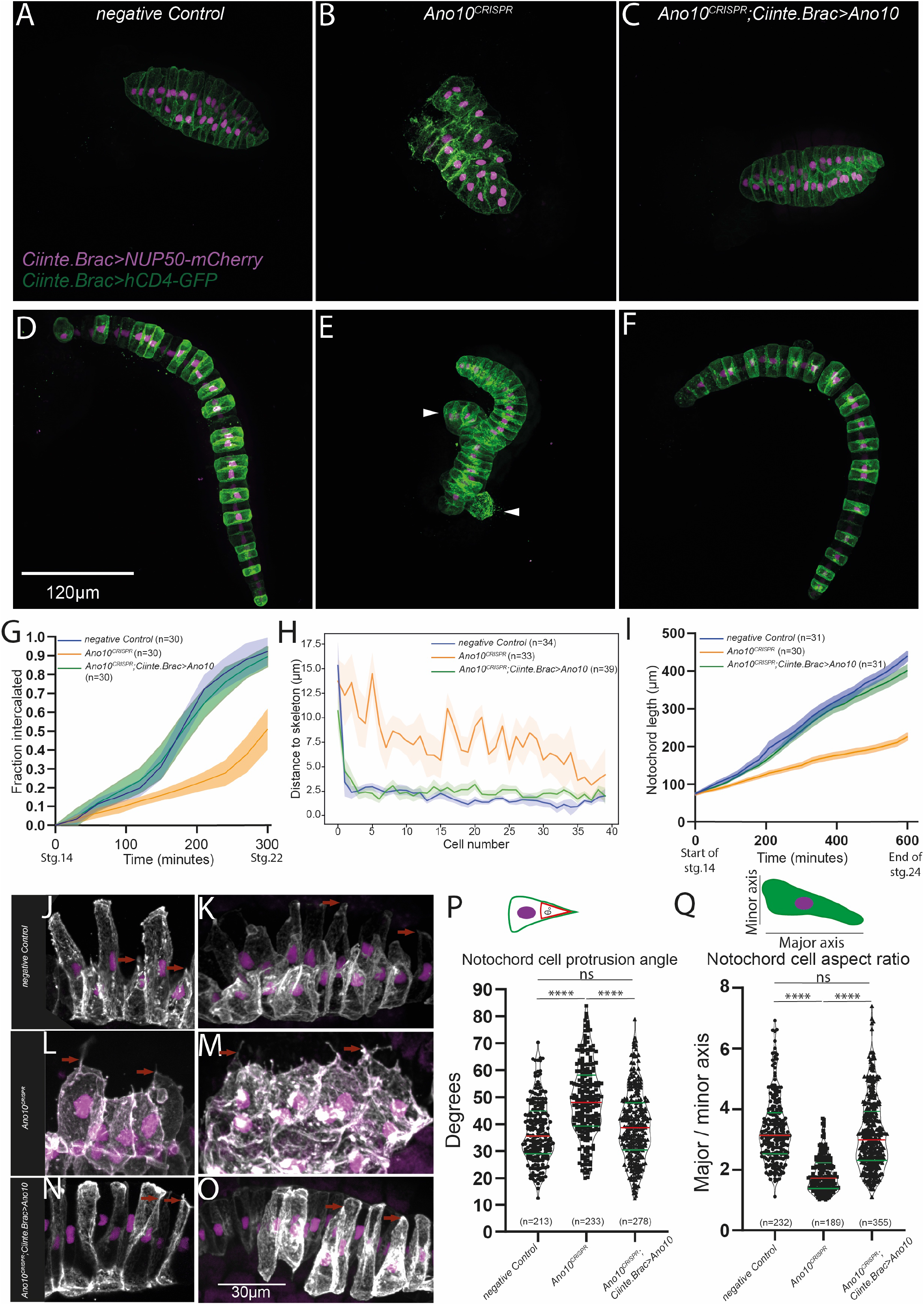
*Ano10* is required for notochord convergent extension. (A-C) Maximal projections of confocal stacks showing stage 17^79^ embryos where the notochord cells undergoing intercalation are labelled using *Ciinte*.*Brac>hCD4GFP* and *Ciinte*.*Brac>NUP50-mCherry* (A) negative control, (B) *Ano10*^*CRISPR*^ and (C) rescue *Ano10*^*CRISPR*^;*Ciinte*.*Brac>Ano10*. (D-F) Maximal projections of stage 23^79^ embryos after the completion of notochord cell intercalation. White arrowheads indicate misaligned cells. (G) Fraction of notochord cells with completed intercalation at a given time point starting from stage 14 and ending at stage 22^79^. Data presented as mean ± SEM, 30 animals analysed for each genotype. For statistical analysis we used a 2-way ANOVA, followed by a Tukey’s multiple comparisons test. (see also Supplementary Table 1). (H) Notochord cell misalignment quantification measured as the distance from notochord cell centroid to the embryo midline skeleton in late tailbud embryos as a function of AP position from cell 0 (anterior) to cell 39 (posterior) for the negative control, *Ano10*^*CRISPR*^ and *Ano10*^*CRISPR*^;*Ciinte*.*Brac>Ano10* rescue embryos. Data presented as mean ± SEM, n= number of animals analysed. For statistical analysis we performed Mann-Whitney test (see also Supplementary Table 1). (I) Change in overall notochord length increase during development. Data presented as mean ± SEM, n= number of animals analysed. We used a mixed-effects model (REML), followed by Tukey’s multiple comparisons test for pairwise comparisons (see also Supplementary Table 1). (J-O) Maximal projections of different genetic background embryos expressing mosaically *Ciinte*.*Brac>hCD4GFP* and *Ciinte*.*Brac>NUP50-mCherry* during initial tailbud I stage. Examples of medial protrusions are highlighted with red arrows. (P, Q) Quantification of notochord cell medial protrusion angles and cell aspect ratios. Red line in violin plots corresponds to the median and green lines indicate quartiles (n=number of cells analyzed). For statistical analysis we performed Kruskal-Wallis test, followed by Dunn’s multiple comparisons test (see also Supplementary Table 1 for p-value details).

To obtain a quantitative measure of the intercalation defects in *Ano10*^CRISPR^ embryos we measured the fraction of notochord cells that have undergone mediolateral intercalation (ML) during notochord development. We find that *Ano10*^CRISPR^ embryos exhibited a significantly lower fraction of successfully intercalated cells over time compared to negative control and rescue embryos (Figure 1G). At the end of ML intercalation in negative control and rescue embryos almost all cells had successfully intercalated, in contrast to *Ano10*^CRISPR^ embryos where on average less than 50% of the notochord cells completed intercalation (Figure 1G). A second approach we took to quantify defects in notochord morphology of *Ano10*^CRISPR^ embryos due to erroneous intercalation was to measure the distance from the notochord cell center to the midline skeleton. In control embryos where the notochord had successfully completed the intercalation process forming a 40-cell long stack of cells centered along the embryo midline, the distance of each cell center relative to the midline of the skeleton was very short (Figure 1H). In contrast, most notochord cells in the *Ano10*^CRISPR^ embryos were positioned significantly further away from the midline skeleton (Figure 1H). Notochord-specific rescue of the *Ano10*^CRISPR^ defect suggests that *Ano10* acts in a cell autonomous manner to regulate notochord cell intercalation (Figure 1H).

**Table 1.**
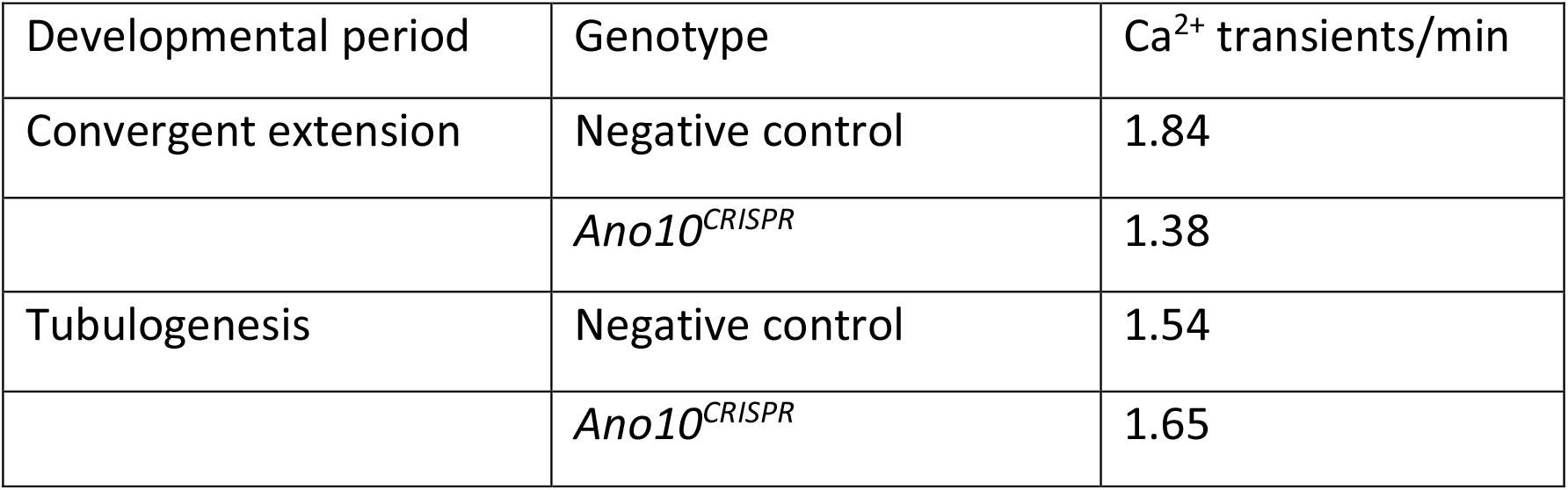
Ca^2+^ transient frequency in negative control and *Ano10*^*CRISPR*^ notochord cells during convergent extension and tubulogenesis.

Following the termination of intercalation, notochord cells will elongate along the A/P axis, taking a barrel-like shape^10,18^. As a result of this process the notochord length progressively increases. We measured the evolution of overall notochord length of the notochord from the point that the sheet of notochord cells invaginates to form a rod of cells until the end of the elongation process. We found that in negative control embryos notochord development proceeded normally resulting in full tail extension (Figure 1I). In contrast, in *Ano10*^CRISPR^ embryos notochord length increased at a slower rate, reaching a shorter final length (Figure 1I) compared to both negative control and rescue embryos (Figure 1I). We then wondered whether the mediolaterally polarized protrusive activity of notochord cells during cell intercalation was perturbed by the genetic ablation of *Ano10*. In contrast to negative control and rescued notochord cells (Figure 1J,K; Figure 1N,O), in *Ano10*^CRISPR^ cells these protrusions appeared shorter and often decorated with multiple filopodia-like structures (Figure 1L,M). In addition, we found that the median value of the angle enclosed by the notochord cell protrusion in Ano*10*^CRISPR^ cells was significantly larger compared to negative control and rescued cells (Figure 1P). Conversely, the median aspect ratio of *Ano10*^CRISPR^ cells was significantly smaller (Figure 1Q). These morphometric parameters indicate that *Ano10* is required for the maintenance of an elongated cell shape during cell intercalation.

The observed cell intercalation phenotypes could also be potentially attributed to a defect in mediolateral polarization or the loss of the notochord boundary. Visual inspection of confocal stacks indicated that *Ano10*^CRISPR^ notochord cells exhibited protrusions with an overall mediolateral bias, while the nuclei exhibited a medial localization. Taken together with the observation that cell protrusions had a mediolateral bias, it would suggest that Ano10 is not required for mediolateral polarization. Analysis of the border of *Ano10*^CRISPR^ notochords (Figure S4B) showed reduced regularity and definition compared to negative control notochords (Figure S4A). To quantify this border defect, we calculated the ratio of total/net boundary length(Figure S4C-I). In *Ano10*^CRISPR^ embryos this ratio was significantly higher compared to negative controls (Figure S4I). Rescued embryos had a fully restored total/net boundary length ratio (Figure S4I). Importantly, we detected laminin by immunostaining at the notochord boundary of both negative control and *Ano10*^CRISPR^ albeit with weaker staining (Figure S4J-L), suggesting that the notochord boundary is present, but deformed, in *Ano10*^CRISPR^. Finally, we note that the defects in intercalation of *Ano10*^CRISPR^ notochord cells did not emerge from spurious cell proliferation since both negative control and *Ano10*^CRISPR^ embryos had a median of 40 cells per notochord (Figure S4M).

### Ano10 is required for notochord tubulogenesis

We then sought to investigate whether Ano10 contributes to the process of tubulogenesis. While *Ano10*^CRISPR^ embryos generated using the *Ciinte*.*Brachyury* promoter (driving Cas9) exhibited strong tubulogenesis phenotypes, we were concerned that these defects could be the indirect outcome of erroneous cell intercalation. To overcome this issue we drove expression of Cas9 using the promoter of carbonic anhydrase (Fig S5A-F) which is expressed in the notochord at later developmental stages^53^ compared to Brachyury^54,55^, thus allowing us to assess the role of Ano10 specifically during tubulogenesis. To obtain independent validation of the genome editing based findings we used an acute pharmacological perturbation by applying three different drugs that are well established blockers of Anoctamins ^56-58^ just prior to the onset of lumen initiation (at the end of stage IV^18^). To quantitatively assess notochord cell behavior during lumen formation, we generated and imaged using time-lapse confocal microscopy transgenic embryos co-expressing the transmembrane proteins CD4::GFP and *Ciiinte*.Caveolin 1-mCherry ^23^ to simultaneously visualize the plasma membrane and the apical/luminal region of the notochord cells. Following cell elongation in negative control embryos (Figure 2A, asterisks) or DMSO treated embryos (Figure S6A, asterisks), apical domains form in the center of the lateral domains as previously shown ^23^. These are also present in *Ano10*^CRISPR^ notochord cells (Figure 2B, asterisks) indicating that Ano10 is not involved in cell polarization and lumen initiation. We also observed apical domains in drug treated animals (Figure S6 B-D). In addition, when we measured the longitudinal radius of the lumen at the end of the initiation period the median value for *Ano10*^CRISPR^ cells was indistinguishable from that of negative controls (Figure 2D, E).

**Figure 2.**
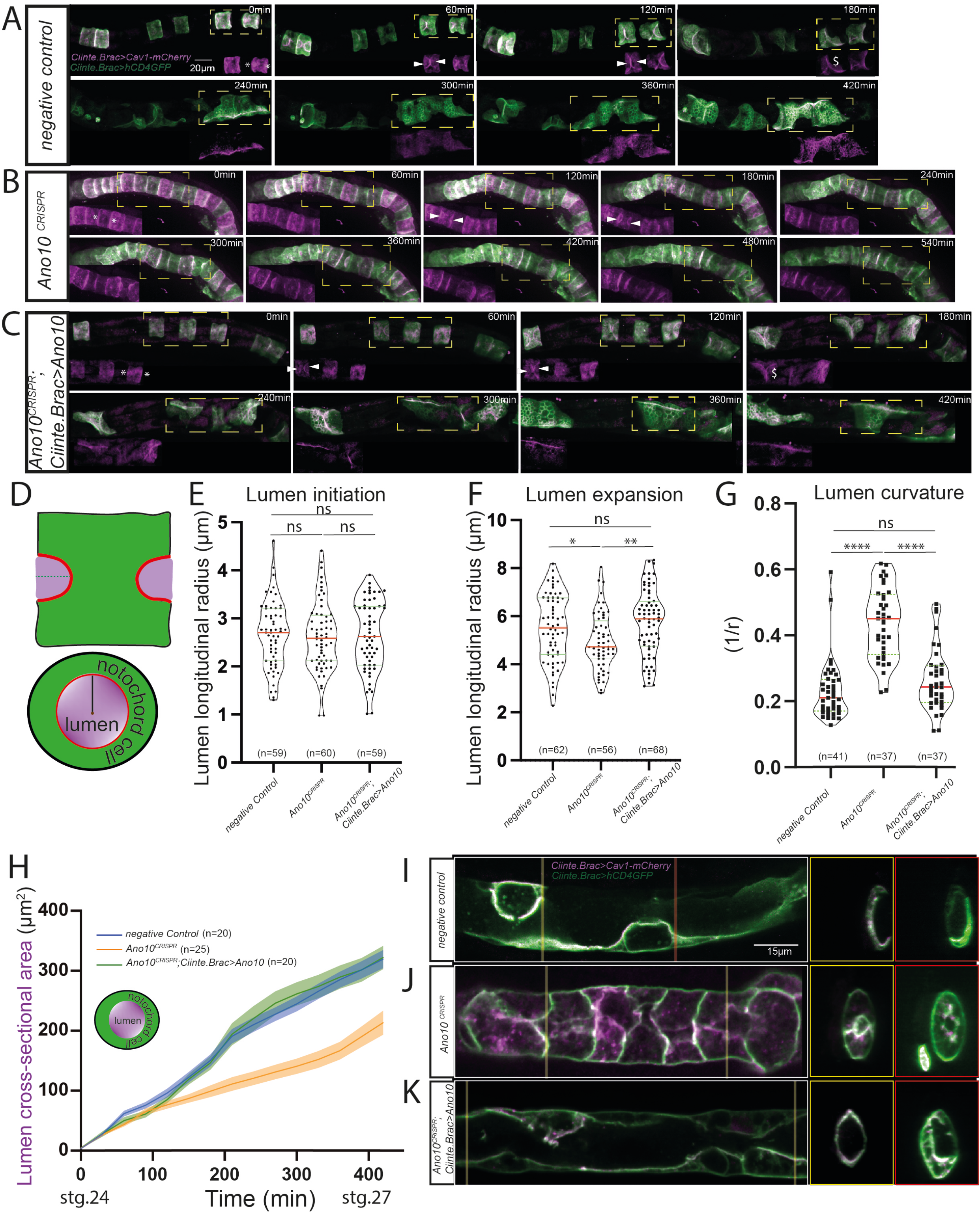
Ano10 is required for notochord tubulogenesis. (A-C) Montage of maximum projections showing confocal time lapse imaging of *Ciiinte*.*Brac>hCD4-GFP; Ciiinte*.*Caveolin 1-mCherry* covering the period of tubulogenesis. Insets show *Ciiinte*.Caveolin 1-mCherry signal from the dashed orange boxes. Asterisks indicate the emerging apical domains. White arrowheads point to the expanding lumen (see also Supplementary Movies 1-3) (D)Top schematic illustrates a longitudinal view of the notochord cell during lumen extension. The red lines define the apical domain and the dashed green line the longitudinal radius. The purple area defines the expanding lumen volume. Lower schematic illustrates a transverse section through the same cell. Black line indicates radius of curvature and the brown point the center of curvature. (E) Quantification of lumen longitudinal radius during lumen initiation. (F) Quantification of lumen longitudinal radius during lumen expansion. (G) Quantification of lumen curvature after lumen connection. For statistical analysis of data shown in panels E-G we performed Kruskal-Wallis test followed by Dunn’s multiple comparisons test, (see also Supplementary Table 2 for p-value details). In parentheses we indicate the number of animals used in our analysis. (H) Evolution of lumen cross-sectional area during tubulogenesis. Schematic shows in purple the lumen cross-section and in green the notochord cell membrane. For statistical analysis we performed a 2-way ANOVA followed by Tukey’s multiple comparison test (see also Supplementary Table 2). In parentheses we indicate the number of animals used in our analysis. (I-K) Longitudinal cross-sections of example notochords from negative control (I), *Ano10*^CRISPR^ (J) and rescue (K) embryos. Transverse sections from two points along the notochord. The most anterior is indicated by a yellow bar and the more posterior section is highlighted with a red bar.

However, *Ano10*^CRISPR^ notochord cells displayed a limited ability to expand their lumen (Figure 2B, white arrowheads). We found that the lumen longitudinal radius at the end of the expansion phase was significantly shorter in *Ano10*^CRISPR^ compared to negative control and rescued embryos (Figure 2F). Additionally, lumen curvature measured from transverse notochord sections was significantly higher in *Ano10*^CRISPR^ cells suggesting that they had a narrower lumen compared to negative control and rescued cells (Figure 2D, G). Drug treated notochord cells exhibited a similar behavior in terms of lumen initiation (Figure S6E), expansion (Figure S6B-D white arrows, S6F) and curvature (Figure S6G) as *Ano10*^CRISPR^ notochord cells. Taken together these findings suggest that Ano10 is not required for lumen initiation, but it is necessary for lumen expansion and lumen geometry. Following lumen expansion negative control notochord cells initiate the process of tilting and connecting the notochord lumen (Figure 2A, C see $ sign). We observed that *Ano10*^CRISPR^ notochord cells were not able to tilt and connect the notochord lumen to the same extent as negative control and rescue cells (Figure 2B). To analyze the dynamics of lumen growth we measured the cross-sectional area change from lumen initiation until the completion of lumen connection (Figure 2H). We found that *Ano10*^CRISPR^ notochord lumens showed a consistently narrower cross-sectional area compared to negative controls and that this defect could be rescued in a tissue specific manner (Figure2 H-K). Drug-treated notochords exhibited a lumen cross-sectional area progression phenotype that was very close to that of *Ano10*^CRISPR^ notochords(Figure S6H). Thus, we believe that Ano10 is required for expanding, tilting and connecting the notochord lumen.

**Table 2.** Scramblase activity of mAno6, C. intestinalis Ano10 and Ano6

| Condition | Total number of cells | AnnexinV (+) | Percentage (%) |
| --- | --- | --- | --- |
| Negative control (DMSO) | 1049 | 16 | 1.53 |
| Negative control (Ionomycin) | 1135 | 21 | 1.85 |
| mAno6 (DMSO) | 1247 | 32 | 2.57 |
| mAno6 (Ionomycin) | 1129 | 185 | 16.39 |
| Cii.Ano10 (DMSO) | 1014 | 11 | 1.08 |
| Cii.Ano10 (Ionomycin) | 905 | 4 | 0.44 |
| Cii.Ano10-CAAX (DMSO) | 849 | 13 | 1.53 |
| Cii.Ano10-CAAX (Ionomycin) | 1021 | 27 | 2.63 |
| Cii.Ano6 (DMSO) | 1182 | 38 | 3.2 |
| Cii.Ano6 (Ionomycin) | 1333 | 408 | 30.61 |

### Ano10 regulates cell behavior and cytoskeletal organization during tubulogenesis

We wondered whether the defect in lumen tilting, and connection exhibited by *Ano10*^CRISPR^ notochords could be attributed to perturbed morphology and locomotion of notochord cells. A characteristic morphological transition associated with the crawling behavior that notochord cells exhibit during lumen expansion and tilting is the development of protruding anterior or posterior leading edges (ALE, PLE) composed of lamelopodia (LA) and new extensions (NE) (Figure 3A,B)^18^. We observed that in confocal micrographs *Ano10*^CRISPR^ notochord cells were rounder, characterized by truncated leading edges that were often missing lamellipodia and had smaller new extensions (NE) (Figure 3C,D). This phenotype could be rescued in a notochord-specific manner (Figure 3E,F). Quantification of the length of the ALE region across negative controls, *Ano10*^CRISPR^ and rescue notochord cells showed that the ALE in *Ano10*^CRISPR^ cells was significantly shorter than negative controls (Figure 3G). We also quantified the aspect ratio of these cells and found that *Ano10*^CRISPR^ cells had a significantly lower aspect ratio (Figure 3H), indicating that they could not switch from the cylindrical shape they had at the onset of tubulogenesis, to the elongated endothelial-like flat cells. Interestingly, we found that these morphological defects were associated with the markedly slower movement of *Ano10*^CRISPR^ cells during tubulogenesis (Figure 3I). The successful notochord-specific rescue of the *Ano10*^CRISPR^ defects suggests that Ano10 is required in a cell autonomous manner to mediate several cell behaviors occurring during the lumen tilting and connection.

**Figure 3.**
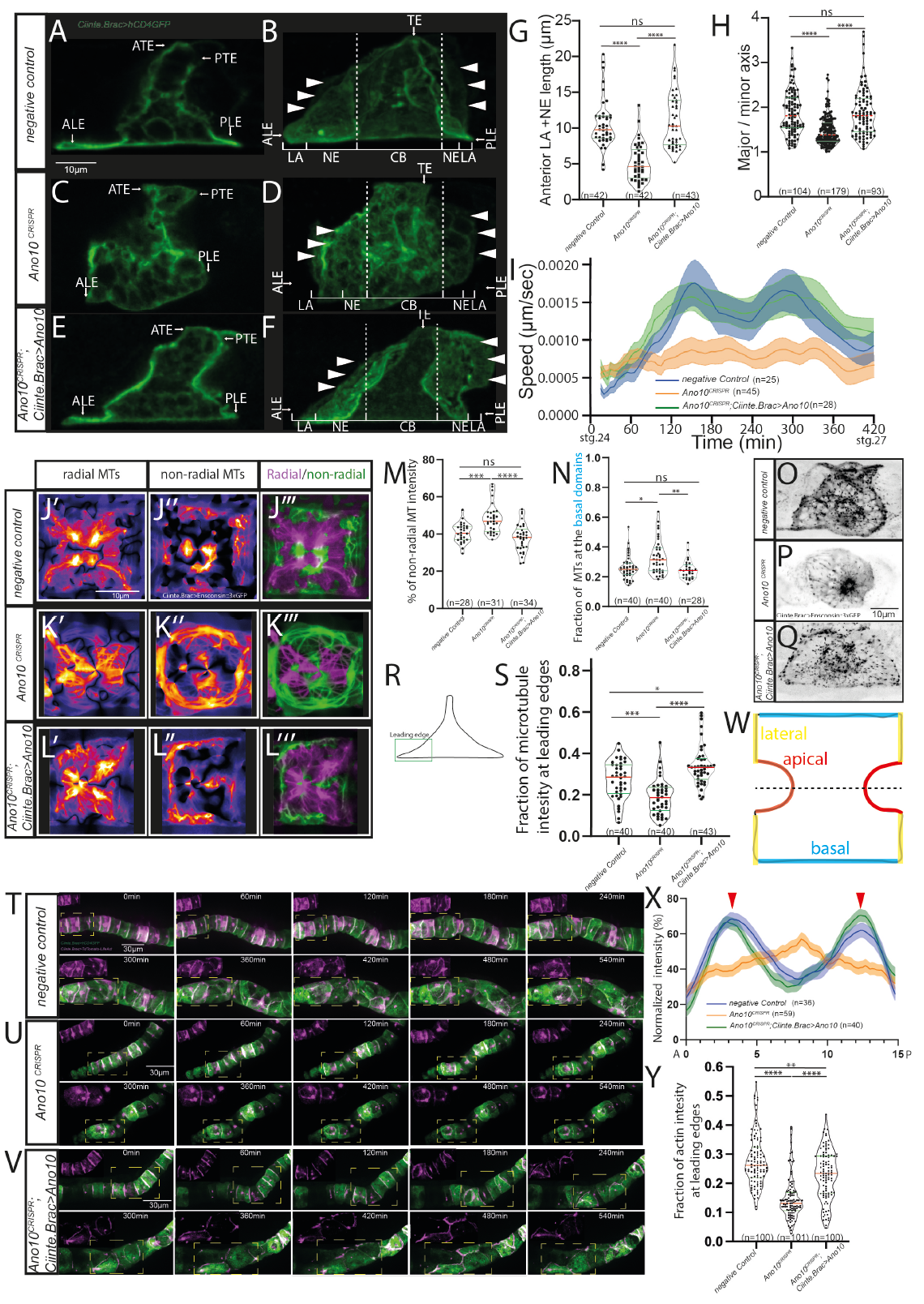
Loss of Ano10 perturbs cell morphology, behavior and the cytoskeletal dynamics during tubulogenesis. (A-F) Confocal images of notochord cells expressing plasma membrane targeted GFP. A,C,E show medial sections while B, D, F show maximal projections of the same cells. In stage VI, the Anterior Trailing Edge (ATE) and the Posterior Trailing Edge (PTE) retreat along the A/P axis, in contrast to the Anterior Leading Edge (ALE) or the Posterior Leading Edge (PLE) which extend along the A/P axis. Landmarks in the cells were determined in accordance with Dong et al^18^. The dashed lines in B, D, F contain the space occupied by the cells prior to the formation of protrusions. White arrowheads show the edges of the lamellipodia, which have wavy outlines. LA, lamellipodia; NE, new cell extension; TE, trailing edge; CB, main cell body. (G) Quantification of Anterior cell protrusion as defined by the combined length of the LA and NE regions. In parentheses we indicate the number of cells used in our analysis. (H) Quantification of the aspect ratio of notochord cells at stage VI. In parentheses we indicate the number of cells used in our analysis. (I) Evolution of apical domain speed from lumen initiation until the lumen connection is complete. Data presented as mean ± SEM. (n=cells used in analysis). For panel I we used a Mixed-effects model (REML) followed by Tukey’s multiple comparisons test. (J-L) Single notochord cells expressing *Ciinte*.*Brac> Ensconsin::3xGFP* where radial (J’,K’,L’),non-radial (J’’,K’’,L’’) MTs are highlighted using the FIJI plugin Radiality Map. A merge of non-radial and radial MTs is shown in (J’’’,K’’’,L’’’). (M) Quantification of non-radial MT intensity (n=cells used in analysis). (N) Quantification of the MTs localized at the basal domains of the notochord cells (n=cells used in analysis). (O-Q) Confocal maximal projection of a notochord cell during crawling from (O) negative control, (P) *Ano10*^CRISPR^ and rescue (Q) embryos expressing *Ciinte*.*Brac>EB3::mNeonGreen*, which allows us to visualize the microtubule plus ends. (R) The schematic illustrates a crawling notochord cell. The leading-edge area is highlighted by a green box. (S) Quantification of the fraction of microtubule intensity at the leading edges, (n=cells used in analysis). (T-V) Montage of notochords from (T) negative control, (U) *Ano10*^CRISPR^ and (V) rescue embryos expressing *Ciinte*.*Brac>hCD4GFP; Ciinte*.*Brac>LifeAct:: tdTomato* during tubulogenesis. Insets show the signal from LifeAct::tdTomato corresponding the region highlighted by the yellow dashed lines. (W) Schematic of a notochord cell during lumen extension. We highlight in red the apical domains, in yellow the lateral domains and in blue the basal domains. (X) Normalized actin profile across notochord cells (corresponding to the dashed black line in W) imaged during lumen extension. Red arrowheads indicate the points that correspond to the apical domains. Data presented as mean ± SEM. For panel I we used a Mixed-effects model (REML) followed by Tukey’s multiple comparisons test (See also Supplementary Table 3). (Y) Quantification of the fraction of actin intensity at the leading edges, (n=cells used in analysis) in notochord cells expressing *Ciinte*.*Brac>LifeAct::tdTomato* in different genetic backgrounds. In all violin plots in this figure the red line indicates the median, while the green lines indicate the quartiles. For statistical analysis of the data shown on all violin plots we performed Kruskal-Wallis tests followed by Dunn’s multiple comparisons tests (See also Supplementary Table 3).

Previous work has shown that cortical actin is essential for lumen formation and that polarized microtubules contribute to lumen development by forming a network that is actin dependent. At later stages when notochord cells perform bi-directional crawling the microtubule network is organized towards the leading edges of the ALE and PLE, a process which is essential to organize the acting-based protrusions of the leading edge ^16^. To assess whether actin and microtubule organization was perturbed in *Ano10*^CRISPR^ notochord cells, we imaged and analyzed the organization the microtubule binding protein ensconsin-3xGFP, the microtubule plus-end marker EB3-mNeonGreen and the actin marker LifeAct-tdTomato throughout tubulogenesis.

We found that at the onset of lumen formation microtubules labelled with ensconsin-3xGFP showed a bias for radial distribution in negative controls (Figure 3J’-J’
s’’;M), arrayed mostly parallel to the A/P axis, with a higher concentration at the lateral surface (Figure 3J’,J’’’) where the lumen emerges, as previously shown^16^. Circumferential non-radial microtubules were also present across the different domains of the cell (Figure 3J’’,J’’’). In *Ano10*^CRISPR^ notochord cells microtubules were strongly disrupted (Figure 3K’-K’’’). They formed fewer but thicker bundles with an increase in the abundance of circumferential non-radial bundles (Figure 3K’-K’’’). Notochord-specific expression of a rescue construct in *Ano10*^CRISPR^ notochord cells rescued the mutant phenotype (Figure 3L’-L’’’). Furthermore, we quantified the fraction of microtubules at the basal domains (Figure 3N). We found that this fraction is higher in *Ano10*^CRISPR^ notochord cells compared to negative controls and rescue cells (Figure 3N). Subsequently, we monitored and quantified microtubule dynamics during bi-directional crawling using the microtubule plus end marker EB3-mNeonGreen (Figure 3O-Q). *Ano10*^CRISPR^ notochord cells showed a significantly lower enrichment for EB3 comets in the ALE and PLE regions as compared to negative control and rescue cells (Figure 3R,S).

Subsequently, we analyzed the subcellular organization of the actin marker tdTomato-LifeAct across the three genotypes used in our study (Figure 3T-V). First, we analyzed actin distribution during lumen initiation (Figure 3W,X) by measuring actin intensity in the median confocal sections (Figure 3W, X). We found that in negative controls actin is enriched in the apical/luminal domains as previously reported^16^(Figure 3X). In contrast, *Ano10*^CRISPR^ notochord cells showed a more uniform distribution of actin with a small peak around the middle of the cells (Figure 3X). Ano10 rescue cells adopted a similar intensity profile to negative controls (Figure 3X). Finally, we measured the fraction of actin localized to the leading edges of cells that are bi-directionally crawling and we found that actin was enriched in the leading edges as previously observed^16^ (Figure 3Y). The fraction of actin at the leading edges of the notochord cells was significantly reduced in *Ano10*^CRISPR^ notochord cells (Figure 3Y). Expression of a rescue construct restored actin at the leading edges to negative control levels (Figure 3Y).

Given the importance of polarized actin and microtubules in the apical/lateral domains for lumen expansion our results raise the possibility that the *Ano10*^CRISPR^ lumen expansion defect is in part due to deranged microtubule and actin localization. Similarly, the reduced presence of microtubules in the leading edge of *Ano10*^CRISPR^ bi-directionally crawling cells suggests that the regulation exerted by Ano10 on the organization and dynamics microtubule and actin networks is important for notochord cell behaviors during tubulogenesis.

### *Ano10* modulates Ca^2+^ dynamics in the notochord during CE and tubulogenesis

Anoctamins have been shown to modulate intracellular Ca^2+^ dynamics across a range of tissues^59-62^. In *Ciona robusta*, Ca^2+^ activity has been reported in the presumptive notochord area during cell intercalation ^63^. We therefore asked whether the Ca^2+^ activity is modulated by *Ano10*. We imaged Ca^2+^ dynamics in notochord cells of negative control and *Ano10*^CRISPR^ embryos by expressing the Genetically Encoded Calcium Indicator (GECI) GCaMP6s^64^ under the control of the Brachyury promoter. We observed that negative control animals exhibited robust Ca^2+^ dynamics during convergent extension (Figure 4A-C) and lumen formation (Figure 4 H,I). *Ano10*^CRISPR^ embryos also exhibited Ca^2+^ transients during the same developmental periods. To determine whether Ano10 contributes to Ca^2+^ signaling in the notochord we compared standard Ca^2+^ transient peak features between control and *Ano10*^CRISPR^ embryos (Figure 4D-G; 4J-M). During convergent extension, Ca^2+^ dynamics in *Ano10*^CRISPR^ were characterized by a small but statistically significant increase in peak amplitude and falling slope values (Figure 4D,F), however we observed that the frequency of Ca^2+^ transients in *Ano10*^CRISPR^ was lower compared to controls (Table 1). During the process of tubulogenesis *Ano10*^CRISPR^ Ca^2+^ activity was characterized by higher amplitude, steeper rising and falling slopes (Figure 4J-L) and a significantly shorter peak duration (Figure 4K). The frequency of Ca^2+^ transients in *Ano10*^CRISPR^ was comparable to negative controls (Table 1). Our data suggest that Ano10 may play a small role in dampening the max amplitude of Ca^2+^ transients, but it likely plays an important role in the generation of Ca^2+^ transients characterized by a long duration, with slow rise and fall kinetics.

**Figure 4.**
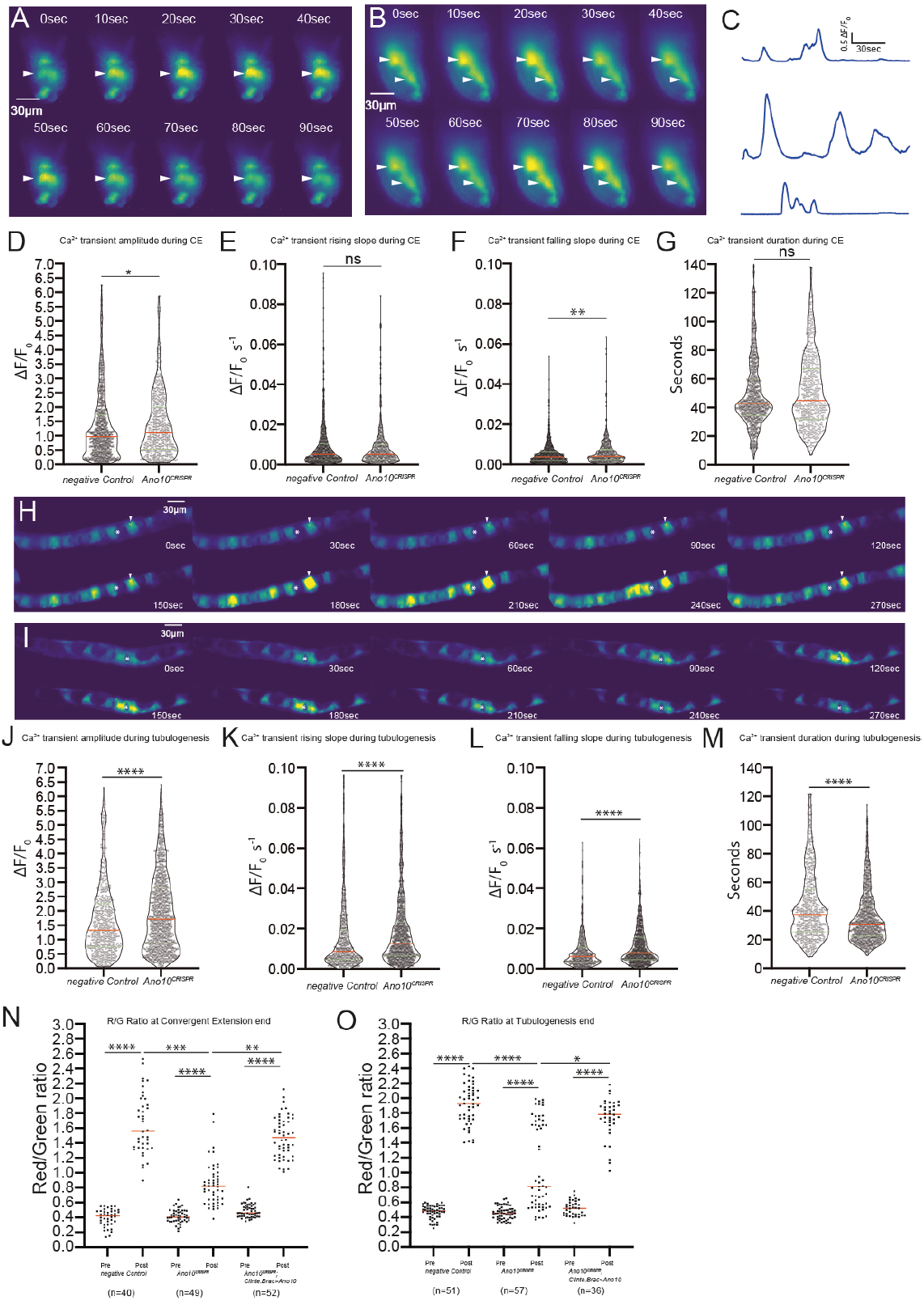
Ano10 modulates Ca^2+^ dynamics during convergent extension and lumen formation and extension. (A-B) Montage of two time-lapse movies from different embryos expressing *Ciinte*.*Brac>GCaMP6s* during notochord cell intercalation. White arrowheads point to notochord cells with changing fluorescence intensity in response to changes in Ca^2+^ concentration (Supplementary Movies 4-8). (C) Three example Ca^2+^ traces from notochord cells. (D-G) Quantification of features from negative control and *Ano10*^CRISPR^ Ca^2+^ transients during convergent extension. Violin plots for (D) amplitude (E) rising slope, (F) falling slope (G) duration. The red line indicates the median and green lines indicate the quartiles. (H, I) Montage of two-time lapse movies from different embryos expressing *Ciinte*.*Brac>GCaMP6s* during lumen formation and extension. (J-M) Violin plots quantifying (J) amplitude (K) rising slope, (L) falling slope and (M) duration of Ca^2+^ transients from control and *Ano10*^CRISPR^ embryos during tubulogenesis. For statistical analysis of data shown in panels ((D-G; J-M) we performed Mann-Whitney tests (see also Supplementary Table 4). (K,L) Quantification of Red/Green ratio of CAMPARI Ca^2+^ integrator before (pre) and after (post) photoconversion at the end of CE (K) and tubulogenesis (L). For statistical analysis we performed a Kurskal-Wallis test followed by Dunn’s multiple comparisons test (see also Supplementary Table 4)

A limitation of our Ca^2+^ imaging method is the short duration of the recordings (5 minutes per recording) which does not allow us to assess differences in Ca^2+^ activity between negative control and *Ano10*^CRISPR^ notochords over extended developmental time-windows. To overcome this limitation, we used the photoconvertible Genetically Encoded Integrator CAMPARI2 which can integrate Ca^2+^ levels over extended periods of time^65^. Photoconversion of green fluorescence from CAMPARI2 to red fluorescence in the presence of UV light and Ca^2+^ ions means that CAMPARI2 can act as a ratiometric integrator, since the Red/Green ratio values correlate with the levels of accumulated Ca^2+^ activity in the cell/tissue. We imaged embryos expressing CAMPARI2 in the notochord either at the onset of convergent extension or tubulogenesis (pre data points), we then illuminated the embryos periodically with low levels of UV light and imaged again at end of convergent extension or tubulogenesis (post). We quantified the Red/Green ratio (R/G) of negative control, *Ano10*^CRISPR^ and rescue embryos. As expected, before photoconversion we obtained a low R/G across all conditions (Figure 4N,O). Following photoconversion at the end of convergent extension or tubulogenesis negative controls showed a large increase in R/G ratio suggesting that the notochord cells have exhibited high Ca^2+^ activity during these developmental windows (Figure 4N,O). *Ano10*^CRISPR^ cells exhibited an increase in R/G ratio, however this was significantly smaller than that of negative controls or rescues (Figure 4N,O) indicating that loss of Ano10 results in reduced Ca^2+^ activity over these two developmental windows. Taken together our data suggest that Ano10 regulates Ca^2+^ activity during convergent extension and tubulogenesis. Aberrant Ca^2+^ activity in *Ano10*^CRISPR^ may underly the defects in convergent extension and tubulogenesis exhibited by *Ano10*^CRISPR^ mutants.

### Ano10 functions as a channel and likely not as a scramblase

Next, we evaluated the ability of Ano10 to act as a phospholipid scramblase in cell culture. Phosphatidylserine (PS) a type of phospholipid, is normally constrained to the inner leaflet of the plasma membrane and gets exposed to the outer leaflet under various conditions and stimuli including large increases in Ca^2+^ concentrations inside cells. Phospholipid scramblases including certain Anoctamin family members are responsible for the translocation of PS molecules between the inner and outer layers of the plasma membrane^38,46^.

We expressed Ano10-GFP (Figure 5C, C’, C’’; Table 2), or CAAX tag fused Ano10-GFP (targeting the protein to the PM) (Figure 5D, D’, D’’; Table 2) in HEK293T cells and manipulated the intracellular Ca^2+^ concentration with ionomycin. We then stained with AnnexinV-Alexa Fluor 568 antibody to detect PS activity. In both instances, we failed to observe phospholipid scramblase activity at higher levels compared to mock transfected controls(Figure 5A,A’,A’’; Table 2). Our positive control mAno6 yielded robust scramblase activity (Figure 5B,B’,B’’; Table 2). Importantly, during the process of screening *C. intestinalis* Anoctamins for scramblase we discovered that Ciona Ano6 displayed strong scramblase activity (Figure 5 E,E’,E’’) indicating that *C*.*intestinalis* Anoctamins can be functional and show scramblase activity in a heterologous context.

**Figure 5.**
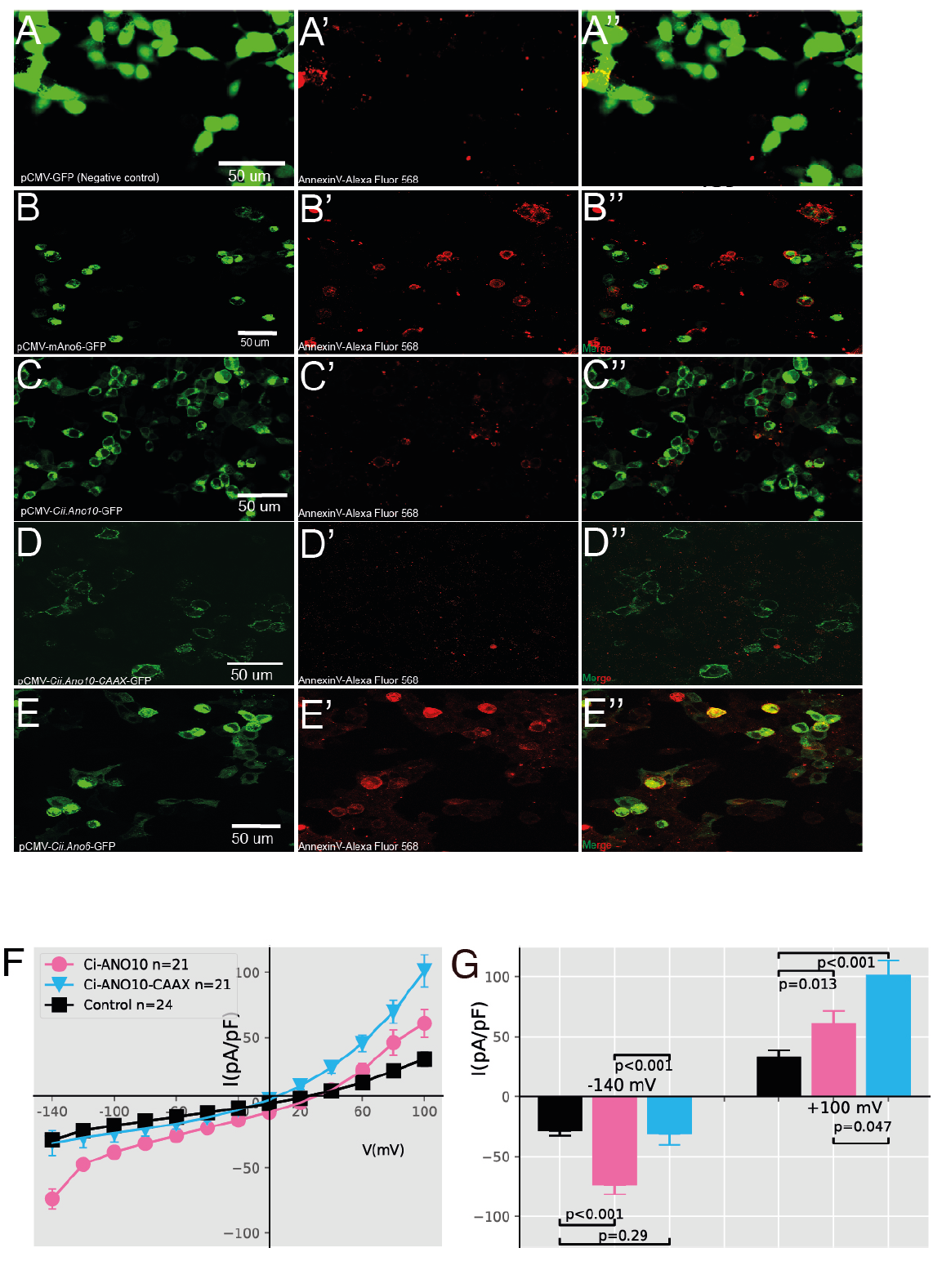
Ano10 functions as a channel and likely not as a scramblase. Representative confocal micrographs showing the results of the phospholipid scramblase assay in HEK293T cells. overexpressing (A, A’, A’’) pCMV-GFP (negative control), (B,B’,B’’) pCMV-mAno6-GFP (positive control), (C,C’,C’’) pCMV-Cii.Ano10-GFP; (D,D’,D’’) pCMV-Cii.Ano10-CAAX-GFP and (E,E’,E’’) pCMV-Cii.Ano6-GFP. The left column shows GFP expression from the transfected constructs. The middle column shows Annexin V-Alexa Fluor 568 signal. The right column is the merge of the two channels. For quantification of the phospholipid scramblase assay result please see Table 2. (F) Stimulation with 100 μM ATP elicits currents in patch-clamp experiments for control and Cii.Ano10 and Cii.Ano10-CAAX expressed in HEK293T cells. Cells were held at 0 mV and pulsed to voltages from -140 mV to +100 mV (in steps of 20 mV). (G) Quantification of the current/voltage relationships between control, Cii.Ano10 and Cii.Ano10-CAAX expressing HEK293T cells. For statistical analysis we used Mann-Whitney tests (see also Supplementary Table 5).

Finally, we employed electrophysiology to determine whether Ano10 possesses an ion channel functionality. We co-expressed Ano10-GFP and the purinergic receptor P2Y2R (which is coupled to Gq signaling and results in an increase in intracellular Ca^2+^ upon stimulation by ATP) together in HEK293T cells and tested for current responses to ATP stimulation in a whole-cell configuration. Cells transfected with Ano10 exhibited a larger current than mock transfected, and the current-voltage relationship was nonlinear (Figure 5F). When we used the CAAX tag to tether Ano10 at the PM, we obtained larger currents at positive voltages (Figure 5F). Our experimental evidence argues in favor of Ano10 working as a channel rather than a phospholipid scramblase.

## Discussion

The present study has been designed to investigate the roles that Ano10 may play in the context of the developing *Ciona* notochord. It adds Ano10 to the collection of membrane proteins responsible for tubulogenesis^22-25,66^. Specifically, our study demonstrates that ANO10 is required for both convergent extension and tubulogenesis steps during notochord formation. *Ano10*^CRISPR^ notochord cells exhibit defective convergent extension, which is manifested in defects in cell morphology, lower rates of successful intercalation events, irregular positioning of the cells along the A/P axis and reduced A/P axis length. To the best of our knowledge ANO/TMEM16 proteins have not been previously implicated in the developmentally important process of cell intercalation^67^. Interestingly, members of the ANO/TMEM16 family are expressed in epithelial tissues that undergo cell intercalation during embryonic development^43,68^. For example, several murine Ano/Tmem16 gene family members including Ano10/Tmem16k are expressed in the neural tube^43^. This raises the possibility that some vertebrate Anoctamin proteins including ANO10/TMEM16K are involved in cell intercalation.

We additionally uncover a role for Ano10 in lumen expansion and lumen connection during but not lumen initiation during tubulogenesis. Previous work has shown that the anion transporter Slc26a*α* is specifically required for lumen expansion but not for lumen specification or lumen connection during *Ciona* notochord development^23^. Ano10 shows more pleiotropic phenotypes compared to Slc26a*α*, since Slc26a*α* is not involved in convergent extension and it is required for only one of the steps during lumen formation. However, neither of these transmembrane proteins seem to be involved in lumen initiation, thus the channels, transporters involved in this stage of lumen formation remain elusive. More generally, despite the fact that several Anoctamins have been shown to express in tubular structures such as kidney tubules, airways, intestine and different types of glands^37-43^, evidence for the involvement of Anoctamins in biological tube development has been very limited, with the notable exception of two studies that implicated ANO1/TMEM16A in epithelial morphogenesis^69^ and the development of the murine trachea^68^. Our work is complementary to these two studies, since it implicates a new member of the ANO/TMEM16 protein family in tubulogenesis and provides a quantitative analysis of the role of Ano10 in regulating the cell behaviors and cytoskeletal organization of the notochord across different stages of development. We hope that our study will prompt new studies on the role of TMEM16K/ANO10 and other Anoctamins in the development of tubular structures.

This study further demonstrates that Ano10 is important for regulating Ca^2+^ dynamics during notochord development. These effects on Ca^2+^ activity coincide with the two important morphogenetic steps of convergent extension and tubulogenesis, which are characterized by different cell behaviors and morphogenetic events. This raises the possibility that robust Ca^2+^ signaling which is at least in part mediated by Ano10 may act as a pleotropic signal integrator^70^ during notochord development.

More generally, our finding that *C. intestinalis* Ano10 contributes to Ca^2+^ homeostasis in the notochord is in agreement with previous studies in the field which have shown that mutations in Anoctamins, including ANO10 lead to deranged Ca^2+^ activity, which in turn results in pathological conditions^58-60,71^ For example, the defects in ion transport, cell volume regulation and cell migration exhibited by Ano10 knockout mice have been attributed to deranged local Ca^2+^ signaling ^58^. Taking these different lines of evidence together raises the exciting possibility that at least some of the vertebrate ANO10/TMEM16K proteins may play a critical role in regulating Ca^2+^ dynamics during development in a broad spectrum of biological tissues.

Different studies have indicated that vertebrate ANO10 can function either as a scramblase or an ion channel ^72,73^. An *in vitro* study has suggested that ANO10 may act as a CaCC^73^, however, other studies have reported that ANO10 is an ER-residing phospholipid scramblase ^72,74,75^.Our *in vivo* and *in vitro* work suggests that *C. intestinalis* Ano10 functions as a channel rather than as a phospholipid scramblase.

In conclusion, cells must integrate a diversity of local signals to coordinate tissue-level processes. Our study suggests that Ano10 is likely part of the machinery that integrates these signals and coordinates the generation of appropriate cellular behavioral output to build one of the defining characteristics of all chordates, the notochord.

## Methods

### Animal collection

Adult *C. intestinalis* (Type B) were collected from: Døsjevika, Bildøy Marina AS, postcode 5353, Bergen, Norway. The GPS coordinates of the site are the following: 60.344330, 5.110812.

### Rearing conditions for adult *C. intestinalis*

Adult C. intestinalis were housed in a purpose-made facility at the Sars Centre. Approximately, 50 to 100 adults were housed in large 50L tanks with running sea water. The temperature of the sea water was maintained at 10°C with constant illumination and food supply (various species of diatoms and brown algae) to enhance egg production and prevent spawning^76^.

### Electroporations of zygotes and staging of embryos

Egg collection, fertilization as well as rearing were done following standard protocols^77^. Gravid adult *C. intestinalis* were dissected to obtain mature eggs and sperm to perform fertilization *in vitro*. Eggs were dechorionated using chemical dechorionation in a 1% sodium thioglycolate (Sigma Aldrich) and 0.05% pronase (from Streptomyces griseus, Sigma Aldrich) mix dissolved in filtered seawater. Eggs in the dechorionation solution were placed on a rocker for approximately 6 min until zygotes were completely dechorionated. Dechorionated eggs were washed several times using artificial sea water and then fertilized with sperm for ∼10 min. Electroporation was performed according to published protocols with minor modifications ^78^. After thoroughly washing, the fertilized eggs were electroporated in a mannitol solution with 100 to 200 *μ*g of DNA depending on the expected levels of expression for each given construct. We electroporated fertilized eggs in electroporation cuvettes with a 4 mm gap (MBP Catalog #5540) using a BIORAD GenePulserXcell equipped with a CE-module. The settings we used were as follows: Exponential Protocol: 50 V, Capacitance was set to: 800–1300*μ*F, Resistance was set to ∞ and we aimed to achieve an electroporation time constant between 15–30 milliseconds. Embryos were cultured in ASW (artificial sea water, Red Sea Salt) at 14°C until they were used for experiments. The pH of the ASW was set to 8.4 at 14°C. The salinity of the ASW we prepared was set to 3.3–3.4%. Embryos were cultured during early development (zygote to stage 13^79^) at 14°C. For most live-imaging experiments we tried to ensure that the temperature would stay stable around 18°C using room and/or microscope cooling.

### Molecular biology procedures to obtain expression constructs for cell culture and C. intestinalis transient transgenesis

The primers for cloning *C. intestinalis Ano10* cDNA were designed according to the *Ciona intestinalis* genome databases (Aniseed Database gene model KH2012:KH.C3.109; and Ghost Database KY21.Chr3.1036 80,81). A FLAG-tag sequence as DYKDDDDK was directly added to the primers for *Ano10* cDNA amplification. Around 2kb Ano10 cDNA was amplified by PCR using Norwegian *Ciona intestinalis* cDNA as the template. The PCR amplified cDNA fragments were cloned into PEGF-N1 vector (Clontech,6085-1,) with XhoI/BamhI enzyme sites. To generate the CAAX tagged *C. intestinalis Ano10*, CAAX tag sequence KKKKSKTKCVIM was directly added to the primers for *Ano10* cDNA amplification, then followed with the same PCR and ligation process as mentioned above.

For experimental use in *C. intestinalis* transgenic embryos the *Ano10* cDNA was amplified We then using cDNA at a concentration of 100ng/*μ*l, a dNTPs mix (Thermofisher, R0182) and Q5 High-Fidelity DNA Polymerase (M0491L, NEB) to perform the PCR reaction. Subsequently, we gel purified the PCR products using Zymogclean Gel DNA Recovery Kit (Zymo research, D4002). The purified product was inserted into the pDONR221 vector using BP Clonase II (Invitrogen, P/N56480). We identified positive clones using restriction digest and we sequenced multiple clones with Sanger sequencing. Both the cDNA and the promoter sequence of *Ano10* were cloned into the Gateway system for experimental use in *C. intestinalis*. We inserted the PCR products of *Ano10* cDNA into PDONR221 vector and the promoter into pDONR P4-P1R vector with BP Clonase II (Invitrogen) respectively. Expression vectors were recombined from LR reaction with pDESTII vector and LR Clonase II (Invitrogen).

To amplify promoters for the following genes: *Brachyury*, Ano10, Carbonic anhydrase 2 (Aniseed Database gene model KH.C1.423; Ghost Database KY21.Chr1.1715) we extracted genomic DNA from local Norwegian animals using the Wizard Genomic DNA Purification Kit (A1120, Promega). We then used the purified gDNA at a concentration of 100-150ng/*μ*l, the primers shown in Supplementary Table, a dNTPs mix (Thermofisher, R0182) and Q5 High-Fidelity DNA Polymerase (M0491L, NEB) to perform PCR reactions. Subsequently, we gel purified the PCR products using Zymogclean Gel DNA Recovery Kit (Zymo research, D4002). These were inserted into the P4-P1R vector using BP Clonase II (Invitrogen, P/N56480). We identified positive clones using restriction digest and we sequenced multiple clones with Sanger sequencing.

For the middle position we used hCD4GFP, GCaMP6s, nls::Cas9::nls (subcloned from Eef1a-1955/- 1>nls::Cas9::nls a gift from Lionel Christiaen, Addgene plasmid # 59987 ;http://n2t.net/addgene:59987 ; RRID:Addgene_59987)^52^, EB3-mNeonGreen^82^ a gift from Dorus Gadella (Addgene plasmid # 98881 ; http://n2t.net/addgene:98881 ; RRID:Addgene_98881), TdTomato-LifeAct tdTomato-Lifeact-7 was a gift from Michael Davidson (Addgene plasmid # 54528 ; http://n2t.net/addgene:54528 ; RRID:Addgene_54528). Primers for all subcloned constructs are shown in Supplementary Table 8. We performed a four-way Gateway choosing one of the promoters at a time in the first position, with an appropriate middle position entry and unc-54 3’ UTR in the third position. These were recombined into a pDEST II backbone using LR Clonase II (Invitrogen, P/N56485).

All expression constructs generated in this work were then midi-prepped using NucleoBond Xtra Midi kit (Macherey-Nagel 740410.50).

### Whole mount in situ hybridization

For in situ hybridization targeting *Ano10* mRNA, 700bp *Ano10* cDNA was amplified and inserted into pCRII-TOPO vector (TOPO Cloning Kit, Thermofisher). Primers are shown in Supplementary Table 8. The antisense DNP-UTP (Roche) labeled probe for *Ano10* was generated by *in vitro* transcription with MEGAscript™ SP6 Transcription kit (Invitrogen). Whole-mount in situ hybridization was performed according to the procedure described by Christiaen et al^78^. Embryos and probes were hybridized at 55°C for 16–48 hrs with 200ng/ml of probe. For imaging, embryos were suspended in 2% DABCO/50% glycerol in PBS and mounted on slides with zero thickness coverslips. We imaged our samples were on a Nikon Eclipse E800 upright compound microscope. We performed three independent experiments, and we included negative controls (hybridization with a sense Ano10 probe).

### Phylogenetic analysis

89 orthologous Anoctamin protein sequences from 22 species (*Homo sapiens, Mus musculus, Xenopus tropicalis, Danio rerio, Petromyzon marinus, Eptatretus burgeri, Ciona intestinalis, Saccoglossus kowalevskii, Asterias rubens, Capitella teleta, Drosophila melanogaster, Oikopleura dioica, Branchiostoma floridae, Acyrthosiphon pisum, Strongylocentrotus purpuratus, Nematostella vectensis, Lottia gigantea, Caenorhabditis elegans, Aspergillus fumigatus, Nectria haematococca, Saccharomyces cerevisiae, Dictyostelium discoideum*), which were longer than 300 amino acids were downloaded from Uniprot^83^. 4 *Ciona intestinalis* Anoctamins were searched and download from *Ciona* genome database (Ghost^81^ and Aniseed^80,84^). Amino acid alignments were carried out by using the online version of MAFFT (https://www.ebi.ac.uk/Tools/msa/mafft/). Poorly aligned regions of the multiple protein alignment of Anoctamins were removed with Gblocks (http://phylogeny.lirmm.fr/phylo_cgi/one_task.cgi?task_type=gblocks) with the least stringent parameters. Maximum likelihood (ML) phylogenetic analyses were conducted with RAxML v8 (https://raxml-ng.vital-it.ch/#/) with the autoMRE option to calculate the bootstrap support values. The tree was manipulated and annotated by iTOL (https://itol.embl.de/). Through phylogenetic analysis, we named four *Ciona* Anoctamins respectively by KH.C3.109 / KY21.Chr3.1036 as *C. intestinalis Ano10*, KH.C8.274 / KY21.Chr8.1253 as *C. intestinalis Ano5*, KH.C10.524 / KY21.Chr10.513 as *C. intestinalis Ano6*, and KH.C8.585 / KY21.Chr8.1090as *C. intestinalis Ano7*.

### RNA isolation and cDNA preparation

Total embryos’ RNA was extracted at different developmental stages using TRIZOL (Thermofisher) according to the manufacturer’s instructions. Contaminating DNA was degraded by treating each sample with Dnase (Roche) then heating up to 95°C to eliminate the enzyme activity. The amount of total RNA was measured by Nanodrop (ND-1000 UV–Vis spectrophotometer; NanoDrop Technologies) according to the absorbance at 260 nm and the purity by 260/280 and 260/230 nm ratios. 500 ng of total RNA was retrotranscribed with the kit (SuperScript™ IV First-Strand Synthesis System, Invitrogen) for each sample according to the manufacturer’s instructions.

### Generation of negative control and Ano10^CRISPR^ mutant embryos by CRISPR/Cas9

The guide RNA(gRNA) sequence targeting *C. intestinalis Ano10* was designed through online tools E-CRISP (http://www.e-crisp.org/E-CRISP/)^85^ with relaxed selection and CRISPOR (http://crispor.tefor.net/)^86^. 6 pairs of gRNAs were designed, and only 1 pair which target GTTTAAATGAGGTAACGCAG sequence on Exon 6 finally was used in subsequent experiments. The negative control gRNA was designed to not target any sequence in the *C. intestinalis* genome. The gRNA oligonucleotides were inserted into the U6>sgRNA(F+E) vector which was a gift from Lionel Christiaen (Addgene plasmid # 59986 ; http://n2t.net/addgene:59986 ; RRID:Addgene_59986) ^52^. Primers used to build reagents used in these experiments are shown in Supplementary Table 8. 30*μ*g *Cii*.*Brachyury>nls::Cas9::nls* or Carbonic anhydrase 2>nls::Cas9::nls and 70*μ*g *U6>Ano10gRNA* or *U6>ControlgRNA* plasmids were electroporated together into fertilized, dechorionated eggs. Embryos were fixed for genomic DNA extraction (JetFlex™ Genomic DNA Purification Kit, Thermofisher) according to the manufacturer’s instructions. To detect the mutation, we used a melting curve assay, using single embryo genomic DNA as our template (CFX Connect Real-Time PCR Detection System of Bio-rad). Positive samples from the melting curve assay were cloned into pCRII-TOPO vector and sent for sanger sequencing. The cleavage detection assay was done as described by Stolfi et al.^52^ (GeneArt Genomic Cleavage Detection Kit, Thermofisher). The efficiency of CRISPR/Cas9 editing was ≥39.0% across multiple independent experiments. For rescue experiments, we synthesized (GeneCust, France) an Ano10cDNA where the gRNA target site was replaced with the sequence GTTTAAACGAAGTAACACAA. We used synonymous codons thus we did not change the amino acid sequence encoded by the cDNA. Rescue constructs were electroporate in the same mix as *Cii*.*Brachyury>nls::Cas9::nls* or Carbonic anhydrase 2>nls::Cas9::nls and *U6>Ano10gRNA* at 60-80*μ*g.

### Embryo dissociation and MACS sorting

Transgenic embryos harboring relevant transgenes for negative control or Ano10^CRISPR^ conditions as well *Ciinte*.*Brac>CD4::GFP* were collected at Hotta stages 24∼25^79^. The embryos were incubated in artificial seawater with 0.2% trypsin (Thermofisher) and cell dissociation was facilitated by pipetting up and down the mix. After approximately 4 to 6 minutes, we stoped the trypsin reaction by adding 0.05% bovine serum albumin (BSA) (Thermofisher). The solution was filtered with 35 *μ*m cell strainer cap in 50 ml falcon tubes to remove non-dissociated cells. The supernatant was removed after centrifugation at 4°C with a speed of 3000RCF. Then the remaining sample was washed twice with artificial seawater containing 0.05% BSA. The cells were resuspended in 180 *μ*l artificial seawater containing 0.05% BSA and we added 20 *μ*l of CD4 antibody (Miltenyi Biotec) conjugated with magnetic beads. The sample tubes were kept for 1 hour at 4°C and then centrifuged at 1000RCF at 4°C for 3 minutes. After washing three times, the cells were resuspended with artificial seawater containing 0.05% BSA. We performed the CD4 positive cell sorting using the OctoMACS starting kit (MIltenyi Biotec) following the manufacturer’s protocol.

### Immunostainings

*Ciona* transgenic embryos used for Ano10 protein subcellular localization experiments were fixed in 4% Paraformaldehyde (PFA) at room temperature for 15 minutes. We then blocked with 10% heat-inactivated goat serum in PBS for 2 hours at room temperature. Commercial anti-FLAG antibody (Sigma-Aldrich) was used at 1:50 dilution and incubated for 2 hours at room temperature. Secondary goat anti-mouse antibodies conjugated with Alexa Fluor 488 (Invitrogen) was used at 1:200 dilution. Cell nuclei were stained with DAPI (Invitrogen) at 1:1000 dilution in PBT solution. For determining the integrity of the notochord border we performed immunostaining with an antibody against laminin (Merck L9393-100UL). Embryos were fixed in 4% PFA at room temperature for 15-20 minutes. We then blocked with 10% heat-inactivated goat serum in PBS for 2 hours at room temperature. We then transferred the embryos to a PCR tube (with a max capacity of 250*μ*l) applied the anti-laminin antibody at 1:50 dilution and incubated overnight at room temperature with gentle aggetation. A secondary antibody mouse anto-rabbit was applied. This was conjugated with Alexa Fluor 568 (Invitrogen) and was used at 1:200 dilution.

### Tissue culture and cell transfection

HEK293T cells and HeLa cells were cultured in DMEM and DMEM/F12(Gibco™), respectively which were both supplemented with 10% fetal bovine serum, 2 mM L-glutamine, 100 U/ml^-1^ penicillin, and 100 μg/ml^-1^ streptomycin. All transfections were carried out by Lipofectamine™ Transfection Reagent (Invitrogen) following the manufacturer’s protocol. All experiments were performed between 36-48 hours after the transfection.

### Electrophysiology

Electrophysiological studies were carried out in HEK293T cells. The cells were seeded on the round coverslip coated by poly l-lysine. The fluorescence positive cells were selected under microscopy(Zeiss). Recordings were performed using the whole-cell patch-clamp configuration. Patch pipettes had resistances ranging from 3–6 MΩ when filled with the pipette solution. Data were acquired by a HEKA EPC10 amplifier and Patchmaster software (HEKA). Current measurements were performed at room temperature. Capacitance and access resistance were monitored continuously.

Currents were filtered at 2.9 kHz with a low-pass Bessel filter. The Pipette (intracellular) solution contained (mmol/L): 146 CsCl, 5 EGTA, 2 MgCl2, 10 sucrose, 10 HEPES, PH was adjusted to 7.3 with NMDG. The extracellular solution contains (mmol/L): 140 NaCl, 5 KCl, 2 CaCl2, 1 MgCl2, 10 mM HEPES, 15 D-glucose, pH was adjusted to 7.4 with NaOH. Data were analyzed by python 3.8.

### Calcium imaging using GCaMP6s

A Zeiss Inverted microscope (ZEISS Axio Scope.A1) with a water immersion objective ZEISS W B-ACHROPLAN ×40 was used. Illumination was provided using a mercury lamp. The filter-set we used was as follows: BP470/20, FT493, BP505-530. Embryos with similar expression levels were selected under a dissecting fluorescent microscope. They were transferred to a 6 cm cell culture petri dish and embedded in 1% low melting point agarose (Fisher BioReagents, BP1360-100). Imaging was done at 18°C. Data was collected using a Hamamatsu Orca FlashV4 CMOS camera at 10Hz with an exposure time of 100 ms using the acquisition software AwesomeImager^87^. Each movie we acquired was 5 minutes long. Analysis was performed using the Ca^2+^ imaging analysis platform Mesmerize^88^. Briefly, images we motion-corrected using the NoRMCorre module and we performed signal extraction using the CNMFE module with the parameters optimized per video.

### Phospholipid Scramblase assay

For the cell culture phospholipid scramblase assay, we generated and transfected into HEK293T the following constructs pCMV-GFP (negative control), pCMV-mAno6-GFP (positive control), pCMV-*Cii*.*Ano10*-GFP; pCMV-*Cii*.*Ano10*-CAAX-GFP and pCMV-*Cii*.*Ano6*-GFP. Primers for reagents used in these experiments are shown in Supplementary Table 8. Following transfection we waited for 24 hours and we then seed the transfected HEK293T cells in a confocal imaging chamber (Mattek or ibidi chamber with 0 thickness coversleep, 22mm diameter). The cells were washed twice with PBS. Then they were incubated with 1:100 dilution of AnnexinV-Alexa Fluor™ 568 antibody in binding buffer (Thermofisher) for 10 minutes, which also contained 1x RedDot™1 Far-Red Nuclear Stain (Biotium) for dead cell detection and with 10 *μ*M ionomycin or DMSO. We then fixed the cells for 15 minutes with 4% Paraformaldehyde (PFA). The cells were washed three times with PBS and imaged using a Leica TSP5 confocal. The data was analyzed using custom made scripts written in Python 3.8. We used machine vision to segment and extract the contours in the green (transfected construct tagged with GFP), red (Annexin v-AlexaFluor568) and magenta (RedDot1 Far-Red) channels. Manual curation of the contours was performed in ImageJ. Cell numbers of Green and/or Red positive cells were counted using Scikit-image in python 3.8. RedDot1 positive cells were excluded from the analysis.

### Quantification of notochord cell intercalation fraction, notochord length analysis and cell shape parameters

We electroporated *Ciinte*.*Brac>hCD4GFP* in combination with our negative control, Ano10^CRISPR^ or rescue electroporation mixes. At Hotta stage 14^79^ animals were were transferred to Mattek chambers (but NOT embedded in agarose, just immersed in ASW) and imaged using an OLYMPUS spinning disk confocal using a 30x silicon oil objective, illuminated with 488nm laser, with an exposure time of 400ms. Stacks approximately 80*μ*m wide with a z-step size of ca 1.5*μ*m were collected. Using time-lapse mode (with slow rolling of stage so that we don’t displace the embryos) up to 10 animals could be imaged every 15 minutes. We recorded for a maximum of 300 minutes. Temperature was maintained as close as possible to 18°C during the acquisition period. Resulting time lapse stacks were visualized in 3D using Imaris. To score the fraction of notochord cells intercalated over the first 300 minutes, we followed the approach of Veeman and Smith^8^ In brief, we scored each cell manually as being intercalated if the contacts it made with non-notochord cells formed a closed ring in contrast to non intercalated if these contacts formed only a segment of the ring. For notochord length quantification over time, we fixed animals from the same batch as those used for the time-lapse imaging from every 30 minutes grown at 18°C (stage 14 until stage 25 according to Hotta^79^ (0-600min) and imaged those using an OLYMPUS FV3000 point scanning confocal using 20x (air objective) and 30x objective (silicon oil immersion). We did not go beyond 600min post-onset of stg 14 since it was evident that Ano10^CRISPR^ embryos did not ‘catch up’ with negative control and rescue embryos (i.e. that this reduced notochord length was due to a developmental delay). We generated maximal projections of our stacks data and loaded these in Fiji. We used the segmented line tool to draw for each projection/time-point from the Anterior to the Posterior end of the notochord along the midline of the notochord. For the quantification of cell shape parameters we focused our analysis at notochord cells that exhibited mediolateral elongation and alignment as they began to intercalate. Using FIJI we selected a medial slice from each stack and we measured across multiple cells per stack the angle enclosed by the notochord cell protrusion and the aspect ratio of the cells.

### Quantification of notochord cell center to skeleton midline distance metric

We electroporated *Ciinte*.*Brac>hCD4GFP* in combination with our negative control, Ano10^CRISPR^ or rescue electroporation mixes. Embryos were allowed to develop until the end of stage 24 and then we fixed them. We then stained me with a conjugated Alexa Fluor 647 phalloidin antibody at a dilution of 1:250. Confocal stacks were collected using a Leica TSP5 confocal with the following settings: 40x NA1.25 oil objective and 488nm and 633nm Excitation lasers (intensity level 10-15%). Using the collected stacks we generated z-stack maximal projections. Leveraging custom made software we labelled the notochord cell centres from position 0-39 in increasing order along the Anterior-Posterior axis using the hCD4GFP signal to identify individual cells. Subsequently, we segmented the embryo outline using the phalloidin signal. For all the labelled cells, the distances to both the lateral outlines as well as the central skeleton line were calculated. Only the distance to skeleton was used in the manuscript. Comparisons of the distance to the central skeleton line for each cell number is an indication of intercalation quality. For statistical analysis we performed Mann-Whitney U-tests per cell number to determine statistical significance in distance to the skeleton line for negative control, Ano10^CRISPR^ and rescue.

### Notochord lumen curvature and area measurements

For the pharmacology based experiments we co-electroporated *Ciinte*.*Brac>hCD4::GFP* to label the plasma membrane and *Ciinte*.*Brac>Cav1:;mCherry* to label the prospective luminal domains of notochord cells each at 40μg to generate double transgenic embryos. For the genetic experiments, we electroporated Ciinte.Brac>hCD4GFP and *Ciinte*.*Brac>Cav1:;mCherry* in combination with our negative control, Ano10CRISPR or rescue electroporation mixes.

We monitored the development of the embryos at regular intervals until they reached the end of stage IV. At this point we transferred them to a Mattek chamber and they were embedded in low melting point agarose at a concentration of 0.5% in ASW, which did not inhibit their development and tail extension but was sufficient to limit their movement. For the drug experiments we incubated control animals in DMSO and all drugs were diluted in ASW at a final concentration of 100*μ*M. We imaged tubulogenesis using an OLYMPUS spinning disk confocal. We performed simultaneous two colour imaging using 488nm and 561nm lasers and an exposure of 300ms. We collected stacks of 100μm thickness. Multiple animals were imaged in every round. Each animal was revisited every 20 to 30 minutes. To measure average lumen curvature at the end of our recordings, we opened the acquired stacks in ImageJ and generated orthogonal views of the last acquired time point. We then used the Fiji plugin Kappa (https://github.com/fiji/Kappa) to obtain the average curvature values. We traced the shape of the lumen with a B-spline curve (type: closed, stroke thickness 1) using on average ten points and then used the data fitting algorithm: Point Distance Minimization.

### Measurement of morphology features in bi-directionally crawling notochord cells

We electroporated Ciinte.Brac>hCD4GFP in combination with our negative control, Ano10CRISPR or rescue electroporation mixes. Embryos dedicated to the morphology features measurements were allowed to reach bi-directional crawling stage, at which point they were fixed and mounted on slides. They were imaged using a FV3000 point scanning confocal with a 40x silicon oil objective. For the bi-directionally extending cells we quantified the leading-edge length by loading the confocal stacks in Fiji and measured in Fiji with the segmented line tool the sum of the distance corresponding to the Anterior LA plus NE length.

For live tracking and speed imaging of the notochord cells we electroporated Ciinte.Brac>hCD4GFP and *Ciinte*.*Brac>NUP50-mCherry* in combination with our negative control, Ano10CRISPR or rescue electroporation mixes. Late stage 23 embryos were mounted on Mattek chambers embedded n low melting point agarose at a concentration of 0.5% in ASW, which did not inhibit their development and tail extension but was sufficient to limit their movement. Imaging was done using an OLYMPUS SpinSR spinning disk confocal. We opted for a 30x silicon oil objective and illuminated with 488nm and 561nm lasers. We collected data on two cameras. The exposure time was set at 400ms. Embryos were imaged every 5 minutes for up to 7 hours (stage 24 till stage 27). Notochord cell movement was tracked using the Bitplane Imaris Software Spots module.

### Measurement of radial/non-radial organization of microtubules

We electroporated fertilized Ciona eggs with 60μg of *Ciinte*.*Brac>Ensconsin::GFP*^*16*^ (a kind gift from Dr. Di Jiang) in combination with the respective negative control, CRISPR and rescue plasmids. We fixed animals at the onset of stage V^18^ as lumen initiation occurred. We then used a Leica TSP5 confocal microscope to obtain confocal stacks of microtubules. We imaged a field of view (1024 × 512) using a HCX PL APO 40x 1.25NA Oil objective and laser power of 20%. We then generated maximal projections in ImageJ. To separate the images into radial and non-radial components we used a customized ImageJ macro (https://github.com/ekatrukha/radialitymap). We loaded the MAX projections and using the point button we selected the origin of coordinates (this is the center relative to what objects will be considered radial). We loaded the radialitymap plugin and selected the ‘Cubic Spline Gradient’ method and Tensor Sigma parameter of 8 pixels to separate the radial and non-radial image that correspond to the separated radial and non-radial components of the original projection.

Subsequently, we used both the original image and the two images showing the separated radial/non-radial components to generate radial intensity profiles. These were generated using the Radial Profile Angle plugin with the centre located at the centre of the notochord cell and the edge located at the most distant portion of the cell periphery. A 180° integration angle was used. In order to quantify the non-radiality of the microtubule organization for each cell, the areas under the curve (AUC) of the radial intensity profile of the original image (total intensity) and the AUC of the non-radial map image were calculated using GraphPad Prism. Using these values we calculated the non-radial intensity as a percentage of the total intensity: (AUCnon-radial/AUCTotal intensity) *100.

### Live imaging of microtubules using EB3-mNeonGreen

To image microtubule dynamics we used as a marker the *Cii*.*Brachyury>EB-3-mNeonGreen* reporter, which was electroporated at 40μg. This plasmid was added to the control, Ano10^CRISPR^ or rescue electroporation mix. Electroporated embryos were allowed to grow to the end of Stage IV. At this point they were transferred them to a Mattek chamber and they were embedded in low melting point agarose at a concentration of 0.5% in ASW, which did not inhibit their development and tail extension but was sufficient to limit their movement. Our imaging was performed using an OLYMPUS spinning disk confocal. We imaged using a 488nm laser and an exposure of 400ms using a 40x silicon oil objective. We collected stacks of 100μm thickness. Multiple animals were imaged in every round. Each animal was revisited every 5 minutes. Comet count analysis was performed using the ComDet plugin https://github.com/ekatrukha/ComDet in FIJI. We measured the number of comets across the entire cell and the number of comets at the leading edges. We then calculated the ratio of leading-edge comet intensity/ overall comet intensity across the entire cell.

### Actin distribution quantifications

To image actin distribution in the notochord we electroporated Ciinte.Brac>hCD4GFP (40*μ*g) and *Ciinte*.*Brac>TdTomato-Lifeact* (40*μ*g) in combination with our negative control, Ano10CRISPR or rescue electroporation mixes. Electroporated embryos were allowed to grow to the end of Stage IV. At this point they were transferred them to a Mattek chamber and they were embedded in low melting point agarose at a concentration of 0.5% in ASW, which did not inhibit their development and tail extension but was sufficient to limit their movement. Our imaging was performed using an OLYMPUS spinning disk confocal. We imaged using the 488nm and 561nm lasers and an exposure of 400ms using a 40x silicon oil objective. We collected stacks of 80 to 100μm thickness. Multiple animals were imaged in every round. Each animal was revisited every 5 minutes. First, we analyzed actin distribution during lumen initiation by measuring actin intensity in the median confocal sections by drawing a straight line across the Anterior-Posterior axis of each cell, crossing the apical domains. We then generated Maximal projections which were further analyzed in Fiji. To quantify the fraction of actin intensity at leading edge in bidirectionally crawling cells we traced the cell outlines using a freehand selection line. We measured the raw integrated density of the entire cell. Then we drew outlines around the leading edges and measured the raw integrated density. We then calculated the ratio of: raw integrated density (leading edges) / raw integrated density (entire cell).

### CAMPARI imaging and analysis

We co-electroporated *Ciinte*.*Brac>CAMPARI2* at 40-50*μ*g in combination with either negative control, Ano10CRISPR or rescue electroporation mixes. At stage 14 animals were transferred to Mattek chambers and imaged using an OLYMPUS spinning disk confocal using a 30x silicon oil immersion objective. We took a pre-photoconversion stack for each animal assayed using two-color acquisition mode, illuminating our samples with 488nm and 561nm laser lines. We then used a 405nm laser at 40% for 5 second periods every 5 minutes. After 300 minutes we took another stack for each animal using the 488nm and 561nm laser as the post-photoconversion stack. We used FIJI to generate maximal projections. We generated single cell ROIs around multiple cells in each notochord that we imaged. The same ROIs were used to measure Mean intensity in the Green (488nm) and Red (561nm) channel. We then calculated the R/G ratios for pre- and post-photoconversion cells.

### Statistical analysis

All statistical analysis was performed using GraphPad Prism. First we tested for normality using multiple normality tests (Shapiro-Wilk Test and Kolmogorov-Smirnov Test). For all of our violin plots data panels we found that at least for one of the conditions the data was not normally distributed.Thus we opted to use Kruskal-Wallis as a nonparametric test. For data panels with violin plots and a limited number of conditions, we then used Dunn’s Multiple Comparison test to compare the mean rank of each column (condition) with the mean rank of every other column (condition). We corrected for multiple comparisons using statistical hypothesis testing.

For line plots, we used a either a mixed-effects model or a two-way RM ANOVA with the Geisser-Greenhouse correction. In both instances we performed post-hoc Tukey’s multiple comparison test, to assess the significance of the differences between pairs of group means.

## Acknowledgments

We would like to thank Dr. Mie Wong for her feedback on the manuscript.

## Funding

This project was funded by a grant of the Research Council of Norway: grant number 234817 (Sars International Centre for Marine Molecular Biology Research, 2013-2022).

## Author contributions

Z.L. and M.C. designed the project. Z.L. and M.C. carried out the experiments. Z.L., D.D. and M.C. analyzed the data. Z.L. and M.C. wrote the paper with input from D.

---

**Figure S1.**
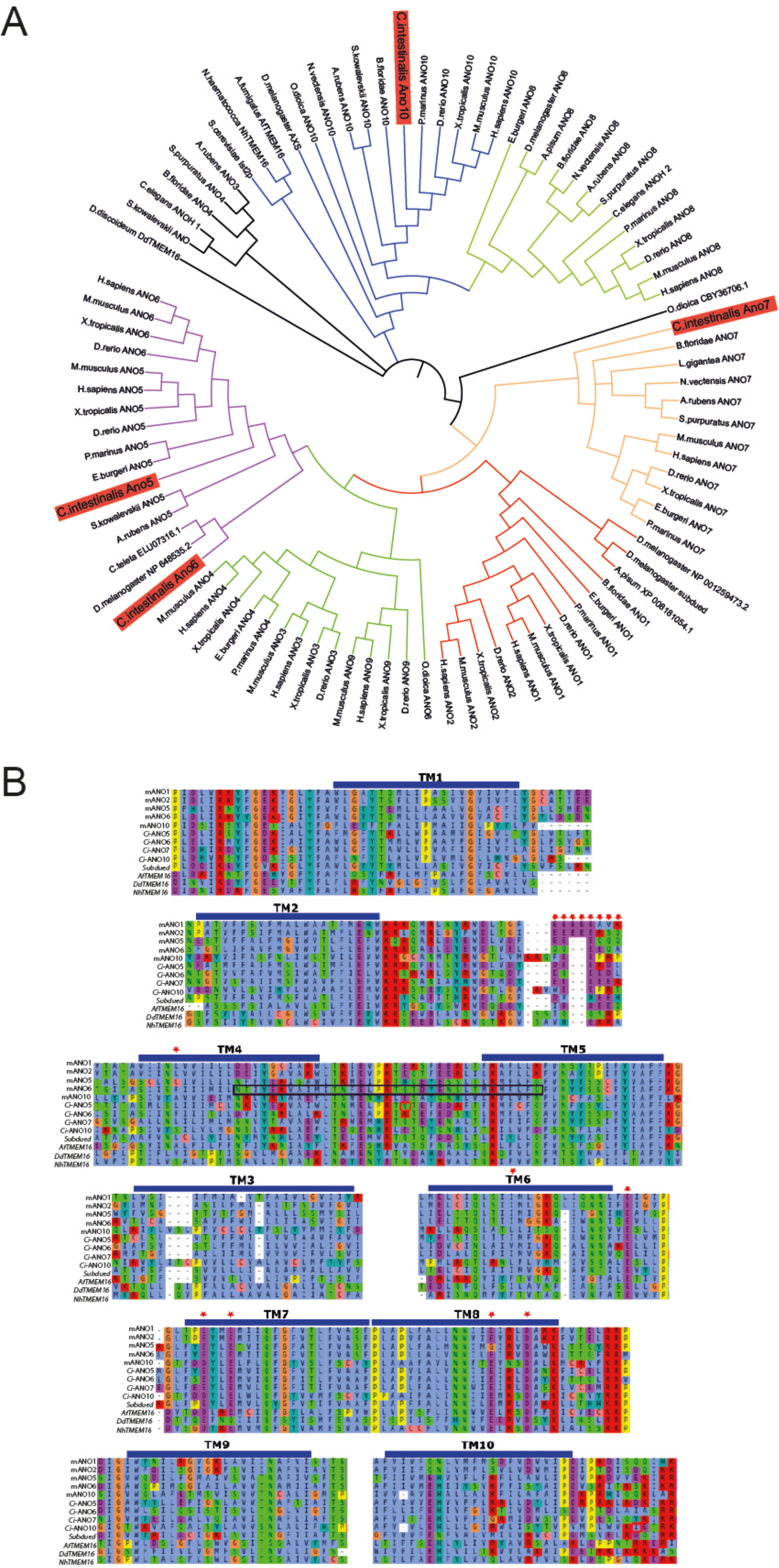
*C. intestinalis* Ano10 clusters together with vertebrate ANO10/TMEM16K. (A) Phylogenetic tree for the ANO/TMEM16 family using 89 sequences from a diversity of eukaryotic species. The different ANO/TMEM16 branches are labelled with the different colours. ANO/TMEM16 proteins from *Ciona intestinalis* are highlighted in red boxes. (B) Multiple sequence alignment for a subset of ANO/TMEM16 proteins which have been reported to exhibit phospholipid scramblase or ion channel activity, using Clustal W and Clustal X. The colours correspond to amino acid identity. The blue lines mark the putative transmembrane domains. The phospholipid scramblase domain of mANO6 is enclosed in the black box. Residues that are important for Ca^2+^ sensitivity are marked with red asterisks.

**Figure S2.**
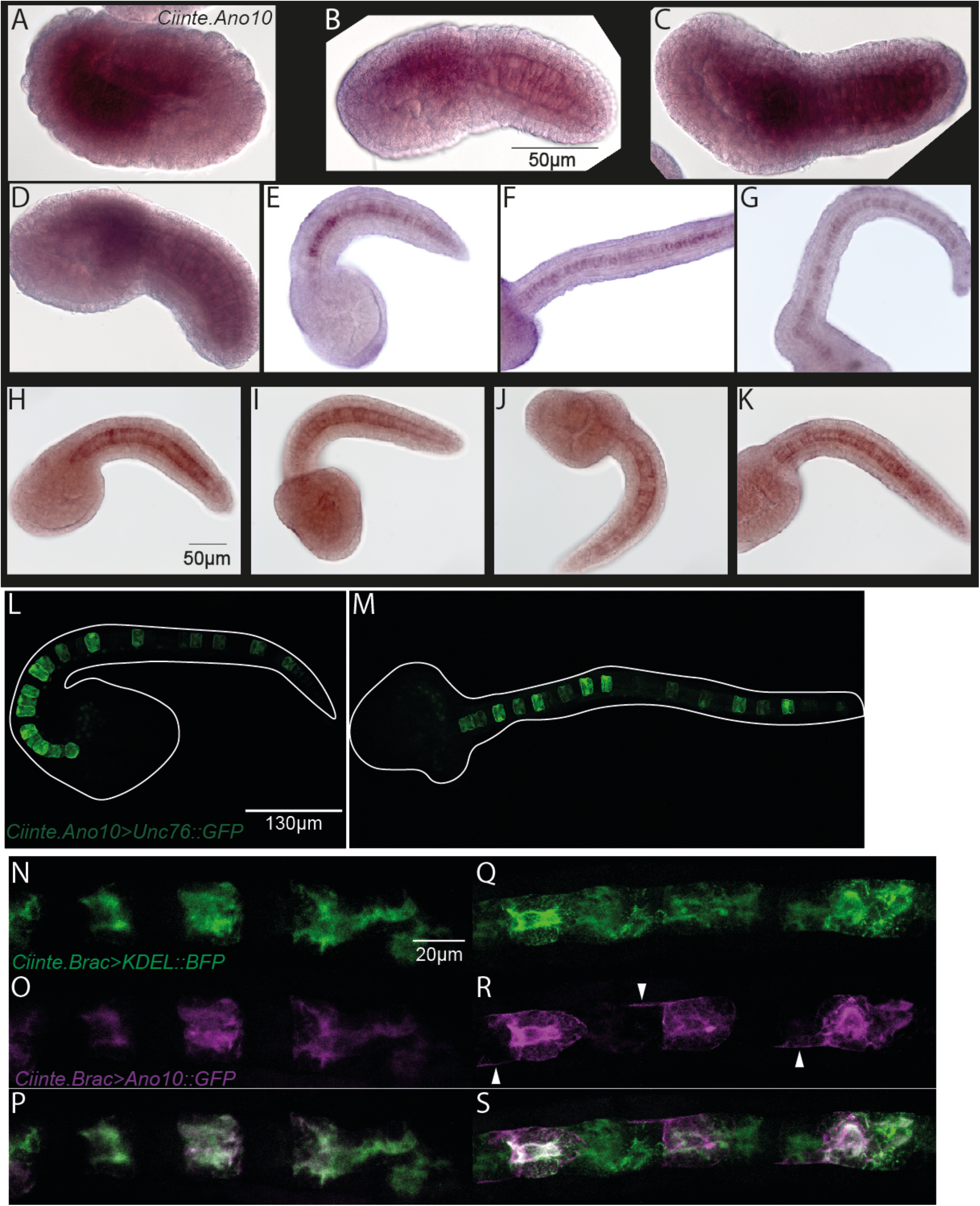
A*no10* is expressed in the notochord during embryonic development and localizes primarily to the ER. (A-K) Representative colorimetric whole-mount in situ with *Ano10* probe showing that initially *Ano10* is expressed broadly during neurula stages including expression in the endoderm, mesenchyme and presumptive notochord. During early, mid and late tailbud stages *Ano10* expression is restricted mostly to mesenchyme and the notochord. (L,M) A 2kb *Ano10* regulatory element drives GFP expression in the notochord during late tailbud stages. In the representative pictures we show maximal projection of late tailbud embryos. (N-P) At the end of notochord cell elongation ANO10 is localized primarily in the Endoplasmic Reticulum. Top panel shows expression of the ER marker *Ciintel*.*Brac>KDEL::BFP*; the middle panel shows *Ciinte*.*Brac>*Ano10::GFP in the same notochord cells. Bottom panel shows the merge between the two channels. (Q-S) During lumen connection and cell flattening Ano10 is expressed in the ER but also at the plasma membrane (marked by white arrowhead).

**Figure S3.**
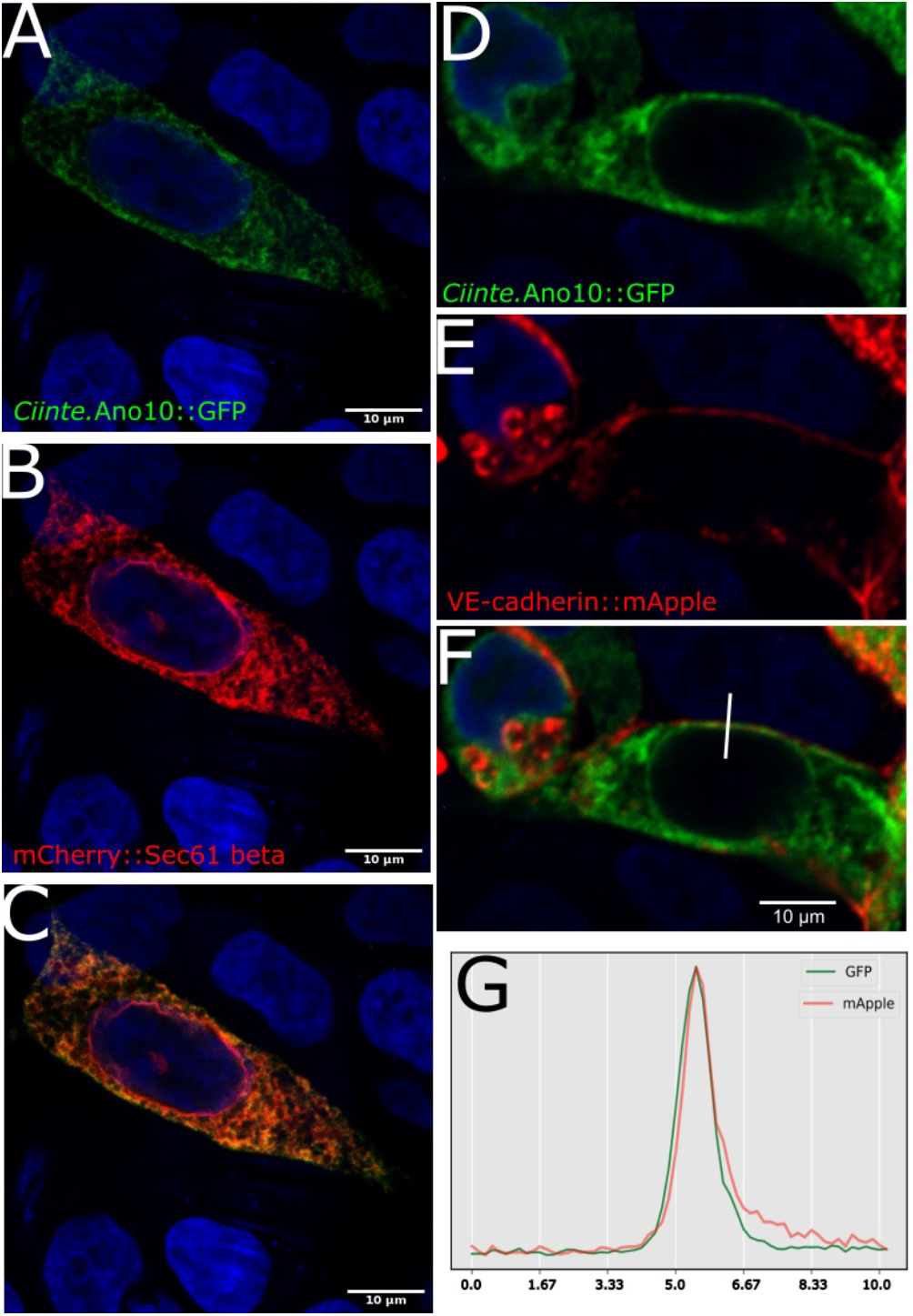
Ano10 localizes primarily to the Endoplasmic Reticulum but also to the Plasma Membrane of mammalian cells. (A) Representative confocal image of *C. intestinalis* Ano10::GFP expressed in HEK293T cells. (B) Representative confocal image showing the localization of the ER marker mCherry::Sec61 beta in HEK293T cells. (C) *C. intestinalis* Ano10::GFP and mCherry::Sec61 beta colocalize in the ER. (D) Additional representative confocal image of *C. intestinalis* Ano10::GFP expression in HEK293T cells. (E) Confocal image showing the localization of VE-cadherin::mApple in the same HEK293T cells. (F) A merge of the images in D and E. The white line corresponds to the site used for generating the intensity profiles shown in panel G. (G) Intensity profiles measured across the white line shown in panel F corresponding to *C. intestinalis* Ano10::GFP (green curve) and VE-cadherin::mApple (red curve).

**Figure S4.**
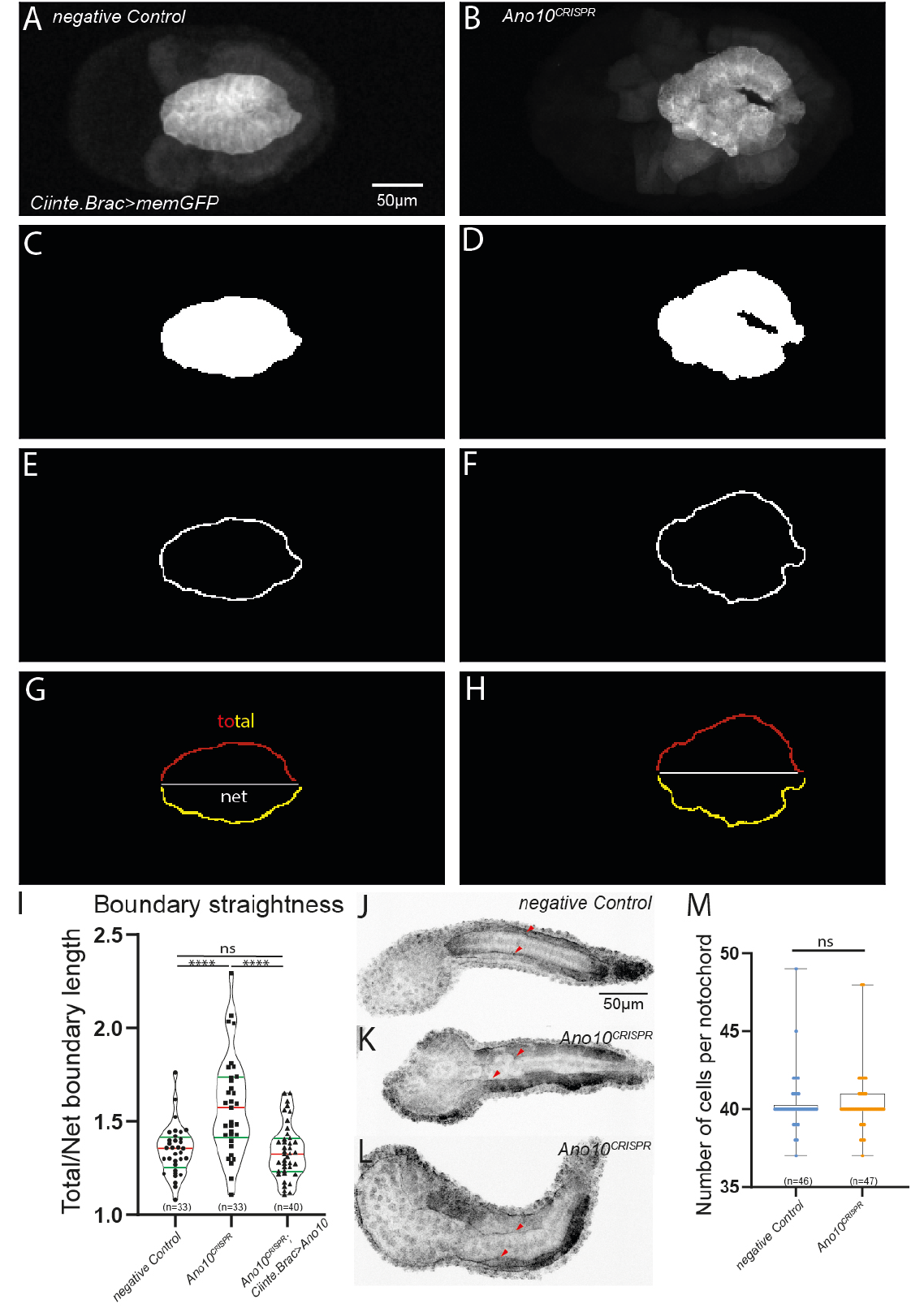
Ano10 is required for maintaining notochord boundary straightness. (A-H) Illustration of the notochord boundary regularity measurement approach. (A, B) Maximal confocal projection examples of negative control and *Ano10*^*CRISPR*^ embryos expressing *Ciinte*.*Brac>memGFP*. (C, D) Segmentation of the notochord using a binary mask operation (Watershed). (E, F) Notochord boundary outlines from the same embryos. (G,H) A straight line along the A-P axis is drawn to divide the notochord boundary to two sides. Both the net and total distance of the border are measured. (I) Quantification of notochord boundary straightness in negative control, *Ano10*^*CRISPR*^ and *Ano10*^*CRISPR*^;*Ciinte*.*Brac>Ano10* embryos as the ratio of Total to net length. n= number of animals analyzed. For statistical analysis we performed a Kruskal-Wallis test, followed by Dunn’s multiple comparisons test(****=<0.0001, ns= not significant). (J-L) Confocal projections showing immunostaining against laminin in negative control and *Ano10*^*CRISPR*^ embryos. Red arrowheads show the laminin staining along the notochord boundary. (M) Box plots quantifying the number of notochord cells in negative control and *Ano10*^*CRISPR*^ embryos. Statistical analysis was performed using Mann-Whitney test.

**Figure S5.**
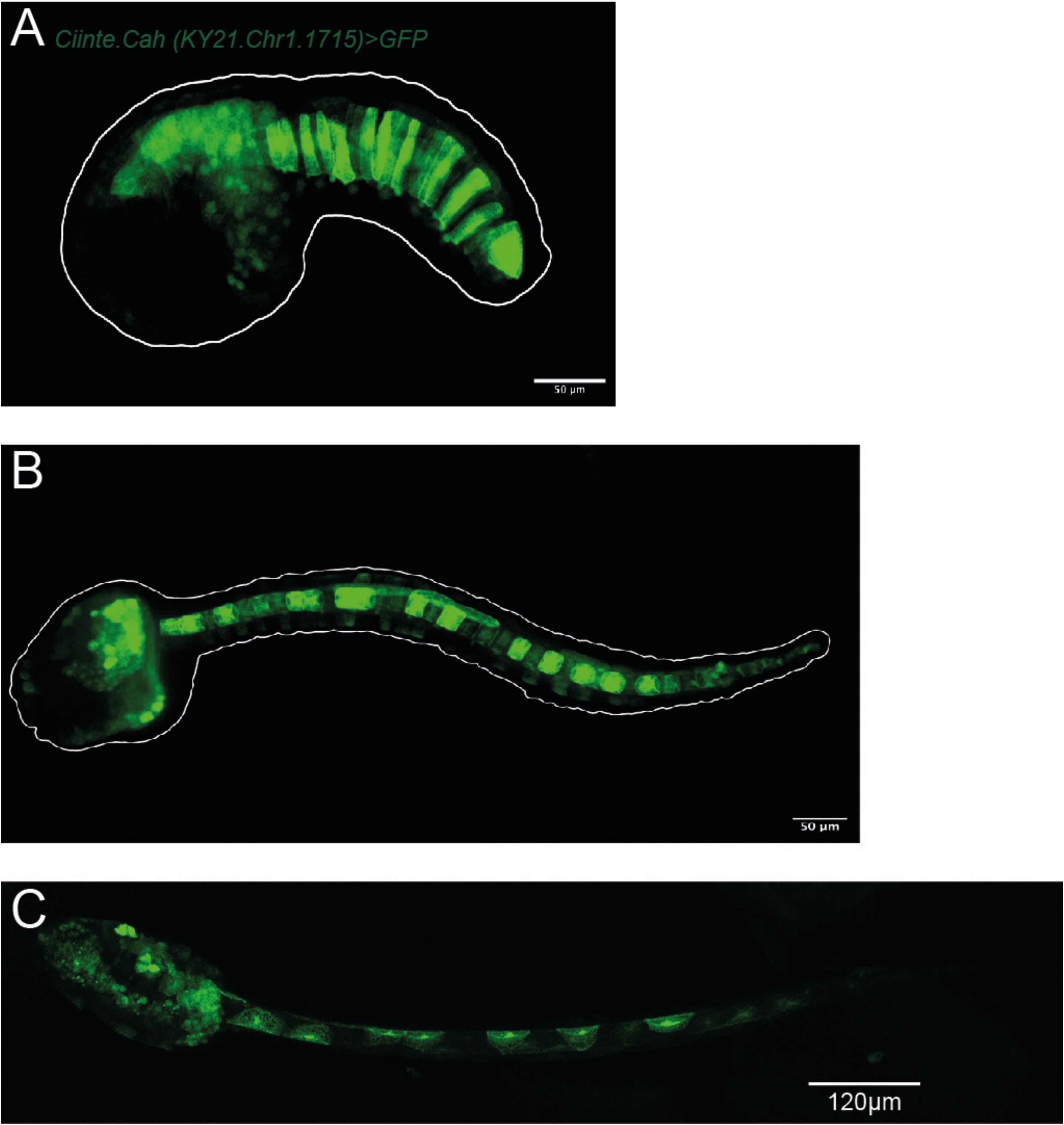
Expression pattern of carbonic anhydrase 2 KH.C1.423; KY21.Chr1.1715. (A-C) Maximal projections of confocal stacks of embryos electroporated with a plasmid harboring a 3kb fragment upstream of the carbonic anhydrase gene KH.C1.423; KY21.Chr1.1715. The promoter drives the expression of GFP primarily in the notochord, in some cases in the mesenchyme and in larvae we also observed expression in a few neurons in the trunk. (A) Early tailbud embryos, (B) Late tailbud embryo, (C) Larva. See Supplementary Table 9 for breakdown of % of electroporated animals expressing GFP under either the *Ciinte*.*Brac* or *Ciinte*.*Cah* promoter.

**Figure S6.**
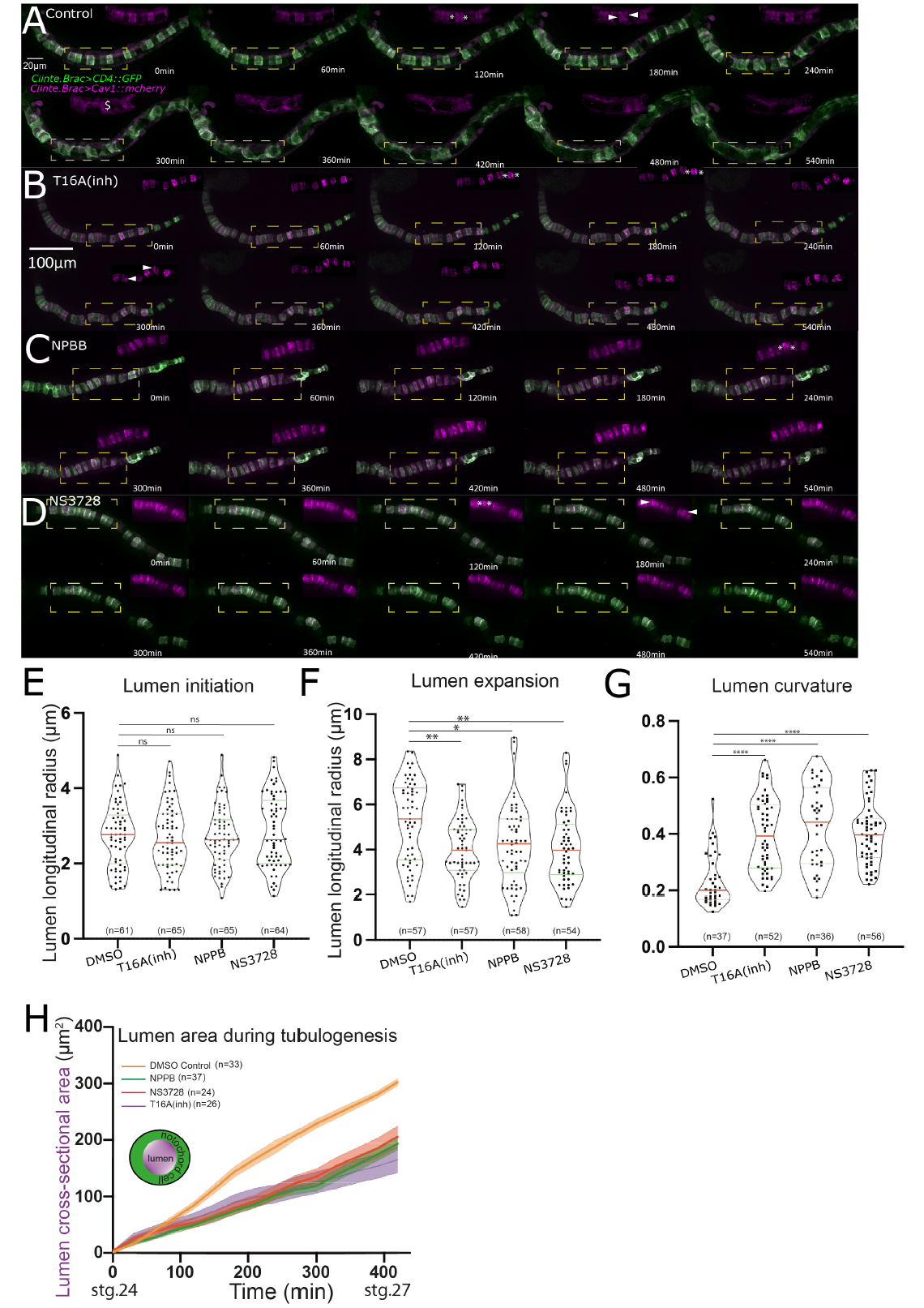
Drugs blocking Ano10 activity perturb notochord tubulogenesis. (A-D) Montage of time-lapse confocal movies from (A) DMSO control, (B) 100*μ*M T16A(inh)-treated, (C) 100*μ*M NPBB-treated and (D) 100*μ*M NS3728-treated embryos. Insets correspond to the regions marked by the yellow boxes. They show the developing lumen as demarcated by *Ciinte*.*Brac> Cav1::mCherry* (see Supplementary Movies 9-12). (E) Quantification of lumen longitudinal radius during lumen initiation in DMSO and drug treated embryos. (F) Quantification of lumen longitudinal radius during lumen expansion in DMSO and drug treated embryos in DMSO and drug treated embryos. (G) Quantification of lumen curvature after lumen connection. For statistical analysis of data shown in panels E-G we performed Kruskal-Wallis test followed by Dunn’s multiple comparisons test. In all panels individual data points are shown. The red and green lines correspond to the median and quartiles respectively. 0.05<ns,* <0.05, **<0.005 (H) Evolution of lumen cross-sectional area during tubulogenesis. Schematic shows in purple the lumen cross-section and in green the notochord cell membrane. For statistical analysis we performed a 2-way ANOVA followed by Tukey’s multiple comparison test. In all panels numbers shown indicate the number of animals analyzed in each condition.

## Notes

### Competing Interest Statement

The authors have declared no competing interest.

