## Supplementary Information for "Anoctamin 10/TMEM16K mediates convergent extension and tubulogenesis during notochord formation in the early chordate *Ciona intestinalis*"

Supplementary Table 1 Statistical analysis of data corresponding to panels in Figure 1.

| Figure & Panel | Test | Data point | Comparison | P value | P value summary |
|---|---|---|---|---|---|
| Fig. 1G | Two-way RM ANOVA | | | | |
| | | | Time | <0,0001 | **** |
| | | | Fraction intercalated | <0,0001 | **** |
| | | | Time x Fraction intercalated | <0,0001 | **** |
| | Tukey's multiple comparisons test | | Number of families<br>Number of comparisons per family | | 11<br>3 |
| | | 0 min | Negative Control vs. *Ano10 CRISPR* | >0,9999 | ns |
| | | | negative control vs. *Ano10 CRISPR; Ciinte.Brac>Ano10* | >0,9999 | ns |
| | | | *Ano10 CRISPR* vs.*Ano10 CRISPR; Ciinte.Brac>Ano10* | >0,9999 | ns |
| | | 30 min | Negative Control vs. *Ano10 CRISPR* | >0,9999 | ns |
| | | | negative control vs. *Ano10 CRISPR; Ciinte.Brac>Ano10* | 0,4716 | ns |
| | | | *Ano10 CRISPR* vs.*Ano10 CRISPR; Ciinte.Brac>Ano10* | 0,4646 | ns |
| | | 60 min | Negative Control vs. *Ano10 CRISPR* | <0,0001 | **** |
| | | | negative control vs. *Ano10 CRISPR; Ciinte.Brac>Ano10* | 0,8146 | ns |
| | | | *Ano10 CRISPR* vs.*Ano10 CRISPR; Ciinte.Brac>Ano10* | <0,0001 | **** |
| | | 90 min | Negative Control vs. *Ano10 CRISPR* | <0,0001 | **** |
| | | | negative control vs. *Ano10 CRISPR; Ciinte.Brac>Ano10* | 0,0818 | ns |
| | | | *Ano10 CRISPR* vs.*Ano10 CRISPR; Ciinte.Brac>Ano10* | <0,0001 | **** |
| | | 120 min | Negative Control vs. *Ano10 CRISPR* | <0,0001 | **** |
| | | | negative control vs. *Ano10 CRISPR; Ciinte.Brac>Ano10* | 0,007 | ** |
| | | | *Ano10 CRISPR* vs.*Ano10 CRISPR; Ciinte.Brac>Ano10* | <0,0001 | **** |
| | | 150 min | Negative Control vs. *Ano10 CRISPR* | <0,0001 | **** |
| | | | negative control vs. *Ano10 CRISPR; Ciinte.Brac>Ano10* | 0,133 | ns |
| | | | *Ano10 CRISPR* vs.*Ano10 CRISPR; Ciinte.Brac>Ano10* | <0,0001 | **** |
| | | 180 min | Negative Control vs. *Ano10 CRISPR* | <0,0001 | **** |
| | | | negative control vs. *Ano10 CRISPR; Ciinte.Brac>Ano10* | 0,5713 | ns |
| | | | *Ano10 CRISPR* vs.*Ano10 CRISPR; Ciinte.Brac>Ano10* | <0,0001 | **** |
| | | 210 min | Negative Control vs. *Ano10 CRISPR* | <0,0001 | **** |
| | | | negative control vs. *Ano10 CRISPR; Ciinte.Brac>Ano10* | 0,0001 | *** |
| | | | *Ano10 CRISPR* vs.*Ano10 CRISPR; Ciinte.Brac>Ano10* | <0,0001 | **** |
| | | 240 min | Negative Control vs. *Ano10 CRISPR* | <0,0001 | **** |
| | | | negative control vs. *Ano10 CRISPR; Ciinte.Brac>Ano10* | 0,0192 | * |
| | | | *Ano10 CRISPR* vs.*Ano10 CRISPR; Ciinte.Brac>Ano10* | <0,0001 | **** |

| | | | | | |
|---|---|---|---|---|---|
| | | 270 min | Negative Control vs. *Ano10 CRISPR* | <0,0001 | **** |
| | | | negative control vs. *Ano10 CRISPR; Ciinte.Brac>Ano10* | 0,0423 | * |
| | | | *Ano10 CRISPR* vs.*Ano10 CRISPR; Ciinte.Brac>Ano10* | <0,0001 | **** |
| | | 300 min | Negative Control vs. *Ano10 CRISPR* | <0,0001 | **** |
| | | | negative control vs. *Ano10 CRISPR; Ciinte.Brac>Ano10* | 0,2524 | ns |
| | | | *Ano10 CRISPR* vs.*Ano10 CRISPR; Ciinte.Brac>Ano10* | <0,0001 | **** |
| Fig. 1H | Mann–Whitney U test | Anterior 0 | Negative Control vs. *Ano10 CRISPR* | 0,124081620527574 | ns |
| | | | negative control vs. *Ano10 CRISPR; Ciinte.Brac>Ano10* | 1,70017429848954e-001 | ns |
| | | 1 | Negative Control vs. *Ano10 CRISPR* | 0,193344857119814 | ns |
| | | | negative control vs. *Ano10 CRISPR; Ciinte.Brac>Ano10* | 0,624553503707712 | ns |
| | | 2 | Negative Control vs. *Ano10 CRISPR* | 0,000257753331640547 | *** |
| | | | negative control vs. *Ano10 CRISPR; Ciinte.Brac>Ano10* | 0,870593452915895 | ns |
| | | 3 | Negative Control vs. *Ano10 CRISPR* | 0,00000119877532261048 | **** |
| | | | negative control vs. *Ano10 CRISPR; Ciinte.Brac>Ano10* | 0,808929661658417 | ns |
| | | 4 | Negative Control vs. *Ano10 CRISPR* | 0,00000854268499831308 | **** |
| | | | negative control vs. *Ano10 CRISPR; Ciinte.Brac>Ano10* | 0,799695342204138 | ns |
| | | 5 | Negative Control vs. *Ano10 CRISPR* | 6,91509019862523e-008 | **** |
| | | | negative control vs. *Ano10 CRISPR; Ciinte.Brac>Ano10* | 0,878684826099851 | ns |
| | | 6 | Negative Control vs. *Ano10 CRISPR* | 0,0000102166132606147 | **** |
| | | | negative control vs. *Ano10 CRISPR; Ciinte.Brac>Ano10* | 2,62854941231908e-001 | ns |
| | | 7 | Negative Control vs. *Ano10 CRISPR* | 0,000062702333667561 | **** |
| | | | negative control vs. *Ano10 CRISPR; Ciinte.Brac>Ano10* | 0,598715692779885 | ns |
| | | 8 | Negative Control vs. *Ano10 CRISPR* | 0,0000054419865572146 | **** |
| | | | negative control vs. *Ano10 CRISPR; Ciinte.Brac>Ano10* | 0,549015962570346 | ns |
| | | 9 | Negative Control vs. *Ano10 CRISPR* | 0,00211333013988449 | ** |
| | | | negative control vs. *Ano10 CRISPR; Ciinte.Brac>Ano10* | 0,938245936779039 | ns |
| | | 10 | Negative Control vs. *Ano10 CRISPR* | 0,0000329801865435444 | **** |
| | | | negative control vs. *Ano10 CRISPR; Ciinte.Brac>Ano10* | 0,622493263686162 | ns |
| | | 11 | Negative Control vs. *Ano10 CRISPR* | 0,000487109696181415 | *** |
| | | | negative control vs. *Ano10 CRISPR; Ciinte.Brac>Ano10* | 0,723307850613262 | ns |
| | | 12 | Negative Control vs. *Ano10 CRISPR* | 5,22831036619063e-007 | **** |
| | | | negative control vs. *Ano10 CRISPR; Ciinte.Brac>Ano10* | 0,760928763302164 | ns |
| | | 13 | Negative Control vs. *Ano10 CRISPR* | 2,28026880166123e-007 | **** |
| | | | negative control vs. *Ano10 CRISPR; Ciinte.Brac>Ano10* | 2,12865163356094e-001 | ns |

| | | | | | |
|---|---|---|---|---|---|
| | | 14 | Negative Control vs. *Ano10 CRISPR* | 0,00000314154127830318 | **** |
| | | | negative control vs. *Ano10 CRISPR; Ciinte.Brac>Ano10* | 2,88370569882536e-002 | * |
| | | 15 | Negative Control vs. *Ano10 CRISPR* | 0,00000325851262446773 | **** |
| | | | negative control vs. *Ano10 CRISPR; Ciinte.Brac>Ano10* | 2,95801129536866e-001 | ns |
| | | 16 | Negative Control vs. *Ano10 CRISPR* | 5,20518860926216e-007 | **** |
| | | | negative control vs. *Ano10 CRISPR; Ciinte.Brac>Ano10* | 4,31940555843752e-001 | ns |
| | | 17 | Negative Control vs. *Ano10 CRISPR* | 9,10788910360626e-008 | **** |
| | | | negative control vs. *Ano10 CRISPR; Ciinte.Brac>Ano10* | 1,09820115206328e-001 | ns |
| | | 18 | Negative Control vs. *Ano10 CRISPR* | 1,79820023075825e-007 | **** |
| | | | negative control vs. *Ano10 CRISPR; Ciinte.Brac>Ano10* | 9,43812407057159e-003 | ** |
| | | 19 | Negative Control vs. *Ano10 CRISPR* | 0,00000139268696654232 | **** |
| | | | negative control vs. *Ano10 CRISPR; Ciinte.Brac>Ano10* | 1,01740379438425e-002 | * |
| | | 20 | Negative Control vs. *Ano10 CRISPR* | 0,00000536326661651774 | **** |
| | | | negative control vs. *Ano10 CRISPR; Ciinte.Brac>Ano10* | 2,48901014078005e-002 | * |
| | | 21 | Negative Control vs. *Ano10 CRISPR* | 1,56323742892306e-007 | **** |
| | | | negative control vs. *Ano10 CRISPR; Ciinte.Brac>Ano10* | 9,77307296519096e-002 | ns |
| | | 22 | Negative Control vs. *Ano10 CRISPR* | 2,43423195031635e-008 | **** |
| | | | negative control vs. *Ano10 CRISPR; Ciinte.Brac>Ano10* | 1,10779026905533e-001 | ns |
| | | 23 | Negative Control vs. *Ano10 CRISPR* | 0,00000134001566773089 | **** |
| | | | negative control vs. *Ano10 CRISPR; Ciinte.Brac>Ano10* | 2,99513085665854e-002 | * |
| | | 24 | Negative Control vs. *Ano10 CRISPR* | 1,03536286813394e-009 | **** |
| | | | negative control vs. *Ano10 CRISPR; Ciinte.Brac>Ano10* | 4,50981090029217e-002 | * |
| | | 25 | Negative Control vs. *Ano10 CRISPR* | 2,1271911322026e-007 | **** |
| | | | negative control vs. *Ano10 CRISPR; Ciinte.Brac>Ano10* | 0,0296356891551094 | * |
| | | 26 | Negative Control vs. *Ano10 CRISPR* | 6,04211785059998e-007 | **** |
| | | | negative control vs. *Ano10 CRISPR; Ciinte.Brac>Ano10* | 4,74842337866955e-003 | ** |
| | | 27 | Negative Control vs. *Ano10 CRISPR* | 2,93136317649951e-008 | **** |
| | | | negative control vs. *Ano10 CRISPR; Ciinte.Brac>Ano10* | 4,37595746717589e-001 | ns |
| | | 28 | Negative Control vs. *Ano10 CRISPR* | 0,00000799075086437363 | **** |
| | | | negative control vs. *Ano10 CRISPR; Ciinte.Brac>Ano10* | 0,803500571370098 | ns |
| | | 29 | Negative Control vs. *Ano10 CRISPR* | 0,00000201929072415253 | **** |
| | | | negative control vs. *Ano10 CRISPR; Ciinte.Brac>Ano10* | 1,92649277965955e-001 | ns |
| | | 30 | Negative Control vs. *Ano10 CRISPR* | 0,0000012940086100217 | **** |
| | | | negative control vs. *Ano10 CRISPR; Ciinte.Brac>Ano10* | 1,69513601705789e-002 | * |
| | | 31 | Negative Control vs. *Ano10 CRISPR* | 0,000002400676925715 | **** |

| | | | | | |
|---|---|---|---|---|---|
| | | | negative control vs. *Ano10 CRISPR; Ciinte.Brac>Ano10* | 1,32277962899872e-002 | * |
| | | 32 | Negative Control vs. *Ano10 CRISPR* | 5,33851153443681e-008 | **** |
| | | | negative control vs. *Ano10 CRISPR; Ciinte.Brac>Ano10* | 3,68593238154156e-004 | *** |
| | | 33 | Negative Control vs. *Ano10 CRISPR* | 0,0000237459213251284 | **** |
| | | | negative control vs. *Ano10 CRISPR; Ciinte.Brac>Ano10* | 2,59462492142822e-002 | * |
| | | 34 | Negative Control vs. *Ano10 CRISPR* | 0,00000493809310694989 | **** |
| | | | negative control vs. *Ano10 CRISPR; Ciinte.Brac>Ano10* | 1,24544359091567e-002 | * |
| | | 35 | Negative Control vs. *Ano10 CRISPR* | 0,000023298386682022 | **** |
| | | | negative control vs. *Ano10 CRISPR; Ciinte.Brac>Ano10* | 2,63383264334484e-001 | ns |
| | | 36 | Negative Control vs. *Ano10 CRISPR* | 0,0000871043818400551 | **** |
| | | | negative control vs. *Ano10 CRISPR; Ciinte.Brac>Ano10* | 2,2926550081987e-002 | * |
| | | 37 | Negative Control vs. *Ano10 CRISPR* | 0,00261630897663883 | ** |
| | | | negative control vs. *Ano10 CRISPR; Ciinte.Brac>Ano10* | 0,539870417815097 | ns |
| | | 38 | Negative Control vs. *Ano10 CRISPR* | 0,0011791887058993 | ** |
| | | | negative control vs. *Ano10 CRISPR; Ciinte.Brac>Ano10* | 9,15809259550564e-002 | ns |
| | | 39 Posterior | Negative Control vs. *Ano10 CRISPR* | 0,00226828954846772 | ** |
| | | | negative control vs. *Ano10 CRISPR; Ciinte.Brac>Ano10* | 0,665568910158757 | ns |
| Fig. 1I | Mixed-effects model (REML) | | | | |
| | | | Time | <0,0001 | **** |
| | | | Notochord length | <0,0001 | **** |
| | | | Time x Notochord length | <0,0001 | **** |
| | Tukey's multiple comparisons test | | Number of families 11<br>Number of comparisons per family 3 | | |
| | | 0 min | Negative Control vs. *Ano10 CRISPR* | 0,8204 | ns |
| | | | negative control vs. *Ano10 CRISPR; Ciinte.Brac>Ano10* | 0,9661 | ns |
| | | | *Ano10 CRISPR* vs.*Ano10 CRISPR; Ciinte.Brac>Ano10* | 0,6349 | ns |
| | | 30 min | Negative Control vs. *Ano10 CRISPR* | 0,2126 | ns |
| | | | negative control vs. *Ano10 CRISPR; Ciinte.Brac>Ano10* | 0,7399 | ns |
| | | | *Ano10 CRISPR* vs.*Ano10 CRISPR; Ciinte.Brac>Ano10* | 0,4921 | ns |
| | | 60 min | Negative Control vs. *Ano10 CRISPR* | 0,049 | * |
| | | | negative control vs. *Ano10 CRISPR; Ciinte.Brac>Ano10* | 0,3338 | ns |
| | | | *Ano10 CRISPR* vs.*Ano10 CRISPR; Ciinte.Brac>Ano10* | 0,2059 | ns |

| | | | | | |
|---|---|---|---|---|---|
| | | 90 min | Negative Control vs. *Ano10 CRISPR* | 0,0277 | * |
| | | | negative control vs. *Ano10 CRISPR; Ciinte.Brac>Ano10* | 0,4505 | ns |
| | | | *Ano10 CRISPR* vs.*Ano10 CRISPR; Ciinte.Brac>Ano10* | 0,0388 | * |
| | | 120 min | Negative Control vs. *Ano10 CRISPR* | 0,0358 | * |
| | | | negative control vs. *Ano10 CRISPR; Ciinte.Brac>Ano10* | 0,7843 | ns |
| | | | *Ano10 CRISPR* vs.*Ano10 CRISPR; Ciinte.Brac>Ano10* | 0,0077 | ** |
| | | 150 min | Negative Control vs. *Ano10 CRISPR* | 0,0088 | ** |
| | | | negative control vs. *Ano10 CRISPR; Ciinte.Brac>Ano10* | 0,5481 | ns |
| | | | *Ano10 CRISPR* vs.*Ano10 CRISPR; Ciinte.Brac>Ano10* | 0,0043 | ** |
| | | 180 min | Negative Control vs. *Ano10 CRISPR* | 0,0022 | ** |
| | | | negative control vs. *Ano10 CRISPR; Ciinte.Brac>Ano10* | 0,4912 | ns |
| | | | *Ano10 CRISPR* vs.*Ano10 CRISPR; Ciinte.Brac>Ano10* | 0,0036 | ** |
| | | 210 min | Negative Control vs. *Ano10 CRISPR* | 0,0021 | ** |
| | | | negative control vs. *Ano10 CRISPR; Ciinte.Brac>Ano10* | 0,3456 | ns |
| | | | *Ano10 CRISPR* vs.*Ano10 CRISPR; Ciinte.Brac>Ano10* | 0,0018 | ** |
| | | 240 min | Negative Control vs. *Ano10 CRISPR* | 0,0005 | *** |
| | | | negative control vs. *Ano10 CRISPR; Ciinte.Brac>Ano10* | 0,5628 | ns |
| | | | *Ano10 CRISPR* vs.*Ano10 CRISPR; Ciinte.Brac>Ano10* | 0,0001 | *** |
| | | 270 min | Negative Control vs. *Ano10 CRISPR* | 0,0001 | *** |
| | | | negative control vs. *Ano10 CRISPR; Ciinte.Brac>Ano10* | 0,8486 | ns |
| | | | *Ano10 CRISPR* vs.*Ano10 CRISPR; Ciinte.Brac>Ano10* | <0.0001 | **** |
| | | 300 min | Negative Control vs. *Ano10 CRISPR* | <0.0001 | **** |
| | | | negative control vs. *Ano10 CRISPR; Ciinte.Brac>Ano10* | 0,8462 | ns |
| | | | *Ano10 CRISPR* vs.*Ano10 CRISPR; Ciinte.Brac>Ano10* | <0.0001 | **** |
| | | 330 min | Negative Control vs. *Ano10 CRISPR* | <0.0001 | **** |
| | | | negative control vs. *Ano10 CRISPR; Ciinte.Brac>Ano10* | 0,7836 | ns |
| | | | *Ano10 CRISPR* vs.*Ano10 CRISPR; Ciinte.Brac>Ano10* | <0.0001 | **** |
| | | 360 min | Negative Control vs. *Ano10 CRISPR* | <0.0001 | **** |
| | | | negative control vs. *Ano10 CRISPR; Ciinte.Brac>Ano10* | 0,8154 | ns |
| | | | *Ano10 CRISPR* vs.*Ano10 CRISPR; Ciinte.Brac>Ano10* | <0.0001 | **** |
| | | 390 min | Negative Control vs. *Ano10 CRISPR* | <0.0001 | **** |
| | | | negative control vs. *Ano10 CRISPR; Ciinte.Brac>Ano10* | 0,8193 | ns |
| | | | *Ano10 CRISPR* vs.*Ano10 CRISPR; Ciinte.Brac>Ano10* | <0.0001 | **** |
| | | 420 min | Negative Control vs. *Ano10 CRISPR* | <0.0001 | **** |
| | | | negative control vs. *Ano10 CRISPR; Ciinte.Brac>Ano10* | 0,6737 | ns |

| | | | | | |
|---|---|---|---|---|---|
| | | | *Ano10 CRISPR* vs.*Ano10 CRISPR; Ciinte.Brac>Ano10* | <0.0001 | **** |
| | | 450 min | Negative Control vs. *Ano10 CRISPR* | <0.0001 | **** |
| | | | negative control vs. *Ano10 CRISPR; Ciinte.Brac>Ano10* | 0,7517 | ns |
| | | | *Ano10 CRISPR* vs.*Ano10 CRISPR; Ciinte.Brac>Ano10* | <0.0001 | **** |
| | | 480 min | Negative Control vs. *Ano10 CRISPR* | <0.0001 | **** |
| | | | negative control vs. *Ano10 CRISPR; Ciinte.Brac>Ano10* | 0,4539 | ns |
| | | | *Ano10 CRISPR* vs.*Ano10 CRISPR; Ciinte.Brac>Ano10* | <0.0001 | **** |
| | | 510 min | Negative Control vs. *Ano10 CRISPR* | <0.0001 | **** |
| | | | negative control vs. *Ano10 CRISPR; Ciinte.Brac>Ano10* | 0,4998 | ns |
| | | | *Ano10 CRISPR* vs.*Ano10 CRISPR; Ciinte.Brac>Ano10* | <0.0001 | **** |
| | | 540 min | Negative Control vs. *Ano10 CRISPR* | <0.0001 | **** |
| | | | negative control vs. *Ano10 CRISPR; Ciinte.Brac>Ano10* | 0,407 | ns |
| | | | *Ano10 CRISPR* vs.*Ano10 CRISPR; Ciinte.Brac>Ano10* | <0.0001 | **** |
| | | 570 min | Negative Control vs. *Ano10 CRISPR* | <0.0001 | **** |
| | | | negative control vs. *Ano10 CRISPR; Ciinte.Brac>Ano10* | 0,366 | ns |
| | | | *Ano10 CRISPR* vs.*Ano10 CRISPR; Ciinte.Brac>Ano10* | <0.0001 | **** |
| | | 600 min | Negative Control vs. *Ano10 CRISPR* | <0.0001 | **** |
| | | | negative control vs. *Ano10 CRISPR; Ciinte.Brac>Ano10* | 0,2551 | ns |
| | | | *Ano10 CRISPR* vs.*Ano10 CRISPR; Ciinte.Brac>Ano10* | <0.0001 | **** |
| Fig. 1P | Kruskal-Wallis test | | All groups | <0,0001 | **** |
| | Dunn's multiple comparisons | | | | |
| | | | Negative Control vs. *Ano10 CRISPR* | <0,0001 | **** |
| | | | negative control vs. *Ano10 CRISPR; Ciinte.Brac>Ano10* | 0,1824 | ns |
| | | | *Ano10 CRISPR* vs.*Ano10 CRISPR; Ciinte.Brac>Ano10* | <0,0001 | **** |
| Fig. 1Q | Kruskal-Wallis test | | All groups | <0,0001 | **** |
| | Dunn's multiple comparisons | | | | |
| | | | Negative Control vs. *Ano10 CRISPR* | <0,0001 | **** |
| | | | negative control vs. *Ano10 CRISPR; Ciinte.Brac>Ano10* | 0,2223 | ns |
| | | | *Ano10 CRISPR* vs.*Ano10 CRISPR; Ciinte.Brac>Ano10* | <0,0001 | **** |

Supplementary Table 2 Statistical analysis of data corresponding to panels in Figure 2.

| Figure & Panel | Test | Data point | Comparison | P value | P value summary |
|---|---|---|---|---|---|
| Fig. 2E | Kruskal-Wallis test | | All groups | 0,9024 | ns |
| | Dunn's multiple comparisons | | | | |
| | | | Negative Control vs. *Ano10 CRISPR* | >0,9999 | ns |

| | | | | | |
|---|---|---|---|---|---|
| | | | negative control vs. *Ano10 CRISPR; Ciinte.Brac>Ano10* | >0,9999 | ns |
| | | | *Ano10 CRISPR* vs.*Ano10 CRISPR; Ciinte.Brac>Ano10* | >0,9999 | ns |
| Fig. 2G | Kruskal-Wallis test | | All groups | <0,0001 | **** |
| | Dunn's multiple comparisons | | Negative Control vs. *Ano10 CRISPR* | <0,0001 | **** |
| | | | negative control vs. *Ano10 CRISPR; Ciinte.Brac>Ano10* | 0,3678 | ns |
| | | | *Ano10 CRISPR* vs.*Ano10 CRISPR; Ciinte.Brac>Ano10* | <0,0001 | **** |
| Fig. 2H | Two-way RM ANOVA | | Time x Lumen cross-sectional area | <0,0001 | **** |
| | | | Time | <0,0001 | **** |
| | | | Lumen cross-sectional area | <0,0001 | **** |
| | | | Subject | <0,0001 | **** |
| | Tukey's multiple comparisons test | | Number of families<br>Number of comparisons per family | | 15<br>3 |
| | | 0 min | Negative Control vs. *Ano10 CRISPR* | 0,9664 | ns |
| | | | negative control vs. *Ano10 CRISPR; Ciinte.Brac>Ano10* | 0,7470 | ns |
| | | | *Ano10 CRISPR* vs.*Ano10 CRISPR; Ciinte.Brac>Ano10* | 0,8533 | ns |
| | | 30 min | Negative Control vs. *Ano10 CRISPR* | 0,9648 | ns |
| | | | negative control vs. *Ano10 CRISPR; Ciinte.Brac>Ano10* | 0,9974 | ns |
| | | | *Ano10 CRISPR* vs.*Ano10 CRISPR; Ciinte.Brac>Ano10* | 0,9637 | ns |
| | | 60 min | Negative Control vs. *Ano10 CRISPR* | 0,9648 | ns |
| | | | negative control vs. *Ano10 CRISPR; Ciinte.Brac>Ano10* | 0,9974 | ns |
| | | | *Ano10 CRISPR* vs.*Ano10 CRISPR; Ciinte.Brac>Ano10* | 0,9637 | ns |
| | | 90 min | Negative Control vs. *Ano10 CRISPR* | 0,0073 | ** |
| | | | negative control vs. *Ano10 CRISPR; Ciinte.Brac>Ano10* | 0,2816 | ns |
| | | | *Ano10 CRISPR* vs.*Ano10 CRISPR; Ciinte.Brac>Ano10* | 0,5573 | ns |
| | | 120 min | Negative Control vs. *Ano10 CRISPR* | 0,3287 | ns |
| | | | negative control vs. *Ano10 CRISPR; Ciinte.Brac>Ano10* | 0,1538 | ns |
| | | | *Ano10 CRISPR* vs.*Ano10 CRISPR; Ciinte.Brac>Ano10* | 0,8767 | ns |
| | | 150 min | Negative Control vs. *Ano10 CRISPR* | 0,1171 | ns |
| | | | negative control vs. *Ano10 CRISPR; Ciinte.Brac>Ano10* | 0,4682 | ns |
| | | | *Ano10 CRISPR* vs.*Ano10 CRISPR; Ciinte.Brac>Ano10* | 0,5818 | ns |
| | | 180 min | Negative Control vs. *Ano10 CRISPR* | 0,0091 | ** |
| | | | negative control vs. *Ano10 CRISPR; Ciinte.Brac>Ano10* | 0,9603 | ns |
| | | | *Ano10 CRISPR* vs.*Ano10 CRISPR; Ciinte.Brac>Ano10* | 0,019 | * |
| | | 210 min | Negative Control vs. *Ano10 CRISPR* | 0,002 | ** |

| | | | | | |
|---|---|---|---|---|---|
| | | | negative control vs. *Ano10 CRISPR; Ciinte.Brac>Ano10* | 0,9629 | ns |
| | | | *Ano10 CRISPR* vs.*Ano10 CRISPR; Ciinte.Brac>Ano10* | 0,0091 | ** |
| | | 240 min | Negative Control vs. *Ano10 CRISPR* | <0,0001 | **** |
| | | | negative control vs. *Ano10 CRISPR; Ciinte.Brac>Ano10* | 0,9859 | ns |
| | | | *Ano10 CRISPR* vs.*Ano10 CRISPR; Ciinte.Brac>Ano10* | 0,0016 | ** |
| | | 270 min | Negative Control vs. *Ano10 CRISPR* | <0,0001 | **** |
| | | | negative control vs. *Ano10 CRISPR; Ciinte.Brac>Ano10* | 0,8999 | ns |
| | | | *Ano10 CRISPR* vs.*Ano10 CRISPR; Ciinte.Brac>Ano10* | 0,0006 | *** |
| | | 300 min | Negative Control vs. *Ano10 CRISPR* | <0,0001 | **** |
| | | | negative control vs. *Ano10 CRISPR; Ciinte.Brac>Ano10* | 0,7134 | ns |
| | | | *Ano10 CRISPR* vs.*Ano10 CRISPR; Ciinte.Brac>Ano10* | <0,0001 | **** |
| | | 330 min | Negative Control vs. *Ano10 CRISPR* | <0,0001 | **** |
| | | | negative control vs. *Ano10 CRISPR; Ciinte.Brac>Ano10* | 0,7579 | ns |
| | | | *Ano10 CRISPR* vs.*Ano10 CRISPR; Ciinte.Brac>Ano10* | <0,0001 | **** |
| | | 360 min | Negative Control vs. *Ano10 CRISPR* | <0,0001 | **** |
| | | | negative control vs. *Ano10 CRISPR; Ciinte.Brac>Ano10* | 0,9355 | ns |
| | | | *Ano10 CRISPR* vs.*Ano10 CRISPR; Ciinte.Brac>Ano10* | <0,0001 | **** |
| | | 390 min | Negative Control vs. *Ano10 CRISPR* | <0,0001 | **** |
| | | | negative control vs. *Ano10 CRISPR; Ciinte.Brac>Ano10* | 0,9813 | ns |
| | | | *Ano10 CRISPR* vs.*Ano10 CRISPR; Ciinte.Brac>Ano10* | <0,0001 | **** |
| | | 420 min | Negative Control vs. *Ano10 CRISPR* | <0,0001 | **** |
| | | | negative control vs. *Ano10 CRISPR; Ciinte.Brac>Ano10* | >0,9999 | ns |
| | | | *Ano10 CRISPR* vs.*Ano10 CRISPR; Ciinte.Brac>Ano10* | 0,0003 | *** |

Supplementary Table 3 Statistical analysis of data corresponding to panels in Figure 3.

| Figure & | Test | Data point | Comparison | P value | P value summary |
|---|---|---|---|---|---|
| Fig. 3G | Kruskal-Wallis test | | All groups | <0,0001 | **** |
| | Dunn's multiple comparisons | | | | |
| | | | Negative Control vs. *Ano10 CRISPR* | <0,0001 | **** |
| | | | negative control vs. *Ano10 CRISPR; Ciinte.Brac>Ano10* | >0,9999 | ns |
| | | | *Ano10 CRISPR* vs.*Ano10 CRISPR; Ciinte.Brac>Ano10* | <0,0001 | **** |
| Fig. 3H | Kruskal-Wallis test | | All groups | <0,0001 | **** |
| | Dunn's multiple comparisons | | Negative Control vs. *Ano10 CRISPR* | <0,0001 | **** |
| | | | negative control vs. *Ano10 CRISPR; Ciinte.Brac>Ano10* | >0,9999 | ns |
| | | | *Ano10 CRISPR* vs.*Ano10 CRISPR; Ciinte.Brac>Ano10* | <0,0001 | **** |

| | | | | | |
|---|---|---|---|---|---|
| Fig. 3I | Mixed-effects model (REML) | | Time | <0,0001 | **** |
| | | | Speed | 0,0011 | ** |
| | Tukey's multiple comparisons | | Number of families | | 125 |
| | | | Number of comparisons per family | | 3 |
| | | 5 min | Negative Control vs. *Ano10 CRISPR* | 0,0045 | ** |
| | | | negative control vs. *Ano10 CRISPR; Ciinte.Brac>Ano10* | 0,0219 | * |
| | | | *Ano10 CRISPR* vs.*Ano10 CRISPR; Ciinte.Brac>Ano10* | 0,9009 | ns |
| | | 10 min | Negative Control vs. *Ano10 CRISPR* | 0,0169 | * |
| | | | negative control vs. *Ano10 CRISPR; Ciinte.Brac>Ano10* | 0,3122 | ns |
| | | | *Ano10 CRISPR* vs.*Ano10 CRISPR; Ciinte.Brac>Ano10* | 0,5361 | ns |
| | | 85min | Negative Control vs. *Ano10 CRISPR* | 0,3289 | ns |
| | | | negative control vs. *Ano10 CRISPR; Ciinte.Brac>Ano10* | 0,3989 | ns |
| | | | *Ano10 CRISPR* vs.*Ano10 CRISPR; Ciinte.Brac>Ano10* | 0,0479 | * |
| | | 90 min | Negative Control vs. *Ano10 CRISPR* | 0,2433 | ns |
| | | | negative control vs. *Ano10 CRISPR; Ciinte.Brac>Ano10* | 0,3793 | ns |
| | | | *Ano10 CRISPR* vs.*Ano10 CRISPR; Ciinte.Brac>Ano10* | 0,0256 | * |
| | | 95 min | Negative Control vs. *Ano10 CRISPR* | 0,1703 | ns |
| | | | negative control vs. *Ano10 CRISPR; Ciinte.Brac>Ano10* | 0,4031 | ns |
| | | | *Ano10 CRISPR* vs.*Ano10 CRISPR; Ciinte.Brac>Ano10* | 0,0142 | * |
| | | 100 min | Negative Control vs. *Ano10 CRISPR* | 0,1964 | ns |
| | | | negative control vs. *Ano10 CRISPR; Ciinte.Brac>Ano10* | 0,4561 | ns |
| | | | *Ano10 CRISPR* vs.*Ano10 CRISPR; Ciinte.Brac>Ano10* | 0,0177 | * |
| | | 105 min | Negative Control vs. *Ano10 CRISPR* | 0,1643 | ns |
| | | | negative control vs. *Ano10 CRISPR; Ciinte.Brac>Ano10* | 0,5159 | ns |
| | | | *Ano10 CRISPR* vs.*Ano10 CRISPR; Ciinte.Brac>Ano10* | 0,0121 | * |
| | | 110 min | Negative Control vs. *Ano10 CRISPR* | 0,1181 | ns |
| | | | negative control vs. *Ano10 CRISPR; Ciinte.Brac>Ano10* | 0,6733 | ns |
| | | | *Ano10 CRISPR* vs.*Ano10 CRISPR; Ciinte.Brac>Ano10* | 0,0106 | * |
| | | 115 min | Negative Control vs. *Ano10 CRISPR* | 0,0762 | ns |
| | | | negative control vs. *Ano10 CRISPR; Ciinte.Brac>Ano10* | 0,838 | ns |
| | | | *Ano10 CRISPR* vs.*Ano10 CRISPR; Ciinte.Brac>Ano10* | 0,0106 | * |
| | | 120 min | Negative Control vs. *Ano10 CRISPR* | 0,05 | ns |
| | | | negative control vs. *Ano10 CRISPR; Ciinte.Brac>Ano10* | 0,967 | ns |
| | | | *Ano10 CRISPR* vs.*Ano10 CRISPR; Ciinte.Brac>Ano10* | 0,0111 | * |

| | | | | | |
|---|---|---|---|---|---|
| | | 125 min | Negative Control vs. *Ano10 CRISPR* | 0,0402 | * |
| | | | negative control vs. *Ano10 CRISPR; Ciinte.Brac>Ano10* | 0,9982 | ns |
| | | | *Ano10 CRISPR* vs.*Ano10 CRISPR; Ciinte.Brac>Ano10* | 0,0135 | * |
| | | 130 min | Negative Control vs. *Ano10 CRISPR* | 0,0338 | * |
| | | | negative control vs. *Ano10 CRISPR; Ciinte.Brac>Ano10* | 0,9497 | ns |
| | | | *Ano10 CRISPR* vs.*Ano10 CRISPR; Ciinte.Brac>Ano10* | 0,0168 | * |
| | | 135 min | Negative Control vs. *Ano10 CRISPR* | 0,0313 | * |
| | | | negative control vs. *Ano10 CRISPR; Ciinte.Brac>Ano10* | 0,9074 | ns |
| | | | *Ano10 CRISPR* vs.*Ano10 CRISPR; Ciinte.Brac>Ano10* | 0,0142 | * |
| | | 140 min | Negative Control vs. *Ano10 CRISPR* | 0,0351 | * |
| | | | negative control vs. *Ano10 CRISPR; Ciinte.Brac>Ano10* | 0,8747 | ns |
| | | | *Ano10 CRISPR* vs.*Ano10 CRISPR; Ciinte.Brac>Ano10* | 0,0165 | * |
| | | 145 min | Negative Control vs. *Ano10 CRISPR* | 0,0343 | * |
| | | | negative control vs. *Ano10 CRISPR; Ciinte.Brac>Ano10* | 0,8589 | ns |
| | | | *Ano10 CRISPR* vs.*Ano10 CRISPR; Ciinte.Brac>Ano10* | 0,016 | * |
| | | 150 min | Negative Control vs. *Ano10 CRISPR* | 0,0376 | * |
| | | | negative control vs. *Ano10 CRISPR; Ciinte.Brac>Ano10* | 0,8613 | ns |
| | | | *Ano10 CRISPR* vs.*Ano10 CRISPR; Ciinte.Brac>Ano10* | 0,0193 | * |
| | | 155 min | Negative Control vs. *Ano10 CRISPR* | 0,0527 | ns |
| | | | negative control vs. *Ano10 CRISPR; Ciinte.Brac>Ano10* | 0,8931 | ns |
| | | | *Ano10 CRISPR* vs.*Ano10 CRISPR; Ciinte.Brac>Ano10* | 0,0283 | * |
| | | 160 min | Negative Control vs. *Ano10 CRISPR* | 0,0527 | ns |
| | | | negative control vs. *Ano10 CRISPR; Ciinte.Brac>Ano10* | 0,8931 | ns |
| | | | *Ano10 CRISPR* vs.*Ano10 CRISPR; Ciinte.Brac>Ano10* | 0,0283 | * |
| | | 165 min | Negative Control vs. *Ano10 CRISPR* | 0,0854 | ns |
| | | | negative control vs. *Ano10 CRISPR; Ciinte.Brac>Ano10* | 0,952 | ns |
| | | | *Ano10 CRISPR* vs.*Ano10 CRISPR; Ciinte.Brac>Ano10* | 0,0481 | * |
| | | 225 min | Negative Control vs. *Ano10 CRISPR* | 0,2064 | ns |
| | | | negative control vs. *Ano10 CRISPR; Ciinte.Brac>Ano10* | 0,9314 | ns |
| | | | *Ano10 CRISPR* vs.*Ano10 CRISPR; Ciinte.Brac>Ano10* | 0,0381 | * |
| | | 230 min | Negative Control vs. *Ano10 CRISPR* | 0,1667 | ns |
| | | | negative control vs. *Ano10 CRISPR; Ciinte.Brac>Ano10* | 0,9421 | ns |
| | | | *Ano10 CRISPR* vs.*Ano10 CRISPR; Ciinte.Brac>Ano10* | 0,0371 | * |
| | | 235 min | Negative Control vs. *Ano10 CRISPR* | 0,1071 | ns |
| | | | negative control vs. *Ano10 CRISPR; Ciinte.Brac>Ano10* | 0,9805 | ns |

| | | | | | |
|---|---|---|---|---|---|
| | | | *Ano10 CRISPR* vs.*Ano10 CRISPR; Ciinte.Brac>Ano10* | 0,0421 | * |
| | | 240 min | Negative Control vs. *Ano10 CRISPR* | 0,0781 | ns |
| | | | negative control vs. *Ano10 CRISPR; Ciinte.Brac>Ano10* | 0,9941 | ns |
| | | | *Ano10 CRISPR* vs.*Ano10 CRISPR; Ciinte.Brac>Ano10* | 0,0406 | * |
| | | 245 min | Negative Control vs. *Ano10 CRISPR* | 0,0594 | ns |
| | | | negative control vs. *Ano10 CRISPR; Ciinte.Brac>Ano10* | 0,9993 | ns |
| | | | *Ano10 CRISPR* vs.*Ano10 CRISPR; Ciinte.Brac>Ano10* | 0,035 | * |
| | | 250 min | Negative Control vs. *Ano10 CRISPR* | 0,0479 | * |
| | | | negative control vs. *Ano10 CRISPR; Ciinte.Brac>Ano10* | 0,9989 | ns |
| | | | *Ano10 CRISPR* vs.*Ano10 CRISPR; Ciinte.Brac>Ano10* | 0,0316 | * |
| | | 255 min | Negative Control vs. *Ano10 CRISPR* | 0,0439 | * |
| | | | negative control vs. *Ano10 CRISPR; Ciinte.Brac>Ano10* | 0,988 | ns |
| | | | *Ano10 CRISPR* vs.*Ano10 CRISPR; Ciinte.Brac>Ano10* | 0,0364 | * |
| | | 260 min | Negative Control vs. *Ano10 CRISPR* | 0,0474 | * |
| | | | negative control vs. *Ano10 CRISPR; Ciinte.Brac>Ano10* | 0,9705 | ns |
| | | | *Ano10 CRISPR* vs.*Ano10 CRISPR; Ciinte.Brac>Ano10* | 0,0494 | * |
| | | 270 min | Negative Control vs. *Ano10 CRISPR* | 0,0473 | * |
| | | | negative control vs. *Ano10 CRISPR; Ciinte.Brac>Ano10* | 0,9538 | ns |
| | | | *Ano10 CRISPR* vs.*Ano10 CRISPR; Ciinte.Brac>Ano10* | 0,0616 | ns |
| | | 275 min | Negative Control vs. *Ano10 CRISPR* | 0,0478 | * |
| | | | negative control vs. *Ano10 CRISPR; Ciinte.Brac>Ano10* | 0,9617 | ns |
| | | | *Ano10 CRISPR* vs.*Ano10 CRISPR; Ciinte.Brac>Ano10* | 0,0627 | ns |
| | | 280 min | Negative Control vs. *Ano10 CRISPR* | 0,0496 | * |
| | | | negative control vs. *Ano10 CRISPR; Ciinte.Brac>Ano10* | 0,98 | ns |
| | | | *Ano10 CRISPR* vs.*Ano10 CRISPR; Ciinte.Brac>Ano10* | 0,0547 | ns |
| | | 285 min | Negative Control vs. *Ano10 CRISPR* | 0,0484 | * |
| | | | negative control vs. *Ano10 CRISPR; Ciinte.Brac>Ano10* | 0,9915 | ns |
| | | | *Ano10 CRISPR* vs.*Ano10 CRISPR; Ciinte.Brac>Ano10* | 0,0465 | * |
| | | 290 min | Negative Control vs. *Ano10 CRISPR* | 0,0396 | * |
| | | | negative control vs. *Ano10 CRISPR; Ciinte.Brac>Ano10* | 0,9928 | ns |
| | | | *Ano10 CRISPR* vs.*Ano10 CRISPR; Ciinte.Brac>Ano10* | 0,0453 | * |
| | | 295 min | Negative Control vs. *Ano10 CRISPR* | 0,0369 | * |
| | | | negative control vs. *Ano10 CRISPR; Ciinte.Brac>Ano10* | 0,9981 | ns |
| | | | *Ano10 CRISPR* vs.*Ano10 CRISPR; Ciinte.Brac>Ano10* | 0,0425 | * |
| | | 300 min | Negative Control vs. *Ano10 CRISPR* | 0,0397 | * |

| | | | | | |
|---|---|---|---|---|---|
| | | | negative control vs. *Ano10 CRISPR; Ciinte.Brac>Ano10* | >0,9999 | ns |
| | | | *Ano10 CRISPR* vs.*Ano10 CRISPR; Ciinte.Brac>Ano10* | 0,0478 | * |
| | | 365 min | Negative Control vs. *Ano10 CRISPR* | 0,2455 | ns |
| | | | negative control vs. *Ano10 CRISPR; Ciinte.Brac>Ano10* | 0,718 | ns |
| | | | *Ano10 CRISPR* vs.*Ano10 CRISPR; Ciinte.Brac>Ano10* | 0,0484 | * |
| | | 370 min | Negative Control vs. *Ano10 CRISPR* | 0,2801 | ns |
| | | | negative control vs. *Ano10 CRISPR; Ciinte.Brac>Ano10* | 0,6578 | ns |
| | | | *Ano10 CRISPR* vs.*Ano10 CRISPR; Ciinte.Brac>Ano10* | 0,0401 | * |
| | | 375 min | Negative Control vs. *Ano10 CRISPR* | 0,3603 | ns |
| | | | negative control vs. *Ano10 CRISPR; Ciinte.Brac>Ano10* | 0,5709 | ns |
| | | | *Ano10 CRISPR* vs.*Ano10 CRISPR; Ciinte.Brac>Ano10* | 0,0379 | * |
| | | 380 min | Negative Control vs. *Ano10 CRISPR* | 0,464 | ns |
| | | | negative control vs. *Ano10 CRISPR; Ciinte.Brac>Ano10* | 0,5019 | ns |
| | | | *Ano10 CRISPR* vs.*Ano10 CRISPR; Ciinte.Brac>Ano10* | 0,0423 | * |
| Fig. 3M | Kruskal-Wallis test | | *All groups* | <0,0001 | **** |
| | Dunn's multiple comparisons test | | Negative Control vs. *Ano10 CRISPR* | 0,0009 | *** |
| | | | negative control vs. *Ano10 CRISPR; Ciinte.Brac>Ano10* | 0,7282 | ns |
| | | | *Ano10 CRISPR* vs.*Ano10 CRISPR; Ciinte.Brac>Ano10* | <0,0001 | **** |
| Fig. 3N | Kruskal-Wallis test | | *All groups* | 0,0011 | ** |
| | Dunn's multiple comparisons | | Negative Control vs. *Ano10 CRISPR* | 0,0138 | * |
| | | | negative control vs. *Ano10 CRISPR; Ciinte.Brac>Ano10* | >0,9999 | ns |
| | | | *Ano10 CRISPR* vs.*Ano10 CRISPR; Ciinte.Brac>Ano10* | 0,002 | ** |
| Fig. 3S | Kruskal-Wallis test | | *All groups* | <0,0001 | **** |
| | Dunn's multiple comparisons | | Negative Control vs. *Ano10 CRISPR* | 0,0005 | *** |
| | | | negative control vs. *Ano10 CRISPR; Ciinte.Brac>Ano10* | 0,0335 | * |
| | | | *Ano10 CRISPR* vs.*Ano10 CRISPR; Ciinte.Brac>Ano10* | <0,0001 | **** |
| Fig.3X | Mixed-effects model (REML) | | *A-P Distance* | <0,0001 | **** |
| | | | Normalized intesity | 0,0049 | ** |
| | Tukey's multiple comparisons test | | Number of families<br>Number of comparisons per family | 99<br>3 | |

| | | | | | |
|---|---|---|---|---|---|
| | | 1,625µm | Negative Control vs. *Ano10 CRISPR* | 0,0257 | * |
| | | | negative control vs. *Ano10 CRISPR; Ciinte.Brac>Ano10* | 0,4496 | ns |
| | | | *Ano10 CRISPR* vs.*Ano10 CRISPR; Ciinte.Brac>Ano10* | 0,3041 | ns |
| | | 1,7875µm | Negative Control vs. *Ano10 CRISPR* | 0,0071 | ** |
| | | | negative control vs. *Ano10 CRISPR; Ciinte.Brac>Ano10* | 0,5834 | ns |
| | | | *Ano10 CRISPR* vs.*Ano10 CRISPR; Ciinte.Brac>Ano10* | 0,0802 | ns |
| | | 1,95µm | Negative Control vs. *Ano10 CRISPR* | 0,0025 | ** |
| | | | negative control vs. *Ano10 CRISPR; Ciinte.Brac>Ano10* | 0,7516 | ns |
| | | | *Ano10 CRISPR* vs.*Ano10 CRISPR; Ciinte.Brac>Ano10* | 0,015 | * |
| | | 2,1125µm | Negative Control vs. *Ano10 CRISPR* | 0,0015 | ** |
| | | | negative control vs. *Ano10 CRISPR; Ciinte.Brac>Ano10* | 0,8969 | ns |
| | | | *Ano10 CRISPR* vs.*Ano10 CRISPR; Ciinte.Brac>Ano10* | 0,0024 | ** |
| | | 2,275µm | Negative Control vs. *Ano10 CRISPR* | 0,0003 | *** |
| | | | negative control vs. *Ano10 CRISPR; Ciinte.Brac>Ano10* | 0,9977 | ns |
| | | | *Ano10 CRISPR* vs.*Ano10 CRISPR; Ciinte.Brac>Ano10* | <0,0001 | **** |
| | | 2,4375µm | Negative Control vs. *Ano10 CRISPR* | <0,0001 | **** |
| | | | negative control vs. *Ano10 CRISPR; Ciinte.Brac>Ano10* | 0,9834 | ns |
| | | | *Ano10 CRISPR* vs.*Ano10 CRISPR; Ciinte.Brac>Ano10* | <0,0001 | **** |
| | | 2,6µm | Negative Control vs. *Ano10 CRISPR* | <0,0001 | **** |
| | | | negative control vs. *Ano10 CRISPR; Ciinte.Brac>Ano10* | 0,9781 | ns |
| | | | *Ano10 CRISPR* vs.*Ano10 CRISPR; Ciinte.Brac>Ano10* | <0,0001 | **** |
| | | 2,7625µm | Negative Control vs. *Ano10 CRISPR* | <0,0001 | **** |
| | | | negative control vs. *Ano10 CRISPR; Ciinte.Brac>Ano10* | 0,9953 | ns |
| | | | *Ano10 CRISPR* vs.*Ano10 CRISPR; Ciinte.Brac>Ano10* | <0,0001 | **** |
| | | 2,925µm | Negative Control vs. *Ano10 CRISPR* | <0,0001 | **** |
| | | | negative control vs. *Ano10 CRISPR; Ciinte.Brac>Ano10* | 0,9981 | ns |
| | | | *Ano10 CRISPR* vs.*Ano10 CRISPR; Ciinte.Brac>Ano10* | <0,0001 | **** |
| | | 3,0875µm | Negative Control vs. *Ano10 CRISPR* | <0,0001 | **** |
| | | | negative control vs. *Ano10 CRISPR; Ciinte.Brac>Ano10* | 0,9179 | ns |
| | | | *Ano10 CRISPR* vs.*Ano10 CRISPR; Ciinte.Brac>Ano10* | <0,0001 | **** |
| | | 3,25µm | Negative Control vs. *Ano10 CRISPR* | <0,0001 | **** |
| | | | negative control vs. *Ano10 CRISPR; Ciinte.Brac>Ano10* | 0,8621 | ns |
| | | | *Ano10 CRISPR* vs.*Ano10 CRISPR; Ciinte.Brac>Ano10* | <0,0001 | **** |
| | | 3,4125µm | Negative Control vs. *Ano10 CRISPR* | <0,0001 | **** |
| | | | negative control vs. *Ano10 CRISPR; Ciinte.Brac>Ano10* | 0,8698 | ns |

| | | | | | |
|---|---|---|---|---|---|
| | | | *Ano10 CRISPR* vs.*Ano10 CRISPR; Ciinte.Brac>Ano10* | <0,0001 | **** |
| | | 3,575μm | Negative Control vs. *Ano10 CRISPR* | <0,0001 | **** |
| | | | negative control vs. *Ano10 CRISPR; Ciinte.Brac>Ano10* | 0,8007 | ns |
| | | | *Ano10 CRISPR* vs.*Ano10 CRISPR; Ciinte.Brac>Ano10* | <0,0001 | **** |
| | | 3,7375μm | Negative Control vs. *Ano10 CRISPR* | <0,0001 | **** |
| | | | negative control vs. *Ano10 CRISPR; Ciinte.Brac>Ano10* | 0,6454 | ns |
| | | | *Ano10 CRISPR* vs.*Ano10 CRISPR; Ciinte.Brac>Ano10* | 0,0003 | *** |
| | | 3,9μm | Negative Control vs. *Ano10 CRISPR* | <0,0001 | **** |
| | | | negative control vs. *Ano10 CRISPR; Ciinte.Brac>Ano10* | 0,5408 | ns |
| | | | *Ano10 CRISPR* vs.*Ano10 CRISPR; Ciinte.Brac>Ano10* | 0,0041 | ** |
| | | 4,0625μm | Negative Control vs. *Ano10 CRISPR* | 0,0002 | *** |
| | | | negative control vs. *Ano10 CRISPR; Ciinte.Brac>Ano10* | 0,3631 | ns |
| | | | *Ano10 CRISPR* vs.*Ano10 CRISPR; Ciinte.Brac>Ano10* | 0,0157 | * |
| | | 4,225μm | Negative Control vs. *Ano10 CRISPR* | 0,0004 | *** |
| | | | negative control vs. *Ano10 CRISPR; Ciinte.Brac>Ano10* | 0,2538 | ns |
| | | | *Ano10 CRISPR* vs.*Ano10 CRISPR; Ciinte.Brac>Ano10* | 0,0393 | * |
| | | 4,3875μm | Negative Control vs. *Ano10 CRISPR* | 0,0014 | ** |
| | | | negative control vs. *Ano10 CRISPR; Ciinte.Brac>Ano10* | 0,1936 | ns |
| | | | *Ano10 CRISPR* vs.*Ano10 CRISPR; Ciinte.Brac>Ano10* | 0,1243 | ns |
| | | 4,55μm | Negative Control vs. *Ano10 CRISPR* | 0,0044 | ** |
| | | | negative control vs. *Ano10 CRISPR; Ciinte.Brac>Ano10* | 0,1523 | ns |
| | | | *Ano10 CRISPR* vs.*Ano10 CRISPR; Ciinte.Brac>Ano10* | 0,3159 | ns |
| | | 4,7125μm | Negative Control vs. *Ano10 CRISPR* | 0,02 | * |
| | | | negative control vs. *Ano10 CRISPR; Ciinte.Brac>Ano10* | 0,1357 | ns |
| | | | *Ano10 CRISPR* vs.*Ano10 CRISPR; Ciinte.Brac>Ano10* | 0,6124 | ns |
| | | 6,175μm | Negative Control vs. *Ano10 CRISPR* | 0,8446 | ns |
| | | | negative control vs. *Ano10 CRISPR; Ciinte.Brac>Ano10* | 0,1309 | ns |
| | | | *Ano10 CRISPR* vs.*Ano10 CRISPR; Ciinte.Brac>Ano10* | 0,033 | * |
| | | 6,3375μm | Negative Control vs. *Ano10 CRISPR* | 0,6529 | ns |
| | | | negative control vs. *Ano10 CRISPR; Ciinte.Brac>Ano10* | 0,2251 | ns |
| | | | *Ano10 CRISPR* vs.*Ano10 CRISPR; Ciinte.Brac>Ano10* | 0,0295 | * |
| | | 6,5μm | Negative Control vs. *Ano10 CRISPR* | 0,3649 | ns |
| | | | negative control vs. *Ano10 CRISPR; Ciinte.Brac>Ano10* | 0,3685 | ns |
| | | | *Ano10 CRISPR* vs.*Ano10 CRISPR; Ciinte.Brac>Ano10* | 0,0196 | * |
| | | 6,6625μm | Negative Control vs. *Ano10 CRISPR* | 0,1676 | ns |

| | | | | | |
|---|---|---|---|---|---|
| | | | negative control vs. *Ano10 CRISPR; Ciinte.Brac>Ano10* | 0,4746 | ns |
| | | | *Ano10 CRISPR* vs.*Ano10 CRISPR; Ciinte.Brac>Ano10* | 0,0101 | * |
| | | 6,825μm | Negative Control vs. *Ano10 CRISPR* | 0,0945 | ns |
| | | | negative control vs. *Ano10 CRISPR; Ciinte.Brac>Ano10* | 0,4716 | ns |
| | | | *Ano10 CRISPR* vs.*Ano10 CRISPR; Ciinte.Brac>Ano10* | 0,0041 | ** |
| | | 6,9875μm | Negative Control vs. *Ano10 CRISPR* | 0,0658 | ns |
| | | | negative control vs. *Ano10 CRISPR; Ciinte.Brac>Ano10* | 0,4066 | ns |
| | | | *Ano10 CRISPR* vs.*Ano10 CRISPR; Ciinte.Brac>Ano10* | 0,0013 | ** |
| | | 7,15μm | Negative Control vs. *Ano10 CRISPR* | 0,0321 | * |
| | | | negative control vs. *Ano10 CRISPR; Ciinte.Brac>Ano10* | 0,3818 | ns |
| | | | *Ano10 CRISPR* vs.*Ano10 CRISPR; Ciinte.Brac>Ano10* | 0,0004 | *** |
| | | 7,3125μm | Negative Control vs. *Ano10 CRISPR* | 0,0127 | * |
| | | | negative control vs. *Ano10 CRISPR; Ciinte.Brac>Ano10* | 0,4357 | ns |
| | | | *Ano10 CRISPR* vs.*Ano10 CRISPR; Ciinte.Brac>Ano10* | 0,0002 | *** |
| | | 7,475μm | Negative Control vs. *Ano10 CRISPR* | 0,0049 | ** |
| | | | negative control vs. *Ano10 CRISPR; Ciinte.Brac>Ano10* | 0,5469 | ns |
| | | | *Ano10 CRISPR* vs.*Ano10 CRISPR; Ciinte.Brac>Ano10* | 0,0001 | *** |
| | | 7,6375μm | Negative Control vs. *Ano10 CRISPR* | 0,0023 | ** |
| | | | negative control vs. *Ano10 CRISPR; Ciinte.Brac>Ano10* | 0,6866 | ns |
| | | | *Ano10 CRISPR* vs.*Ano10 CRISPR; Ciinte.Brac>Ano10* | 0,0001 | *** |
| | | 7,8μm | Negative Control vs. *Ano10 CRISPR* | 0,0014 | ** |
| | | | negative control vs. *Ano10 CRISPR; Ciinte.Brac>Ano10* | 0,7634 | ns |
| | | | *Ano10 CRISPR* vs.*Ano10 CRISPR; Ciinte.Brac>Ano10* | 0,0001 | *** |
| | | 7,9625μm | Negative Control vs. *Ano10 CRISPR* | 0,0006 | *** |
| | | | negative control vs. *Ano10 CRISPR; Ciinte.Brac>Ano10* | 0,845 | ns |
| | | | *Ano10 CRISPR* vs.*Ano10 CRISPR; Ciinte.Brac>Ano10* | <0,0001 | **** |
| | | 8,125μm | Negative Control vs. *Ano10 CRISPR* | 0,0003 | *** |
| | | | negative control vs. *Ano10 CRISPR; Ciinte.Brac>Ano10* | 0,948 | ns |
| | | | *Ano10 CRISPR* vs.*Ano10 CRISPR; Ciinte.Brac>Ano10* | <0,0001 | **** |
| | | 8,2875μm | Negative Control vs. *Ano10 CRISPR* | 0,0005 | *** |
| | | | negative control vs. *Ano10 CRISPR; Ciinte.Brac>Ano10* | >0,9999 | ns |
| | | | *Ano10 CRISPR* vs.*Ano10 CRISPR; Ciinte.Brac>Ano10* | 0,0005 | *** |
| | | 8,45μm | Negative Control vs. *Ano10 CRISPR* | 0,0012 | ** |
| | | | negative control vs. *Ano10 CRISPR; Ciinte.Brac>Ano10* | 0,9928 | ns |
| | | | *Ano10 CRISPR* vs.*Ano10 CRISPR; Ciinte.Brac>Ano10* | 0,0018 | ** |

| | | | | | |
|---|---|---|---|---|---|
| | | 8,6125μm | Negative Control vs. *Ano10 CRISPR* | 0,0044 | ** |
| | | | negative control vs. *Ano10 CRISPR; Ciinte.Brac>Ano10* | 0,9943 | ns |
| | | | *Ano10 CRISPR* vs.*Ano10 CRISPR; Ciinte.Brac>Ano10* | 0,0055 | ** |
| | | 8,775μm | Negative Control vs. *Ano10 CRISPR* | 0,0219 | * |
| | | | negative control vs. *Ano10 CRISPR; Ciinte.Brac>Ano10* | 0,9997 | ns |
| | | | *Ano10 CRISPR* vs.*Ano10 CRISPR; Ciinte.Brac>Ano10* | 0,017 | * |
| | | 8,9375μm | Negative Control vs. *Ano10 CRISPR* | 0,0499 | * |
| | | | negative control vs. *Ano10 CRISPR; Ciinte.Brac>Ano10* | 0,9989 | ns |
| | | | *Ano10 CRISPR* vs.*Ano10 CRISPR; Ciinte.Brac>Ano10* | 0,0323 | * |
| | | 11,05μm | Negative Control vs. *Ano10 CRISPR* | 0,28 | ns |
| | | | negative control vs. *Ano10 CRISPR; Ciinte.Brac>Ano10* | 0,5931 | ns |
| | | | *Ano10 CRISPR* vs.*Ano10 CRISPR; Ciinte.Brac>Ano10* | 0,0192 | * |
| | | 11,2125μm | Negative Control vs. *Ano10 CRISPR* | 0,0942 | ns |
| | | | negative control vs. *Ano10 CRISPR; Ciinte.Brac>Ano10* | 0,5202 | ns |
| | | | *Ano10 CRISPR* vs.*Ano10 CRISPR; Ciinte.Brac>Ano10* | 0,0019 | ** |
| | | 11,375μm | Negative Control vs. *Ano10 CRISPR* | 0,0468 | * |
| | | | negative control vs. *Ano10 CRISPR; Ciinte.Brac>Ano10* | 0,4762 | ns |
| | | | *Ano10 CRISPR* vs.*Ano10 CRISPR; Ciinte.Brac>Ano10* | 0,0008 | *** |
| | | 11,5375μm | Negative Control vs. *Ano10 CRISPR* | 0,0273 | * |
| | | | negative control vs. *Ano10 CRISPR; Ciinte.Brac>Ano10* | 0,4629 | ns |
| | | | *Ano10 CRISPR* vs.*Ano10 CRISPR; Ciinte.Brac>Ano10* | 0,0004 | *** |
| | | 11,7μm | Negative Control vs. *Ano10 CRISPR* | 0,0135 | * |
| | | | negative control vs. *Ano10 CRISPR; Ciinte.Brac>Ano10* | 0,3986 | ns |
| | | | *Ano10 CRISPR* vs.*Ano10 CRISPR; Ciinte.Brac>Ano10* | <0,0001 | **** |
| | | 11,8625μm | Negative Control vs. *Ano10 CRISPR* | 0,003 | ** |
| | | | negative control vs. *Ano10 CRISPR; Ciinte.Brac>Ano10* | 0,3651 | ns |
| | | | *Ano10 CRISPR* vs.*Ano10 CRISPR; Ciinte.Brac>Ano10* | <0,0001 | **** |
| | | 12,025μm | Negative Control vs. *Ano10 CRISPR* | 0,0015 | ** |
| | | | negative control vs. *Ano10 CRISPR; Ciinte.Brac>Ano10* | 0,3287 | ns |
| | | | *Ano10 CRISPR* vs.*Ano10 CRISPR; Ciinte.Brac>Ano10* | <0,0001 | **** |
| | | 12,1875μm | Negative Control vs. *Ano10 CRISPR* | 0,0008 | *** |
| | | | negative control vs. *Ano10 CRISPR; Ciinte.Brac>Ano10* | 0,2654 | ns |
| | | | *Ano10 CRISPR* vs.*Ano10 CRISPR; Ciinte.Brac>Ano10* | <0,0001 | **** |
| | | 12,35μm | Negative Control vs. *Ano10 CRISPR* | 0,0007 | *** |
| | | | negative control vs. *Ano10 CRISPR; Ciinte.Brac>Ano10* | 0,2687 | ns |

| | | | | | |
|---|---|---|---|---|---|
| | | | *Ano10 CRISPR* vs.*Ano10 CRISPR; Ciinte.Brac>Ano10* | <0,0001 | **** |
| | | 12,5125µm | Negative Control vs. *Ano10 CRISPR* | 0,001 | *** |
| | | | negative control vs. *Ano10 CRISPR; Ciinte.Brac>Ano10* | 0,2874 | ns |
| | | | *Ano10 CRISPR* vs.*Ano10 CRISPR; Ciinte.Brac>Ano10* | <0,0001 | **** |
| | | 12,675µm | Negative Control vs. *Ano10 CRISPR* | 0,0029 | ** |
| | | | negative control vs. *Ano10 CRISPR; Ciinte.Brac>Ano10* | 0,255 | ns |
| | | | *Ano10 CRISPR* vs.*Ano10 CRISPR; Ciinte.Brac>Ano10* | <0,0001 | **** |
| | | 12,8375µm | Negative Control vs. *Ano10 CRISPR* | 0,0039 | ** |
| | | | negative control vs. *Ano10 CRISPR; Ciinte.Brac>Ano10* | 0,3063 | ns |
| | | | *Ano10 CRISPR* vs.*Ano10 CRISPR; Ciinte.Brac>Ano10* | <0,0001 | **** |
| | | 13µm | Negative Control vs. *Ano10 CRISPR* | 0,0042 | ** |
| | | | negative control vs. *Ano10 CRISPR; Ciinte.Brac>Ano10* | 0,4007 | ns |
| | | | *Ano10 CRISPR* vs.*Ano10 CRISPR; Ciinte.Brac>Ano10* | <0,0001 | **** |
| | | 13,1625µm | Negative Control vs. *Ano10 CRISPR* | 0,0087 | ** |
| | | | negative control vs. *Ano10 CRISPR; Ciinte.Brac>Ano10* | 0,5792 | ns |
| | | | *Ano10 CRISPR* vs.*Ano10 CRISPR; Ciinte.Brac>Ano10* | <0,0001 | **** |
| | | 13,325µm | Negative Control vs. *Ano10 CRISPR* | 0,0151 | * |
| | | | negative control vs. *Ano10 CRISPR; Ciinte.Brac>Ano10* | 0,7655 | ns |
| | | | *Ano10 CRISPR* vs.*Ano10 CRISPR; Ciinte.Brac>Ano10* | 0,0003 | *** |
| | | 13,4875µm | Negative Control vs. *Ano10 CRISPR* | 0,0264 | * |
| | | | negative control vs. *Ano10 CRISPR; Ciinte.Brac>Ano10* | 0,9217 | ns |
| | | | *Ano10 CRISPR* vs.*Ano10 CRISPR; Ciinte.Brac>Ano10* | 0,0027 | ** |
| | | 13,65µm | Negative Control vs. *Ano10 CRISPR* | 0,0559 | ns |
| | | | negative control vs. *Ano10 CRISPR; Ciinte.Brac>Ano10* | 0,9967 | ns |
| | | | *Ano10 CRISPR* vs.*Ano10 CRISPR; Ciinte.Brac>Ano10* | 0,0215 | * |
| Fig. 3Y | Kruskal-Wallis test | | | <0,0001 | **** |
| | | | Negative Control vs. *Ano10 CRISPR* | <0,0001 | **** |
| | | | negative control vs. *Ano10 CRISPR; Ciinte.Brac>Ano10* | 0,0046 | ** |
| | | | *Ano10 CRISPR* vs.*Ano10 CRISPR; Ciinte.Brac>Ano10* | <0,0001 | **** |

Supplementary Table 4 Statistical analysis of data corresponding to panels in Figure 4.

| Figure & Panel | Test | Data point | Comparison | P value | P value summary |
|---|---|---|---|---|---|
| Fig. 4D | Mann Whitney | Amplitude CE | Negative Control vs. *Ano10 CRISPR* | 0,0268 | * |

| | | | | | |
|---|---|---|---|---|---|
| Fig. 4E | Mann Whitney | Rising slope CE | Negative Control vs. *Ano10 CRISPR* | 0,9475 | ns |
| Fig. 4F | Mann Whitney | Falling slope CE | Negative Control vs. *Ano10 CRISPR* | 0,0019 | ** |
| Fig. 4G | Mann Whitney | Duration CE | Negative Control vs. *Ano10 CRISPR* | 0,3303 | ns |
| Fig. 4J | Mann Whitney | Amplitude tubulogenesis | Negative Control vs. *Ano10 CRISPR* | <0,0001 | **** |
| Fig. 4K | Mann Whitney | Rising slope tubulogenesis | Negative Control vs. *Ano10 CRISPR* | <0,0001 | **** |
| Fig. 4L | Mann Whitney | Falling slope tubulogenesis | Negative Control vs. *Ano10 CRISPR* | <0,0001 | **** |
| Fig. 4M | Mann Whitney | Duration tubulogenesis | Negative Control vs. *Ano10 CRISPR* | <0,0001 | **** |
| Fig. 4N | Kruskal-Wallis test | | *All groups* | <0,0001 | **** |
| | Dunn's multiple comparisons | | Number of families 1<br>Number of comparisons per family 15 | | |
| | | | Pre Negative Control vs. Post Negative Control | <0,0001 | **** |
| | | | Pre Negative Control vs. Pre Ano10 CRISPR | >0,9999 | ns |
| | | | Pre Negative Control vs. Post Ano10 CRISPR | <0,0001 | **** |
| | | | Pre Negative Control vs. Pre Ano10 CRISPR; | 0,8814 | ns |
| | | | Pre Negative Control vs. Post Ano10 CRISPR; | <0,0001 | **** |
| | | | Post Negative Control vs. Pre Ano10 CRISPR | <0,0001 | **** |
| | | | Post Negative Control vs. Post Ano10 CRISPR | 0,0002 | *** |
| | | | Post Negative Control vs. Pre Ano10 CRISPR; | <0,0001 | **** |
| | | | Post Negative Control vs. Post Ano10 CRISPR; | >0,9999 | ns |
| | | | Pre Ano10 CRISPR vs. Post Ano10 CRISPR | <0,0001 | **** |
| | | | Pre Ano10 CRISPR vs. Pre Ano10 CRISPR; | 0,8154 | ns |
| | | | Pre Ano10 CRISPR vs. Post Ano10 CRISPR; | <0,0001 | **** |
| | | | Post Ano10 CRISPR vs. Pre Ano10 CRISPR; | 0,0002 | *** |
| | | | Post Ano10 CRISPR vs. Post Ano10 CRISPR; | 0,0013 | ** |
| | | | Pre Ano10 CRISPR; Ciinte.Brac>Ano10 vs. Post | <0,0001 | **** |

| | | | | | |
|---|---|---|---|---|---|
| Fig. 4O | Kruskal-Wallis test | | *All groups* | <0,0001 | **** |
| | Dunn's multiple comparisons test | | Number of families 1<br>Number of comparisons per family 15 | | |
| | | | Pre Negative Control vs. Post Negative Control | <0,0001 | **** |
| | | | Pre Negative Control vs. Pre Ano10 CRISPR | >0,9999 | ns |
| | | | Pre Negative Control vs. Post Ano10 CRISPR | <0,0001 | **** |
| | | | Pre Negative Control vs. Pre Ano10 CRISPR; | >0,9999 | ns |
| | | | Pre Negative Control vs. Post Ano10 CRISPR; | <0,0001 | **** |
| | | | Post Negative Control vs. Pre Ano10 CRISPR | <0,0001 | **** |
| | | | Post Negative Control vs. Post Ano10 CRISPR | <0,0001 | **** |
| | | | Post Negative Control vs. Pre Ano10 CRISPR; | <0,0001 | **** |
| | | | Post Negative Control vs. Post Ano10 CRISPR; | >0,9999 | ns |
| | | | Pre Ano10 CRISPR vs. Post Ano10 CRISPR | <0,0001 | **** |
| | | | Pre Ano10 CRISPR vs. Pre Ano10 CRISPR; | >0,9999 | ns |
| | | | Pre Ano10 CRISPR vs. Post Ano10 CRISPR; | <0,0001 | **** |
| | | | Post Ano10 CRISPR vs. Pre Ano10 CRISPR; | 0,0017 | ** |
| | | | Post Ano10 CRISPR vs. Post Ano10 CRISPR; | 0,0112 | * |
| | | | Pre Ano10 CRISPR; Ciinte.Brac>Ano10 vs. Post | <0,0001 | **** |

Supplementary Table 5 Statistical analysis of data corresponding to panels in Figure 5.

| Figure & Panel | Test | Data point | Comparison | P value | P value summary |
|---|---|---|---|---|---|
| Fig. 5G | Mann Whitney | HEK293T cells patch clamp(Cii.Ano10) -140 mV | Negative control vs Cii.Ano10 | 0.001964 | ** |
| | Mann Whitney | HEK293T cells patch clamp(Cii.Ano10) 100 mV | Negative control vs Cii.Ano10 | 0.010019 | * |
| | Mann Whitney | HEK293T cells patch clamp(Cii.Ano10-CAAX) -140 mV | Negative control vs Cii.Ano10-CAAX | 0.241607 | ns |
| | Mann Whitney | HEK293T cells patch clamp(Cii.Ano10-CAAX) 100 mV | Negative control vs Cii.Ano10-CAAX | 0.001216 | ** |

Supplementary Table 6 Statistical analysis of data corresponding to panels in Supplemental Figure 4.

| Figure & Panel | Test | Data point | Comparison | P value | P value summary |
|---|---|---|---|---|---|
| Sup. Fig. 4I | Kruskal-Wallis test | | All groups | <0,0001 | **** |
| | Dunn's multiple comparisons test | | Number of families<br>Number of comparisons per family | | 1<br>3 |
| | | | Negative Control vs. *Ano10 CRISPR* | 0,0002 | *** |
| | | | Negative control vs. *Ano10 CRISPR; Ciinte.Brac>Ano10* | >0,9999 | ns |
| | | | *Ano10 CRISPR* vs.*Ano10 CRISPR; Ciinte.Brac>Ano10* | <0,0001 | **** |
| Sup. Fig. 4M | Mann Whitney test | | Negative Control vs. *Ano10 CRISPR* | 0,8921 | ns |

Supplementary Table 7 Statistical analysis of data corresponding to panels in Supplemental Figure 6.

| Figure & | Test | Data point | Comparison | P value | P value summary |
|---|---|---|---|---|---|
| Sup. Fig. 6E | Kruskal-Wallis test | | All groups | 0,7061 | ns |
| | Dunn's multiple comparisons test | | Number of families 1<br>Number of comparisons per family 6 | | |
| | | | DMSO vs T16A(inh)-A01 | >0,9999 | ns |
| | | | DMSO vs NPPB | >0,9999 | ns |
| | | | DMSO vs NS 3728 | >0,9999 | ns |
| | | | T16A(inh)-A01 vs NPPB | >0,9999 | ns |
| | | | T16A(inh)-A01 vs NS 3728 | >0,9999 | ns |
| | | | NPPB vs NS 3728 | >0,9999 | ns |
| Sup. Fig. [illegible] | Kruskal-Wallis test | | All groups | 0,0006 | *** |
| | Dunn's multiple comparisons test | | Number of families 1<br>Number of comparisons per family 6 | | |
| | | | DMSO vs T16A(inh)-A01 | 0,0024 | ** |
| | | | DMSO vs NPPB | 0,0126 | * |
| | | | DMSO vs NS 3728 | 0,0024 | ** |
| | | | T16A(inh)-A01 vs NPPB | >0,9999 | ns |
| | | | T16A(inh)-A01 vs NS 3728 | >0,9999 | ns |
| | | | NPPB vs NS 3728 | >0,9999 | ns |
| Sup. Fig. 6G | Kruskal-Wallis test | | All groups | <0,0001 | *** |

| | | | | | |
|---|---|---|---|---|---|
| | Dunn's multiple comparisons test | | Number of families 1<br>Number of comparisons per family 6 | | |
| | | | DMSO vs T16A(inh)-A01 | <0,0001 | **** |
| | | | DMSO vs NPPB | <0,0001 | **** |
| | | | DMSO vs NS 3728 | <0,0001 | **** |
| | | | T16A(inh)-A01 vs NPPB | >0,9999 | ns |
| | | | T16A(inh)-A01 vs NS 3728 | >0,9999 | ns |
| | | | NPPB vs NS 3728 | >0,9999 | ns |
| Sup. Fig. 6H | Mixed-effects model | | Cross-sectional area | <0,0001 | **** |
| | | | Time | <0,0001 | **** |
| | | | Cross-sectional area x Time | <0,0001 | **** |
| | Within each row, compare columns (simple effects within rows) | | Number of families 15<br>Number of comparisons per family 6 | | |
| | | 0 min | DMSO vs NPPB | 0,2849 | ns |
| | | | DMSO vs NS 3728 | 0,4112 | ns |
| | | | DMSO vs T16A(inh)-A01 | 0,3925 | ns |
| | | | NPPB vs NS 3728 | 0,9105 | ns |
| | | | T16A(inh)-A01 vs NPPB | 0,9919 | ns |
| | | | T16A(inh)-A01 vs NS 3728 | 0,2849 | ns |
| | | 30 min | DMSO vs NPPB | 0,1746 | ns |
| | | | DMSO vs NS 3728 | 0,9943 | ns |
| | | | DMSO vs T16A(inh)-A01 | 0,9986 | ns |
| | | | NPPB vs NS 3728 | 0,6262 | ns |
| | | | T16A(inh)-A01 vs NPPB | 0,4497 | ns |
| | | | T16A(inh)-A01 vs NS 3728 | 0,9870 | ns |
| | | 60 min | DMSO vs NPPB | 0,0176 | * |
| | | | DMSO vs NS 3728 | 0,5287 | ns |
| | | | DMSO vs T16A(inh)-A01 | 0,7378 | ns |
| | | | NPPB vs NS 3728 | 0,8276 | ns |
| | | | T16A(inh)-A01 vs NPPB | 0,7094 | ns |
| | | | T16A(inh)-A01 vs NS 3728 | 0,9959 | ns |
| | | 90 min | DMSO vs NPPB | 0,0005 | *** |
| | | | DMSO vs NS 3728 | 0,0339 | * |

| | | | | | |
|---|---|---|---|---|---|
| | | | DMSO vs T16A(inh)-A01 | 0,1890 | ns |
| | | | NPPB vs NS 3728 | 0,9790 | ns |
| | | | T16A(inh)-A01 vs NPPB | 0,7247 | ns |
| | | | T16A(inh)-A01 vs NS 3728 | 0,9419 | ns |
| | | 120 min | DMSO vs NPPB | <0,0001 | **** |
| | | | DMSO vs NS 3728 | 0,0016 | ** |
| | | | DMSO vs T16A(inh)-A01 | 0,0364 | * |
| | | | NPPB vs NS 3728 | 0,9987 | ns |
| | | | T16A(inh)-A01 vs NPPB | 0,7917 | ns |
| | | | T16A(inh)-A01 vs NS 3728 | 0,9040 | ns |
| | | 150 min | DMSO vs NPPB | <0,0001 | **** |
| | | | DMSO vs NS 3728 | 0,0002 | *** |
| | | | DMSO vs T16A(inh)-A01 | 0,0147 | * |
| | | | NPPB vs NS 3728 | 0,9738 | ns |
| | | | T16A(inh)-A01 vs NPPB | 0,7905 | ns |
| | | | T16A(inh)-A01 vs NS 3728 | 0,9536 | ns |
| | | 180 min | DMSO vs NPPB | <0,0001 | **** |
| | | | DMSO vs NS 3728 | <0,0001 | **** |
| | | | DMSO vs T16A(inh)-A01 | 0,0041 | ** |
| | | | NPPB vs NS 3728 | 0,9838 | ns |
| | | | T16A(inh)-A01 vs NPPB | 0,8971 | ns |
| | | | T16A(inh)-A01 vs NS 3728 | 0,9824 | ns |
| | | 210 min | DMSO vs NPPB | <0,0001 | **** |
| | | | DMSO vs NS 3728 | <0,0001 | **** |
| | | | DMSO vs T16A(inh)-A01 | 0,0006 | *** |
| | | | NPPB vs NS 3728 | 0,9878 | ns |
| | | | T16A(inh)-A01 vs NPPB | 0,9991 | ns |
| | | | T16A(inh)-A01 vs NS 3728 | 0,9994 | ns |
| | | 240 min | DMSO vs NPPB | <0,0001 | **** |
| | | | DMSO vs NS 3728 | <0,0001 | **** |
| | | | DMSO vs T16A(inh)-A01 | 0,0003 | *** |
| | | | NPPB vs NS 3728 | 0,8698 | ns |
| | | | T16A(inh)-A01 vs NPPB | 0,9963 | ns |
| | | | T16A(inh)-A01 vs NS 3728 | 0,9862 | ns |
| | | 270 min | DMSO vs NPPB | <0,0001 | **** |

| | | | | | |
|---|---|---|---|---|---|
| | | | DMSO vs NS 3728 | <0,0001 | **** |
| | | | DMSO vs T16A(inh)-A01 | <0,0001 | **** |
| | | | NPPB vs NS 3728 | 0,7752 | ns |
| | | | T16A(inh)-A01 vs NPPB | 0,9917 | ns |
| | | | T16A(inh)-A01 vs NS 3728 | 0,9726 | ns |
| | | 300 min | DMSO vs NPPB | <0,0001 | **** |
| | | | DMSO vs NS 3728 | <0,0001 | **** |
| | | | DMSO vs T16A(inh)-A01 | <0,0001 | **** |
| | | | NPPB vs NS 3728 | 0,8250 | ns |
| | | | T16A(inh)-A01 vs NPPB | 0,9968 | ns |
| | | | T16A(inh)-A01 vs NS 3728 | 0,8411 | ns |
| | | 330 min | DMSO vs NPPB | <0,0001 | **** |
| | | | DMSO vs NS 3728 | <0,0001 | **** |
| | | | DMSO vs T16A(inh)-A01 | <0,0001 | **** |
| | | | NPPB vs NS 3728 | 0,8999 | ns |
| | | | T16A(inh)-A01 vs NPPB | 0,9598 | ns |
| | | | T16A(inh)-A01 vs NS 3728 | 0,7697 | ns |
| | | 360 min | DMSO vs NPPB | <0,0001 | **** |
| | | | DMSO vs NS 3728 | 0,0001 | *** |
| | | | DMSO vs T16A(inh)-A01 | <0,0001 | **** |
| | | | NPPB vs NS 3728 | 0,9570 | ns |
| | | | T16A(inh)-A01 vs NPPB | 0,8282 | ns |
| | | | T16A(inh)-A01 vs NS 3728 | 0,6720 | ns |
| | | 390 min | DMSO vs NPPB | <0,0001 | **** |
| | | | DMSO vs NS 3728 | 0,0002 | *** |
| | | | DMSO vs T16A(inh)-A01 | <0,0001 | **** |
| | | | NPPB vs NS 3728 | 0,9431 | ns |
| | | | T16A(inh)-A01 vs NPPB | 0,6841 | ns |
| | | | T16A(inh)-A01 vs NS 3728 | 0,5279 | ns |
| | | 420 min | DMSO vs NPPB | <0,0001 | **** |
| | | | DMSO vs NS 3728 | 0,0005 | *** |
| | | | DMSO vs T16A(inh)-A01 | <0,0001 | **** |
| | | | NPPB vs NS 3728 | 0,8563 | ns |
| | | | T16A(inh)-A01 vs NPPB | 0,7593 | ns |
| | | | T16A(inh)-A01 vs NS 3728 | 0,4911 | ns |

Supplementary Table 8 Primers used in this study

| ID of amplification product | Fw/Rv primer | Rv Primer |
|---|---|---|
| *Cii.Brachyury promoter gateway* | | ggggacaactttgtatagaaaagtt<br>gCTGAACAAGCCATGTGCCGG |
| | | ggggactgcttttttgtacaaactt<br>gTCTGCCTCCAAATCACACTCG |
| Cii.Carbonic anhydrase 2 promoter KH.C1.423; KY21.Chr1.1715 | AscI-Fw | taaggcgcgccAATAAAGGGATGCTCGCCCCATG |
| | NotI-Rv | aatgcggccgcGCTTCTATGCCTGTTTATAAATAC |
| Cii.Carbonic anhydrase 2 promoterKH.C1.423; KY21.Chr1.1715 | | ggggacaactttgtatagaaaagtt<br>gCACGGTCTGTTGTTCGGTAACAG |
| | | ggggactgcttttttgtacaaactt<br>gTAACACGATGTGCTTCGATTGC |
| 2nd position CD4GFP | Fw | ggggacaagtttgtacaaaaaagcaggc<br>taaccATGAATCCCAAGAGCGAAGTCC |
| | Rv | ggggaccactttgtacaagaaagctggg<br>tTTAGAGGGCAACTTCATTTTCATAG |
| 2nd position tdTomato-Lifeact | Fw | ggggacaagtttgtacaaaaaagcaggc<br>taaccATGGGCGTGGCCGACTTGAT |
| | Rv | ggggaccactttgtacaagaaagctggg<br>tTTACTTGTACAGCTCGTCCATGCC |
| EB3-mNeonGreen | Fw | ggggacaagtttgtacaaaaaagcaggc<br>taaccATGGCCGTCAATGTGTACTCCAC |
| | Rv | ggggaccactttgtacaagaaagctggg<br>tTTACTTGTACAGCTCGTCCATGCC |
| nls::Cas9::nls | Fw | ggggacaagtttgtacaaaaaagcaggc<br>taaccATGGCTAGCCCCAAAAAGAAGAGG |
| | Rv | ggggaccactttgtacaagaaagctggg<br>tTCATTAACCTACCTTGCGCTTTTTCTTG |
| Crispr/Cas9 control gRNA | | agatGCTTTGCTACGATCTACATT |
| | | aaacAATGTAGATCGTAGCAAAGC |
| Crispr/Cas9 *Ano10* gRNA | | agatGTTTAAATGAGGTAACGCAG |
| | | aaacCTGCGTTACCTCATTTAAAC |
| *Cii.Ano10* for pEGFP-N1 | XhoI | taactcgagATGAGTGAAACTGAAAAGTTGCATG |
| | ApaI | atagggccccGAACGGCGTTGTTTCTTTTTCAT |
| Rat *Ano6* for pEGFP-N1 | XhoI | taactcgagATGCAGATGATGACTAGGAACG |
| | ApaI | atagggccccTTCGAGTTTCGGCCTCACG |
| *Cii.Ano6* for pEGFP-N1 | XhoI | aggctcgagATGAACACGTTTCGAGACGGAA |
| | ApaI | aatgtcgacCTACTTGTCGTCATCGTCTTTGTAGTCCATTTGGGTATTCTCTATGTCTTTG |
| Cii.Ano10 for in situ probe | Fw | CTGAGACCACAGAATACACCAG |
| | Rv | CAAGACATGAATCACTGACAGAAC |

Supplementary Table 9 Percentage of animals expressing *Cii.Cahy>GFP* or *Cii.Brachyury>GFP* at certain developmental stages.

| Stage (according to Hotta et al.) | *Cii.Cah>GFP* (+)signal/out 360 embryos (3 trials) | *Cii.Cah>GFP* (% expressing) | *Cii.Brachyury>GFP* (+)signal/out 210 embryos (3 trials) | *Cii.Brachyury>GFP (% expressing)* |
|---|---|---|---|---|
| 14 | 11 | 3.06 | 143 | 68.10 |
| 16 | 19 | 5.28 | 171 | 81.43 |
| 18 | 78 | 21.67 | 188 | 89.52 |
| 20 | 170 | 47.22 | 195 | 92.86 |
| 22 | 249 | 69.17 | 197 | 93.80 |
| 24 | 261 | 72.50 | 197 | 93.80 |
| 26 | 287 | 79.72 | 197 | 93.80 |

**Supplementary Movie 1**

Confocal time lapse imaging of *Ciiinte.Brac>hCD4-GFP (green); Ciiinte.Caveolin 1-mCherry* (magenda) covering the period of tubulogenesis in a negative control embryo.

**Supplementary Movie 2**

Confocal time lapse imaging of *Ciiinte.Brac>hCD4-GFP (green); Ciiinte.Caveolin 1-mCherry* (magenda) covering the period of tubulogenesis in an Ano10$^{CRISPR}$ embryo.

**Supplementary Movie 3**

Confocal time lapse imaging of *Ciiinte.Brac>hCD4-GFP (green); Ciiinte.Caveolin 1-mCherry* (magenda) covering the period of tubulogenesis in an Ano10$^{CRISPR}$ ; *Ciiinte.Brac>Ano10cDNA rescue* embryo.

**Supplementary Movie 4**

First example movie of *Ciiinte.Brac>GCaMP6s* showing $Ca^{2+}$ activity during notochord cell intercalation.

**Supplementary Movie 5**

Second example movie of *Ciiinte.Brac>GCaMP6s* showing $Ca^{2+}$ activity during notochord cell intercalation.

**Supplementary Movie 6**

Example movie of *Ciiinte.Brac>GCaMP6s* showing $Ca^{2+}$ activity during notochord lumen extension.

**Supplementary Movie 7**

Example movie of *Ciiinte.Brac>GCaMP6s* showing $Ca^{2+}$ activity during notochord lumen tilting.

**Supplementary Movie 8**

Example movie of *Ciiinte.Brac>GCaMP6s* showing $Ca^{2+}$ activity during notochord cell bi-directional cloning.

**Supplementary Movie 9**

Confocal time lapse imaging of *Ciiinte.Brac>hCD4-GFP (green); Ciiinte.Caveolin 1-mCherry* (magenda) covering the period of tubulogenesis in DMSO incubated control embryo.

**Supplementary Movie 10**

Confocal time lapse imaging of *Ciiinte.Brac>hCD4-GFP (green); Ciiinte.Caveolin 1-mCherry* (magenda) covering the period of tubulogenesis in a T16A inhibitor (100μM) incubated embryo.

**Supplementary Movie 11**

Confocal time lapse imaging of *Ciiinte.Brac>hCD4-GFP (green); Ciiinte.Caveolin 1-mCherry* (magenda) covering the period of tubulogenesis in a NPPB (100μM) inhibitor incubated embryo.

**Supplementary Movie 12**

Confocal time lapse imaging of *Ciiinte.Brac>hCD4-GFP (green); Ciiinte.Caveolin 1-mCherry* (magenda) covering the period of tubulogenesis in a NS3728 (100μM) inhibitor incubated embryo.
